# A Scalable, Open-Source Implementation of a Large-Scale Mechanistic Model for Single Cell Proliferation and Death Signaling

**DOI:** 10.1101/2020.11.09.373407

**Authors:** Cemal Erdem, Arnab Mutsuddy, Ethan M. Bensman, William B. Dodd, Michael M. Saint-Antoine, Mehdi Bouhaddou, Robert C. Blake, Sean M. Gross, Laura M. Heiser, F. Alex Feltus, Marc R. Birtwistle

**Affiliations:** Department of Chemical & Biomolecular Engineering, Clemson University, Clemson, SC; Computer Science, School of Computing, Clemson University, Clemson, SC; Center for Bioinformatics and Computational Biology, University of Delaware, Newark, DE; Department of Cellular and Molecular Pharmacology, University of California San Francisco, San Francisco, CA; Center for Applied Scientific Computing, Lawrence Livermore National Laboratory, Livermore, CA; Department of Biomedical Engineering, Oregon Health & Science University, Portland, OR; Department of Genetics and Biochemistry, Clemson University, Clemson, SC; Biomedical Data Science and Informatics Program, Clemson University, Clemson, SC; Center for Human Genetics, Clemson University, Clemson, SC; Department of Bioengineering, Clemson University, Clemson, SC

## Abstract

Mechanistic models of how single cells respond to different perturbagens can help integrate disparate big data sets or predict response to varied drug combinations. However, the construction and simulation of such models have proved challenging. Our lab previously constructed one of the largest mechanistic models for single mammalian cell regulation of proliferation and death (774 species, 141 genes, 8 ligands, 2400 reactions). However, this, as many other large-scale models, was written using licensed software (MATLAB) with intricate programming structure, impeding alteration, expansion, and sharing. Here, we generated a new foundation for this model, which includes a python-based creation and simulation pipeline converting a few structured text files into an SBML-compatible format. This new open-source model (named SPARCED) is high-performance- and cloud-computing compatible and enables the study of virtual cell population responses at the single-cell level. We applied this new model to a subset of the LINCS MCF10A Data Cube, which observed that IFNγ acts as an anti-proliferative factor, but the reasons why were unknown. After expanding the SPARCED model with an IFNγ signaling module (to 950 species, 150 genes, 9 ligands, 2500 reactions), we ran stochastic single-cell simulations for two different putative crosstalk mechanisms and looked at the number of cycling cells in each case. Our model-based analysis suggested, and experiments support that these observations are better explained by IFNγ-induced SOCS1 expression sequestering activated EGF receptors, thereby downregulating AKT activity, as opposed to direct IFNγ-induced upregulation of p21 expression. This work forms a foundation for increased mechanistic model-based data integration on a single-cell level, an important building block for clinically predictive mechanistic models.

## INTRODUCTION

The ever-increasing availability and accumulation of FAIR (1) (findable, accessible, interoperable, and reproducible) and big (omics) datasets requires new computational methods and models to integrate, analyze, and interpret the underlying information (2–4). How can we leverage the totality of available information not only to learn more about biology but also to make predictions, especially those that are clinically relevant? Advances in statistical and machine learning approaches enable (mostly) data-driven exploration and hypothesis generation from big datasets (5–8). Trained on features of the input dataset(s), such models can be used for, as just a few examples, to predict drug responses (9–11) or decide tumor type/stage (12–15). Although transformative, such machine learning and statistical models have shortcomings. Most notably, they often fail to explain predicted outcomes with detailed mechanistic reasoning (16–20) – a major scientific gap and a roadblock to reconciling and integrating such models.

Besides such “black-box” modeling approaches, an alternative and complementary vehicle for data integration are so-called “mechanistic models” (20). Mechanistic models provide an interpretable integration of different data types, because they have explicitly modeled biophysical correlates, while enabling further exploration for underlying logic behind heterogeneous, nonlinear, and often unintuitive relationships across big datasets (21). If mechanistic models are available towards the whole-genome or whole-single-cell scale, one can start to predict complex, multi-network, and emergent cellular behaviors (22, 23), elucidate phenotypic responses to multiple perturbations (24, 25), tailor and train on patient-specific data for personalized, pharmacologic decision making (26, 27), or use them as “data integrators” for data consistency checking (28). However, most published mechanistic models are “small” scale; built for single pathways with a handful of genes, meant to interpret a single dataset (29–38). Such small-scale mechanistic models provided important insights into processes such as yeast response to pheromones (35), *lac* operon regulation in *E. coli* (34), or phenotypic responses to different ligand stimulations (29). However, the limited scope of small-scale models means they inherently will struggle to integrate multiple datasets. Large-scale mechanistic models (23,39–41), on the other hand, can provide a more extensive representation of cellular interactions and are thus well-poised for data integration that complement shortcomings of machine learning approaches.

One of the many ways of mechanistic model construction is the use and modification of existing models by inserting new species or interactions to explain new experimental observations (38,42,43). Model merging, the act of stitching pre-existing models together, is an extension of this method for creating larger models. However, such an approach requires extensive detail checking and harmonizing species/parameter definitions. Often, unfortunately, sufficient annotation is not provided which makes this task harder. Moreover, while most mechanistic models are comprised of ordinary differential equations (ODEs), many large-scale models require multiple sub-modules of different mathematical formalisms. For example, metabolic processes are usually described by steady-state flux-balance models (44, 45), gene expression events are stochastic (46–48), and protein signaling events are represented by a system of ODEs (29,30,38). Thus, sorting out a single platform for different modeling formalisms to create a large-scale model is a daunting task. It is so far only achieved by creating highly custom-structured and custom-coded model-agglomerates that are not well-suited to further alterations or re-use (23, 40). The latter, Bouhaddou2018 pan-cancer model (40), is previously published by our group to study single-cell responses to mitogens and drugs.

A second way of constructing models is to build them bottom-up by writing out every reaction one by one. In this regard, rule-based modeling (RBM) provides an innovative approach (49). RBM software, such as BioNetGen (50, 51), Kappa (52), and PySB (53), enables researchers to write “rules” for repeated reaction events following specific patterns. RBM software then creates the reaction network by propagating the rules from the initial set of species. Although RBM revolutionized large-scale model construction by minimizing manual equation scripting (i.e., writing out every differential equation), some limitations exist. First, it can generate a vast (even infinite) number of reactions from a small set of rules (usually called the curse of combinatorial complexity). This makes interpreting, analyzing, and debugging such models cumbersome, if possible. Tools like NFsim (54) can overcome such problems by simulating events based on the rules rather than a priori generating the entire reaction network. Thus, such software becomes advantageous when a small number of rules create a very large number of reactions, e.g., polymerization, aggregation, or multi-site phosphorylation (55). However, such network-free simulators typically require an explicit representation of every molecule in the system, which dramatically increases the computational cost and renders such methods inefficient for large-scale mechanistic models. Secondly, current RBM implementations dictate that reactions taking place via the same rule have the same rate constant parameter values. Often, allostery or site cooperativity precludes this simplifying assumption, leading to manually writing out every such reaction in the model (or writing one rule for each reaction), which then obviates the advantages of RBM. Finally, with its capability of capturing biological complexity via simple rules, the RBM concept is quite powerful but additional efforts are needed to enable merging of existing non-rule-based models, creating a mixture of different modeling formats (i.e. mixed-grain modeling), and defining different simulation settings (i.e. hybrid modeling = deterministic + stochastic parts).

Regardless of how a large-scale model is constructed, it should have certain properties for FAIRness (findable, accessible, interoperable, and reproducible) and re-useability (56–58). Porubsky et al. (57) recently summarized the *best modeling practices* and reinforced: providing metadata/annotations and model creation steps/files (Practices 1-5), using standard and cross-platform model files (Practice 3), and open-source, license-free, version-controlled, and reproducible model dissemination (Practices 8-9). As the size of the model increases, conforming to modeling standards (e.g. simulation type, simulation speed, software to use, scripting package to use, algorithm to use) gets harder. That is why most of the large-scale (many genes or whole-cell) models are necessarily custom-structured, are composed of multiple submodules, or are lacking sufficient annotations and metadata (e.g. ENSEMBL or HGNC identifiers) (23,39–41). These custom-made models also do not yet follow a single standard format, a key property for easy distribution, re-use, and model merging and expansion with other models. The SBML (Systems Biology Markup Language) format (59, 60) offers a long-established and well-defined way of specifying annotated model structures, with an explicit and structured definition of each element of a mechanistic model (species, reactions, volumes, initial concentrations, parameters, rules, events, equations). SBML is an extensible, machine-readable markup language and not a simple text file. SBML has interfaces and packages in most programming languages (like Python, C++, Perl) and can be imported by most software (Python, MATLAB, COPASI (61), Virtual Cell (62), and another ∼300 packages). However, it is non-trivial to write thousands of reactions in SBML standards, directly or with available GUI-based software. To circumvent this problem, there are efforts to convert other model formats to SBML, like Antimony (63). The Antimony format is defined in simple text format and is human readable and interpretable. Regardless, any constructed mechanistic model, in SBML format or not, must be simulated with reasonable CPU time. Although simulating models on local machines is often done, High Performance (HPC) or Cloud Computing (CC) platforms are suitable for larger tasks such as parameter sensitivity/estimation or multiple single-cell simulations (64–67). Therefore, another milestone for large-scale mechanistic models is inherent HPC/CC compatibility, especially for single-cell simulations and heterogeneous data integration.

Here, we provide a framework for large-scale mechanistic modeling that converts our lab’s previous large-scale pan-cancer model into a format that conveys several crucial properties noted above. First, we define a simple set of structured and annotated input text files that set model specifics: genes, species, reactions, reaction stoichiometry, cellular compartments, transcriptional regulations, input omics data, and parameter values (Fig. 1). These text files enable easy creation or alteration of the model network, without any coding or software usage requirements (but they are easily amenable to such things if desired). We then use Jupyter notebooks (68) to process the input files and to create a human-interpretable Antimony file, which is then converted into an SBML (community gold-standard) model file. We simulate the model using SBML compatible Python packages including AMICI, specifically designed for efficient simulation of large-scale models (67, 69), and our own Python submodule for stochastic gene expression that enables single-cell simulations. We also develop an HPC/CC (Kubernetes) compatible version of the pipeline that enables simulating large number of single cells and/or stimulation conditions. To apply our work, we re-create and extend our previous single mammalian cell mechanistic model of proliferation and death signaling and regulation (40), which we call SPARCED (**S**BML, **P**roliferation, **A**poptosis, **R**eceptor Tyrosine Kinases, **C**ell cycle, **E**xpression, **D**NA damage). The pipeline and model are available on GitHub (github.com/birtwistlelab/SPARCED). Using the newly created model format, we explore different mechanisms to explain the experimental observation that interferon-gamma (IFNγ) inhibits epidermal growth factor (EGF)-induced cell proliferation. Our model analysis suggests, and experiments support the hypothesis that IFNγ inhibits cell proliferation through SOCS1-induction and reduced AKT activity, rather than by IFNγ-induced p21 expression upregulation. This large-scale mechanistic model construction framework and the SPARCED model outlined here is an important step towards creating and testing large-scale mechanistic models as data integration and clinical decision-making tools.

**Fig 1.**
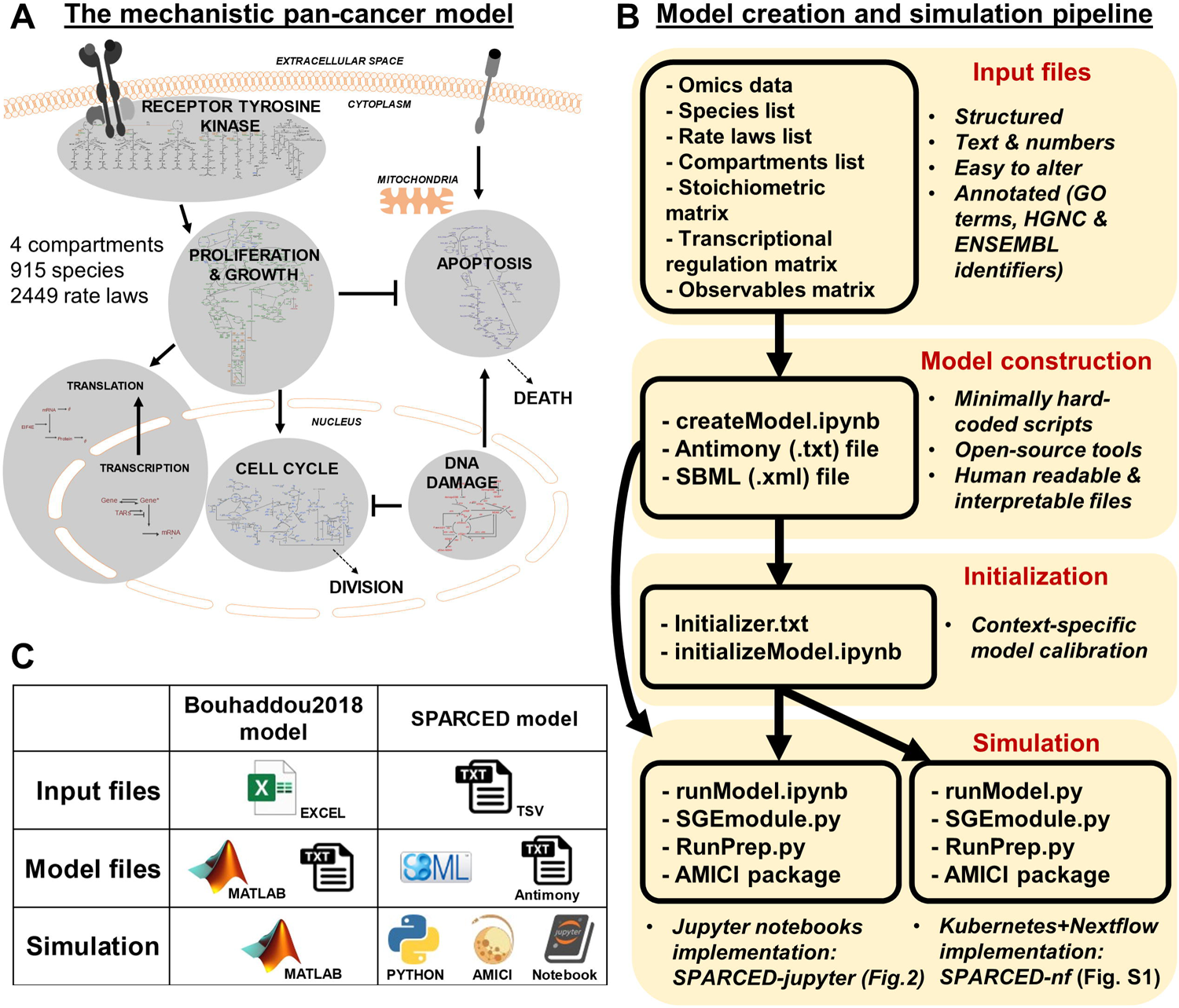
SPARCED is a structured, human interpretable, and easy to modify big mechanistic model. (A) The schematic of the underlying model for SPARCED. Image adapted from (40). (B) The pan-cancer mechanistic model Bouhaddou2018 is re-written in open-source and structured file format. The steps of model construction include input file creation and conversion into an SBML file. The optional initialization step calibrates model parameters for new cellular contexts and phenotypic behaviors. The annotated SBML model file and stochastic module are simulated together at single-cell level locally or by using cloud-computing. The benefits of the new SPARCED model include easy alteration and expansion capabilities through text file editing, human-readable annotated input files, and use of Jupyter notebooks for model creation and simulation. The modeling pipeline introduced here are inline with good practices of re-usable big mechanistic models (57). (C) The Bouhaddou2018 model file types are simplified and converted into open-source platforms.

## RESULTS

### SPARCED Model Construction and Unit Testing

Current large-scale mechanistic models are agglomerates of smaller models and tools, used mainly within the same research lab. Most such models also lack clear and satisfactory annotation and metadata, making them harder to understand and alter (23, 40). The goals of this work were (i) to build tools that help large-scale mechanistic model construction and alteration, that is simple, efficient, open-source, and cloud computing compatible; (ii) to provide a scalable and re-useable big mechanistic model for a single mammalian cell; and (iii) demonstrate the work through application to a biological question.

We first created a set of simple input files and scalable processing scripts for one of the broadest cancer signaling models in the literature (40), called the Bouhaddou2018 model here (Fig. 1A). The input files (Supp. Files 1-7) are simple tab-separated text files (Fig. 1B-C), unlike licensed file formats with a mixture of hard coded information in multiple interconnected scripts commonly used in modeling literature. A Jupyter notebook (Supp. File 8) processes the input files into an Antimony text file (Supp. File 9). The model creation code generates the SPARCED model file in SBML format (Supp. File 10) using the Antimony text file and annotations from the model input files (Supp. Files 2 and 6). When the model construction step is complete and the SBML file is created, it is imported and simulated using a Python package called AMICI (67, 69). For every new cell line model, a pre-calibration step called *Initialization* is employed to tune parameter values. Here, we ensure total protein levels match experimental observations and particular phenotypic criteria are met; for example, we specify that serum and growth factor starved cells on average do not traverse the cell cycle and do not die by apoptosis within 48 hours. The resulting *initialized* parameter values and species concentrations are saved in a new SBML file, and the model is compiled for model testing and other simulations.

The result is what should be a replica of the Bouhaddou2018 model, which we call SPARCED. Like the Bouhaddou2018 model, the initial SPARCED model is based on non-transformed breast epithelial MCF10A cell line data. We annotated all the species in the model with HGNC gene identifiers, providing easier programmatic filtering and curation of species list, while keeping the user defined simpler names for complicated species structures. However, the extent to which the models are congruent was not yet clear, and thus we next set out to examine agreement between the two. We verified that the previous Bouhaddou2018 model simulations are reproducible and match expected experimental observations through the same unit test concept (Table 1) introduced for the original model (40). Each unit test has a dedicated Jupyter notebook on GitHub repository (github.com/birtwistlelab/SPARCED/SPARCED_Brep). We illustrate select unit testing examples below, but all results are presented in supplementary figures (Figs. S2-11).

**Table 1.**
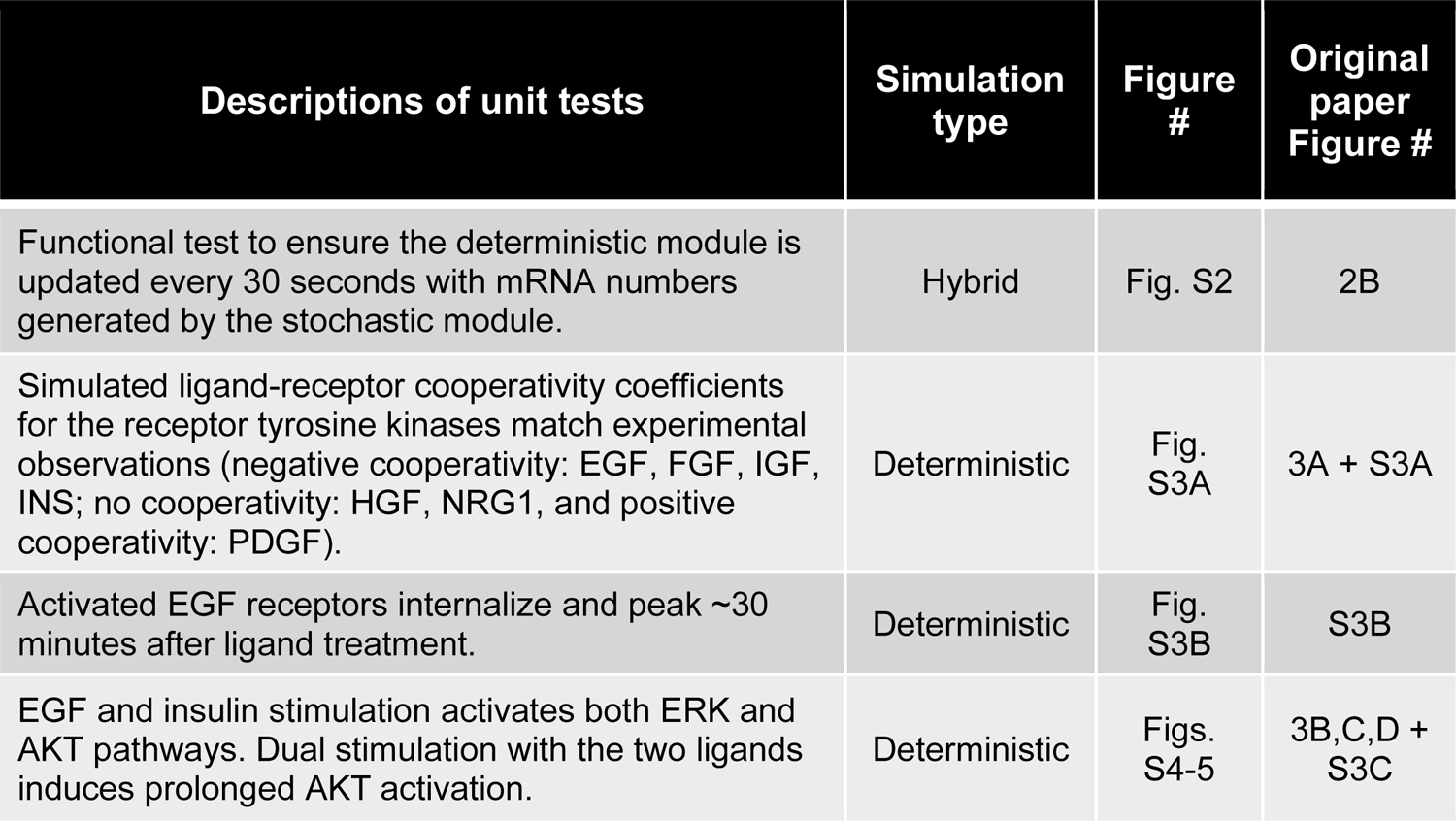
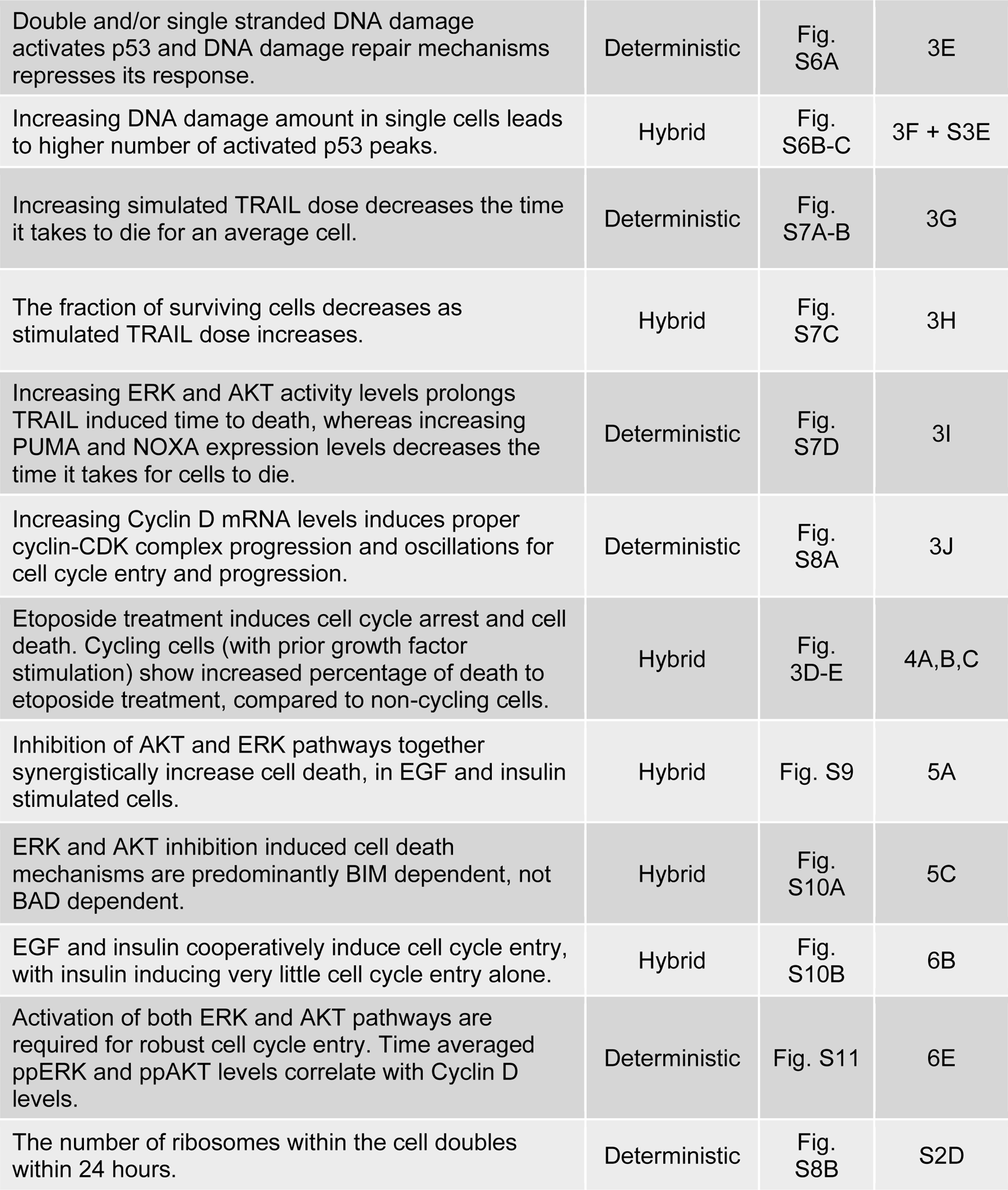
List of SPARCED model unit testing and comparisons to Bouhaddou2018 model. The SPARCED model passed each test depicted below and recapitulated experimental and simulation observations reported by the Bouhaddou2018 model.

| Descriptions of unit tests | Simulation type | Figure # | Original paper Figure # |
| --- | --- | --- | --- |
| Functional test to ensure the deterministic module is updated every 30 seconds with mRNA numbers generated by the stochastic module. | Hybrid | Fig. S2 | 2B |
| Simulated ligand-receptor cooperativity coefficients for the receptor tyrosine kinases match experimental observations (negative cooperativity: EGF, FGF, IGF, INS; no cooperativity: HGF, NRG1, and positive cooperativity: PDGF). | Deterministic | Fig. S3A | 3A + S3A |
| Activated EGF receptors internalize and peak ~30 minutes after ligand treatment. | Deterministic | Fig. S3B | S3B |
| EGF and insulin stimulation activates both ERK and AKT pathways. Dual stimulation with the two ligands induces prolonged AKT activation. | Deterministic | Figs. S4-5 | 3B,C,D + S3C |
| Double and/or single stranded DNA damage activates p53 and DNA damage repair mechanisms represses its response. | Deterministic | Fig. S6A | 3E |
| Increasing DNA damage amount in single cells leads to higher number of activated p53 peaks. | Hybrid | Fig. S6B-C | 3F + S3E |
| Increasing simulated TRAIL dose decreases the time it takes to die for an average cell. | Deterministic | Fig. S7A-B | 3G |
| The fraction of surviving cells decreases as stimulated TRAIL dose increases. | Hybrid | Fig. S7C | 3H |
| Increasing ERK and AKT activity levels prolongs TRAIL induced time to death, whereas increasing PUMA and NOXA expression levels decreases the time it takes for cells to die. | Deterministic | Fig. S7D | 3I |
| Increasing Cyclin D mRNA levels induces proper cyclin-CDK complex progression and oscillations for cell cycle entry and progression. | Deterministic | Fig. S8A | 3J |
| Etoposide treatment induces cell cycle arrest and cell death. Cycling cells (with prior growth factor stimulation) show increased percentage of death to etoposide treatment, compared to non-cycling cells. | Hybrid | Fig. 3D-E | 4A,B,C |
| Inhibition of AKT and ERK pathways together synergistically increase cell death, in EGF and insulin stimulated cells. | Hybrid | Fig. S9 | 5A |
| ERK and AKT inhibition induced cell death mechanisms are predominantly BIM dependent, not BAD dependent. | Hybrid | Fig. S10A | 5C |
| EGF and insulin cooperatively induce cell cycle entry, with insulin inducing very little cell cycle entry alone. | Hybrid | Fig. S10B | 6B |
| Activation of both ERK and AKT pathways are required for robust cell cycle entry. Time averaged ppERK and ppAKT levels correlate with Cyclin D levels. | Deterministic | Fig. S11 | 6E |
| The number of ribosomes within the cell doubles within 24 hours. | Deterministic | Fig. S8B | S2D |

### SPARCED model simulation

Before presenting particular unit test applications, we wanted to provide an overview of model simulation. We built a Jupyter notebook called `RunModel.ipynb` (Supp. File 11) to simulate the SPARCED model (Fig. 2). This notebook requires the model SBML (from `CreateModel.ipynb`, Fig. 2A), along with the simulation duration (th), the ligand concentrations (if desired), the name for the output files, and whether the simulation should be deterministic only or hybrid mode (flagD). The rest of the notebook imports necessary packages and model files and runs the simulation (Fig. 2B).

**Figure 2.**
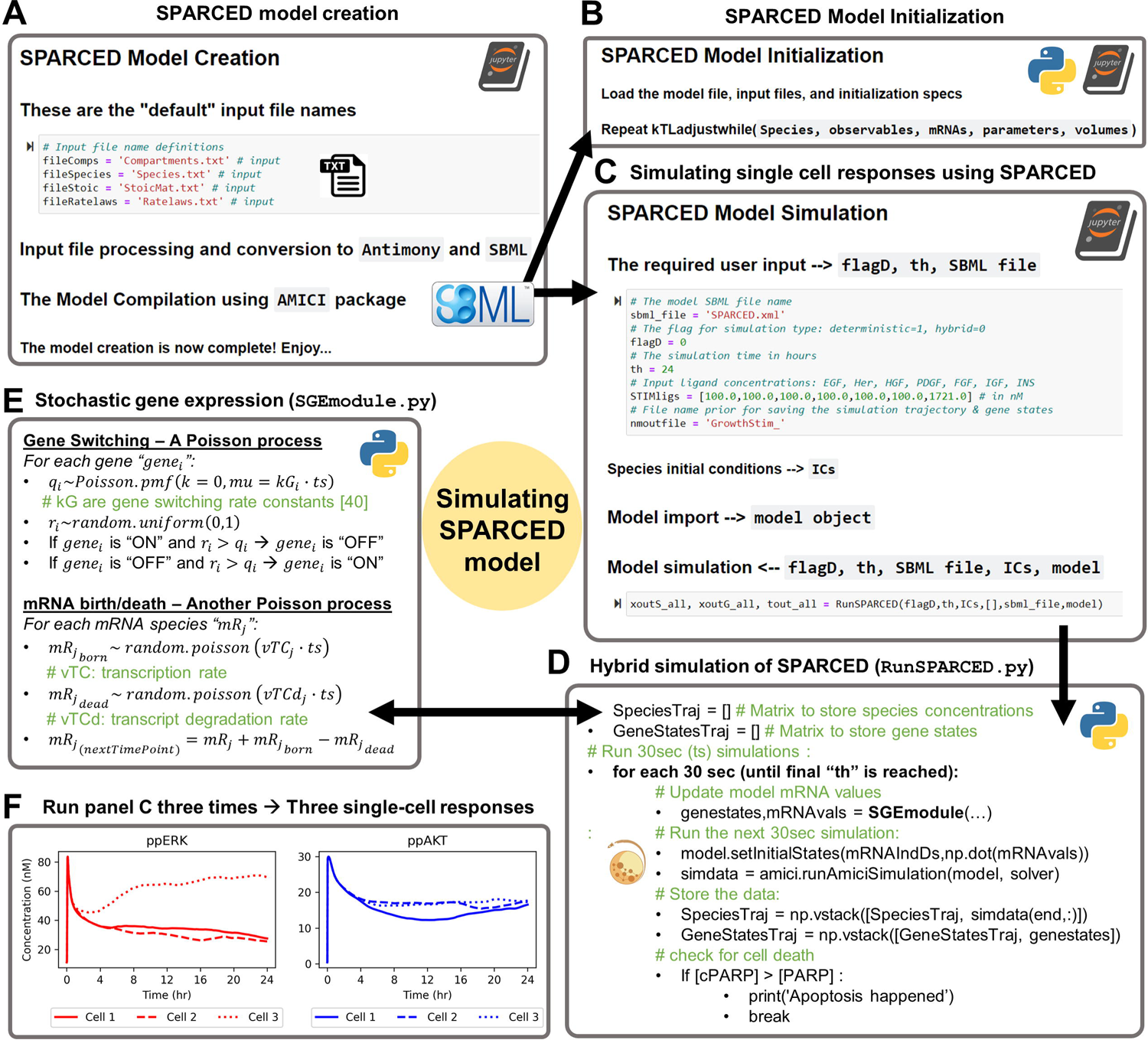
SPARCED-jupyter enables single-cell response simulations using Jupyter Notebooks. (A) The model creation notebook processes the user defined input files and converts them into the model SMBL file. The model (SBML file) is compiled for simulations using the AMICI python package. (B) When a model is generated for a new cellular context (using new omics input data), the model creation step is followed by an initialization step to adjust protein translation rate constants and cell death/DNA damage related parameters. (C) The model simulation starts with specifying and importing the SPARCED model SBML (see panel A). The user defines the model file name and the sets four additional parameters: (i) The flag (1 or 0) to specify if the model should run in deterministic or in hybrid mode (see D and E), respectively. (ii) The time duration in hours for which the model should run. (iii) The vector of ligand concentrations (in nM) to stimulate the cells. (iv) The output file name. Next, the species initial conditions are, by default, read-in from the “Species” input file. Then, the model file is imported, and the model is simulated according to the specified input. The model outputs three matrices of species concentrations over time at every 30seconds, the activation states of genes over time (every 30 seconds), and the time points of simulation in seconds. The two former matrices are saved using the user define file name (iv). (D) The model is simulated iteratively for each 30 seconds, where the current species concentrations are inputs for the gene expression module, which then outputs new mRNA levels to update the SBML model states. The model is then run for another 30 seconds, until the total simulation time reaches the user input (th) or until the cell dies. The cell death is decided based on cleaved-PARP levels surpassing the PARP levels. (E) In the gene expression module, in hybrid mode, the model randomly decides which genes become active or inactive, and which mRNAs are transcribed or degraded. This SGEmodule.py script is called every 30 seconds with updated species concentrations, simulated using the models SBML with AMICI package. (F) When the model is hybrid-simulated three times, the different cell responses are observed. Shown are serum-starved average cells stimulated with full growth media for 24 hours. Plotted are free ppERK and ppAKT species concentrations (nM).

As mentioned, the SPARCED model consists of two modules: deterministic and stochastic. The SBML file forms the basis of the deterministic module whereas the stochastic module describes gene states (active/inactive) and mRNA birth/death events for the genes (Fig. 2C). When run in the hybrid simulation mode, the deterministic and stochastic modules exchange information every 30 simulated seconds (Fig. 2D). The current levels of select protein states can induce changes in gene activation/deactivation or mRNA transcription/decay rates. The newly updated mRNA copy numbers change nascent protein translation rates in the deterministic module (Fig. 2D). When run deterministically, the model does not stochastically sample gene activation or mRNA transcription events, and such simulations correspond to an average cell state.

When the `RunModel.ipynb` notebook is run multiple times in hybrid mode, different single-cell responses are simulated (Fig. 2E). For instance, the activation and phosphorylation of ERK (Fig. 2E left, red lines) and AKT (Fig. 2E right, blue lines) proteins in response to growth factor treatment will show variability across three example cells. Although the amplitude of initial response is similar for all three cells, the longer-term responses are quite different. Our previous analyses showed that such single-cell heterogeneity in the initial concentrations of these proteins could help predict cellular fate, namely cell division (40). These jupyter notebooks provide a simple interface to interact with the SPARCED model.

### SPARCED model Unit Testing: deterministic

We first tested agreement between deterministic Bouhaddou2018 and SPARCED model simulations. The SPARCED model simulations recapitulated the response of an average (deterministic) cell under different stimulation conditions, to within simulation error (Fig. 3A). As an example, we highlight SPARCED model simulations of the cell response (MCF10A cells) to treatment with EGF alone or EGF+insulin (Figs. 3B and S12). Treating growth factor and serum-starved MCF10A cells with EGF and insulin induces activation of ERK, AKT, and their downstream signaling partners, which together influence cell proliferation (40,70,71). The Bouhaddou2018 model showed that compared to single ligand treatments, EGF+insulin stimulation increases and prolongs AKT and its downstream EIF4EBP1 phosphorylation (Fig. 3B).

**Fig 3.**
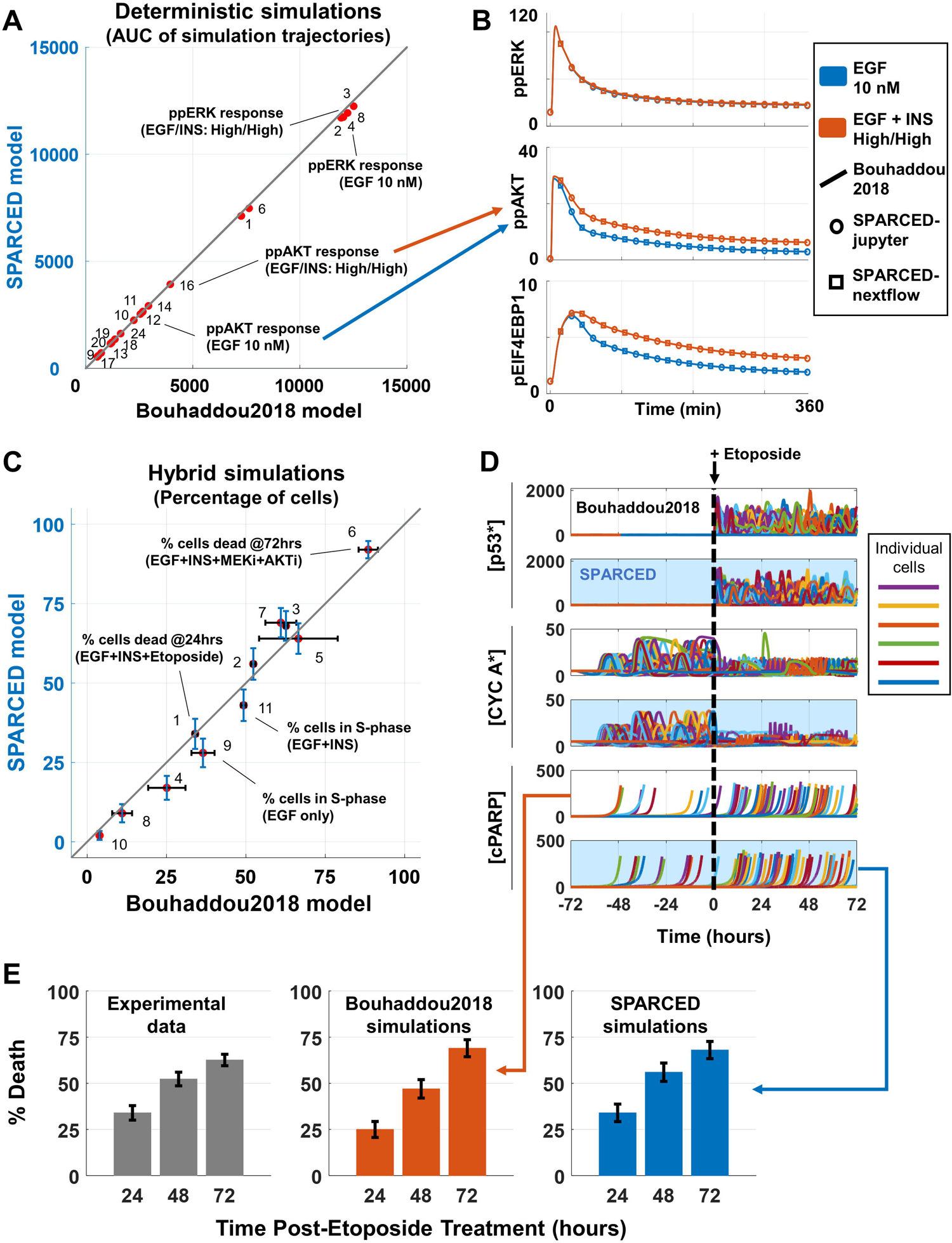
SPARCED model recapitulates experimental observations and deterministic/hybrid (deterministic + stochastic) simulation results of the Bouhaddou2018 model. (A) Summary of comparisons of SPARCED model deterministic simulations to Bouhaddou2018 model simulations. The area under the curve (AUC) values of each simulation (see Fig. S12) are calculated and plotted for the two model results. (B) Simulation results from Bouhaddou2018 model (line) and SPARCED-nf model (triangle) run on Kubernetes cluster workflow are the same as SPARCED model (circle) results. Comparisons of selected panels from (A) are shown only. (C) Experimental and stochastic simulation results from Bouhaddou 2018 model are reproduced by SPARCED model simulations. Each dot is a different condition, explained in Fig. S13A. Error bars show experimental or simulation standard error of the mean. Simulations are of at least 100 cells, and three independent experimental observations where applicable. (D) Stochastic simulation of 100 cells recapture protein level trajectories (active p53, Cyclin A, and cPARP) from older model qualitatively. Panels with blue background are SPARCED simulations and white background panels are from Bouhaddou2018 model. 100 stochastic cells are stimulated with EGF+Insulin for 72 hours before Etoposide treatment for another 72 hours. Etoposide is stimulated also with EGF+Insulin. Results for Etoposide treatment without prior growth factor stimulations are shown in Fig. S13B. (E) Quantification of results in (D) shows that SPARCED model simulations coincide with earlier observations in percentage of death induced by etoposide treatment. See Fig. S13C for the effect of no growth factor stimulation before Etoposide treatment. Bars represent mean ± s.e.m.

The simulation results from the Bouhaddou2018 model (the solid lines) and SPARCED model (circles) are indistinguishable. The SPARCED-nf implementation, which runs on a high-performance cloud computing infrastructure, similarly reproduces the original simulation data (Fig. 3B, triangles). These results, together with all other deterministic tests in Table 1 (Figs. S3-8 and 11), confirm that the SPARCED model recapitulates the Bouhaddou2018 model simulations and unit tests in deterministic settings. Thus, the simple input file structure combined with automatic model generation is equivalent to the prior MATLAB instantiation in this regard.

### SPARCED model Unit Testing: stochastic (hybrid)

Next, we evaluated the SPARCED model for stochastic unit tests in single cell simulations. Each single simulated cell has different initial protein levels and dynamics due to stochastic gene expression, and thus may respond differently to the same treatment. The SPARCED model stochastic simulations closely matched Bouhaddou2018 model results, to within simulation error (Figs. 3C and S13A). As an example, we highlight here how single cells respond stochastically to DNA damage. Etoposide, a chemotherapy drug, induces double-and single-stranded DNA damage, causes cell cycle arrest, and leads to cell death (72). Experiments showed that in the absence of EGF and insulin (to promote cell cycle exit), there is minimal etoposide-induced cell death (Fig. S13B-C) (40). However, in the presence of EGF and insulin (to drive cell cycle progression), etoposide-induced cell death increases over time and reaches around 60% of the cells (Fig. 3D-E). Simulating etoposide treatment of cycling cells induces robust p53 pulses, disruption of Cyclin A dynamics/cell cycle arrest (Fig. 3D), and more cell death relative to non-cycling cells (Fig. 3E). These results closely match experimental data and Bouhaddau2018 simulations. We conclude that SPARCED model captures DNA damage induced single-cell death percentage and cell cycle state-dependent effect of etoposide. The SPARCED model also passed all other stochastic/hybrid unit tests (Table 1, Figs. S2, 6,7,9, and 10).

### SPARCED model Unit Testing: context change

Different cell types have different mRNA and protein expression levels, and many mechanistic models assume that it is different expression levels that drive different phenotypes, as opposed to changes in biochemical rate constants. These constants are based on biophysical events like binding, which are based on molecular structures. Here, we tested the ability of the SPARCED model to be re-“initialized” to study different cell types by changing initial levels of total proteins and mRNAs without changing the model topology. Thus, we introduced a protocol to enable SPARCED model context change (Fig. S14A and Supp. Files 12 and 13). In short, `OmicsData`, `Species`, and `Ratelaws` input files are updated with new cell line information, including mRNA levels, protein/species levels, and constitutive translation rate constants. Then, the new model is created by running the “`createModel`” Jupyter notebook or by submitting a new SPARCED-nf job.

The re-calibration step for context change followed in Bouhaddou2018 model was called *Initialization*, where protein-specific translation rates and key parameters important for cell decision making are estimated to ensure agreement with new omics datasets and expected phenotypic behavior with respect to proliferation and apoptosis. Here we also provide a new, python-based version of the *Initialization* procedure for SPARCED models (see Computational Methods), where the outputs are species concentrations and rate parameter values updated in a new SBML file. Here, to test the drug combination response differences in different cell lines, we changed SPARCED model context (i.e. parameter values and species concentrations) by initializing the model to the U87 glioma cell line. Following the protocol outlined in Fig. S14A, we replaced MCF10A cell line values in the input files with values from U87 cell line data.

U87 cells are PTEN-deficient and more sensitive to AKT inhibition compared to MCF10A cells (40). Both cell lines show minimal sensitivity to MEK inhibition alone and AKT & MEK inhibitors are both needed to kill MCF10A cells. In contrast, AKT inhibition alone is sufficient to kill U87 cells. To simulate the U87 cell response to AKT and MEK inhibitors, we first updated the `OmicsData` input file (Supp. File 1) using U87 mRNAseq data from Bouhaddou2018 model (Supp. File 14). Here, we did not have U87 cell line proteomic data and estimated the initial total protein levels using the new mRNA levels and gene-level mRNA/protein ratios from MCF10A data (Supp. File 15). We set PTEN translation rate to zero and set values of rate parameters dictated by *Initialization* in the `Ratelaws` input file (Fig. S14B and Supp. File 12). Additionally, we provide an improved Python based initializer `InitializeModel.ipynb` notebook (Supp. Files 16 and 17), which re-creates (Fig. S15) the un-stimulated steady-state initial conditions for species and adjusts translation rate constants using cell-line specific initialization input file (Supp. File 18). We also updated the species initial conditions in `Species` input file using steady-state values for U87 cells from the Bouhaddou2018 model (Supp. File 19). We created a new model SBML file (SPARCED_U87) using the updated input files. SPARCED_U87 model simulations of response to MEK and AKT inhibitors reproduced the Bouhaddou2018 model results and experimental observations (Fig. S14C). We conclude that changing model context by changing input files accomplishes the goal of easy model alteration to study of different cell types.

When the cellular context (omics input data) for the SPARCED model is changed, all appropriate Unit Tests should pass. We expect that addition and alteration of the list provided (Table 1) will accommodate increasingly different prior knowledge about the new context. Examples of such information include cell line mutations, growth condition differences, or tumor cell behavior.

### Illustrating easy model expansion by application to the IFNγ pathway: SPARCED-I

The SPARCED model reproduces original model results and experimental observations, but how can we use the new simple model expansion capabilities? Here, we focused on experimental observations that interferon-gamma (IFNγ) inhibits MCF10A cell proliferation. Specifically, as part of a much larger LINCS consortium effort to deeply profile the MCF10A cell line dynamic response to perturbations (synapse.org/#!Synapse:syn12526172), we observed that IFNγ inhibits EGF-induced cell proliferation (Fig. 4A). We wanted to use the SPARCED model expansion functionality to evaluate the suitability of candidate mechanisms by which IFNγ might inhibit proliferation.

**Fig 4.**
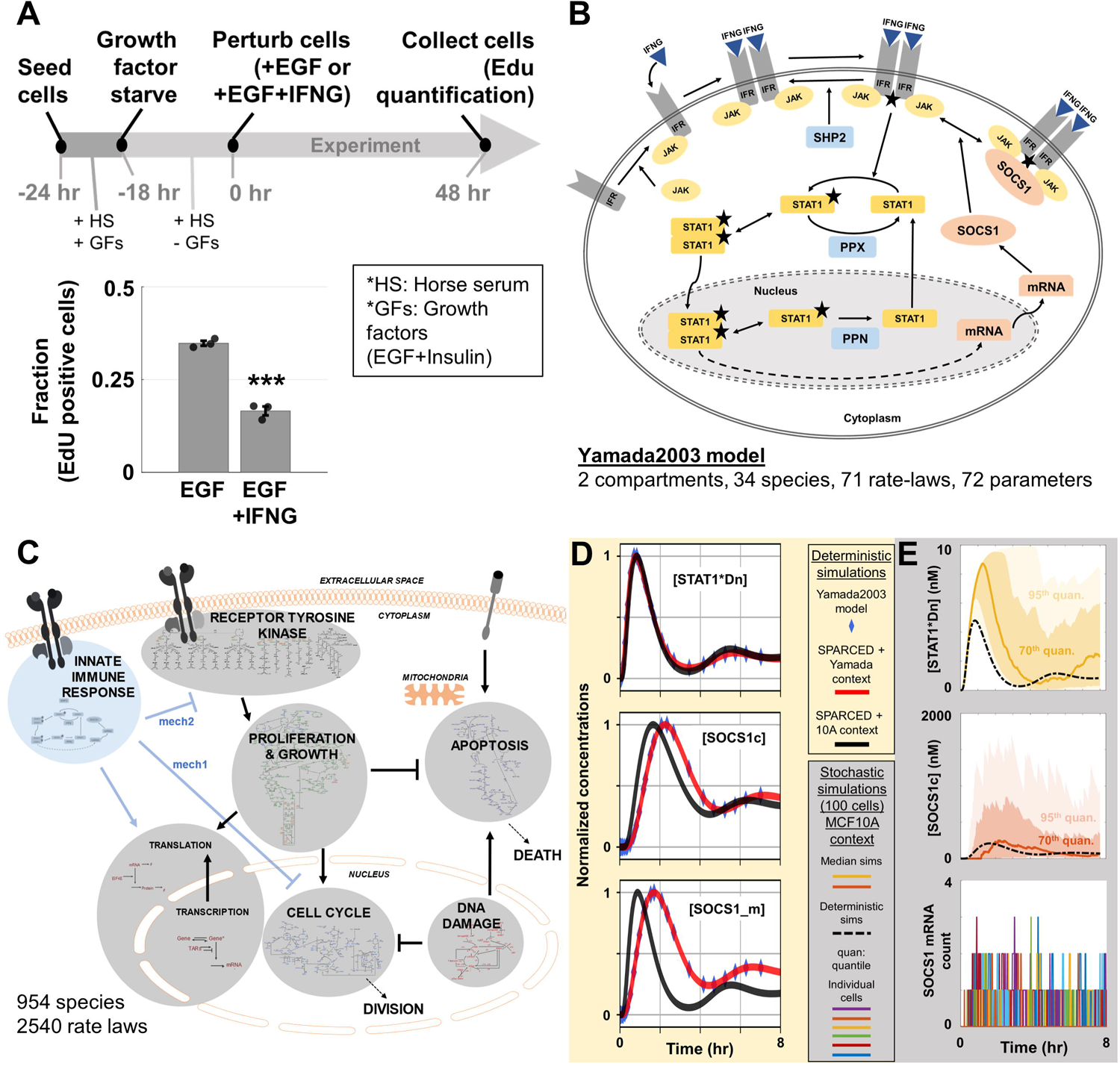
SPARCED model is enlarged to include interferon-gamma (IFNγ) signaling pathway. (A) Experiments showed that IFNγ treatment decreases cell proliferation induced by EGF alone. Fraction of EdU positive cells at 48 hours after EGF alone or EGF+IFNγ treatment are calculated. Bars represent mean ± s.e.m of three independent biological replicates. Significance tested using two-sided two-sample t-test, where * indicates p-value < 0.05, *** indicates p-value < 0.001. (B) Yamada2003 model schematic of IFNγ pathway added into SPARCED model. (C) Overview of the added IFNγ-IFNGR pathway in relation to the Bouhaddou2018 model pathways. The “mech1 and mech2” links are candidate mechanisms tested in the next section. (D) Simulations of the Yamada2003 model, SPARCED-I model, and SPARCED-I model with MCF10A context show qualitative and quantitative agreements. The Yamada2003 model results (blue diamonds) are obtained by running the model file in COPASI (61). (E) 100 stochastic cell simulations (area plots) and the deterministic (dashed-black lines) simulation of 10 nM IFNγ stimulation are shown. Colored dark lines represent median cell trajectories, dark and light-colored regions represent 70^th^ and 95^th^ quantiles, respectively. The right-most plot shows mRNA count of SOCS1 in each cell, colored differently.

We thus merged a model for interferon-gamma receptor (IFNGR) signaling into SPARCED, creating the model variant SPARCED-I. The newly added Yamada2003 model (73) captures how IFNγ binds to pre-JAK-bound receptors, inducing dimerization. The ligand-bound receptor homodimers are activated by JAK, which also phosphorylates STAT1 when bound to the active receptor complex. Activated STAT1 then dimerizes and translocates to the nucleus, inducing transcription of SOCS1. SOCS1 mRNA is exported to the cytoplasm and translated into SOCS1 protein. SOCS1 protein binds to and inhibits activated receptor homodimers. In this model, there are three phosphatases (SHP2, PPX, and PPN) acting on multiple species.

Following the model alteration protocol in Fig. S14A and Supp. File 13, we added 34 new species, 8 corresponding genes, and ∼70 reactions (Fig. 4B) based on the Yamada2003 model (73). New genes (IFNG, IFNGR, JAK2, STAT1, SHP2, SOCS1, PPN, PPX) corresponding to the proteins in the model are added as new rows and mRNA levels are inserted into the `OmicsData` input file. Gene copy numbers are taken as two (40). The genes are also added as new rows to the `GeneReg` file (Supp. File 20). In the Yamada2003 model, activated nuclear STAT1 dimers (STAT1*Dn) induce SOCS1 mRNA transcription and this is captured by adding a new column in the `GeneReg` input file, with the only non-zero element at the STAT1*Dn and SOCS1 gene intersection. Next, each protein, protein complex, and mRNA species are inserted into the `Species` and `StoichiometricMatrix` input files as new rows. Each new reaction is inserted into the `Ratelaws` input files as rows, and into the `StoichiometricMatrix` input file as new columns. This expansion brought the total number of species of SPARCED-I model to 954, and the number of reactions to 2540 (Fig.4C, Supp. File 21).

### SPARCED-I model Unit Testing

The SPARCED-I model, with parameter values from the Yamada2003 model, should reproduce the original results exactly, which we verified (Fig. 4D, red lines & diamonds, respectively). We then modified the mRNA, protein, and compartment volume values to that of MCF10A cell context (data from (40)). However, the MCF10A data (Supp. File 22) had missing values for IFNGR and (arbitrary) phosphatase species PPN and PPX. So, we initialized the concentration of IFNGR as half of JAK2 concentration (the receptor is typically rate limiting (73)). The concentrations of PPN and PPX phosphatases were equal to half of SHP2 concentration in Yamada2003 model, so we updated their values to half of SHP2 concentration in MCF10A cells. In addition to the reactions from the Yamada2003 model, we added new translation (for the new eight genes) and degradation (for all new species) reactions into the model. The rate constants of these extra reactions are initially assumed to be equal to the average of corresponding reactions of SPARCED model genes. Starting from these parameter values, the SPARCED-I model showed unrealistic (ultrafast) receptor activation and STAT1 phosphorylation/nuclear transport rates. Therefore, we varied six parameters that have high impact on STAT1 activation dynamics (see Methods) to approximate the timing (within the first hour) of STAT1*Dn pulses reported in Yamada2003 model (Fig. 4D) and others (74, 75). Changing rate constants in such a manner accounts for the entangled effects of unmodeled cellular context and mechanisms. Tuning these parameters produced expected pulsing times and response behavior of STAT1*Dn, SOCS1, and SOCS1 mRNA levels (Fig. 4D black lines). The final values are updated in the SPARCED-I model file (Supp. File 23).

A key feature of SPARCED-I is its ability to simulate single cell behavior and the above-observed reduction in proliferation induced by IFNγ is inherently a single cell property. As a unit test, we simulated 100 single cell trajectories (Fig. 4E) of SPARCED-I model and concluded that SPARCED-I model recapitulates observations from earlier models and passes all unit tests.

### SPARCED-I model variant analysis: hypotheses testing

Next, we wanted to use the expanded model to help us interpret the experimental observations. H*ow does IFN*γ *inhibit EGF-induced cell proliferation?* The SPARCED model captures regulation of cell proliferation via the ERK and AKT pathways. Growth-inducing ligands, like EGF, bind to and activate receptor tyrosine kinases (RTKs), which in turn leads to upregulation of AKT and ERK phosphorylation. The two pathways together induce upregulation of cyclin D through cJUN, cFOS, and cMYC activities (40,70,76).

In the literature, there are different mechanisms by which IFNγ was suggested to play a role in cell proliferation (77–80). The SPARCED-I model enabled us to evaluate these hypotheses for consistency with experimental observations (Fig. 4A), where the simulation steps are matched to the experimental setup (Fig. 5A). The first proposed mechanism (mech 1, Fig. 5B) was based on reports that activated STAT1 can induce p21 (cell cycle inhibitor) expression (77, 81). This mechanism was modeled (Supp. File 24) by modifying the `GeneReg` input file (Supp. File 21), where active STAT1 dimers can increase p21 transcription rate. The SPARCED-I model already had p21 transcriptional activation by p53, and the rate constant for transcriptional activation of p21 by STAT1 was taken to be equal to the parameter values from the p53 mechanism.

**Fig 5.**
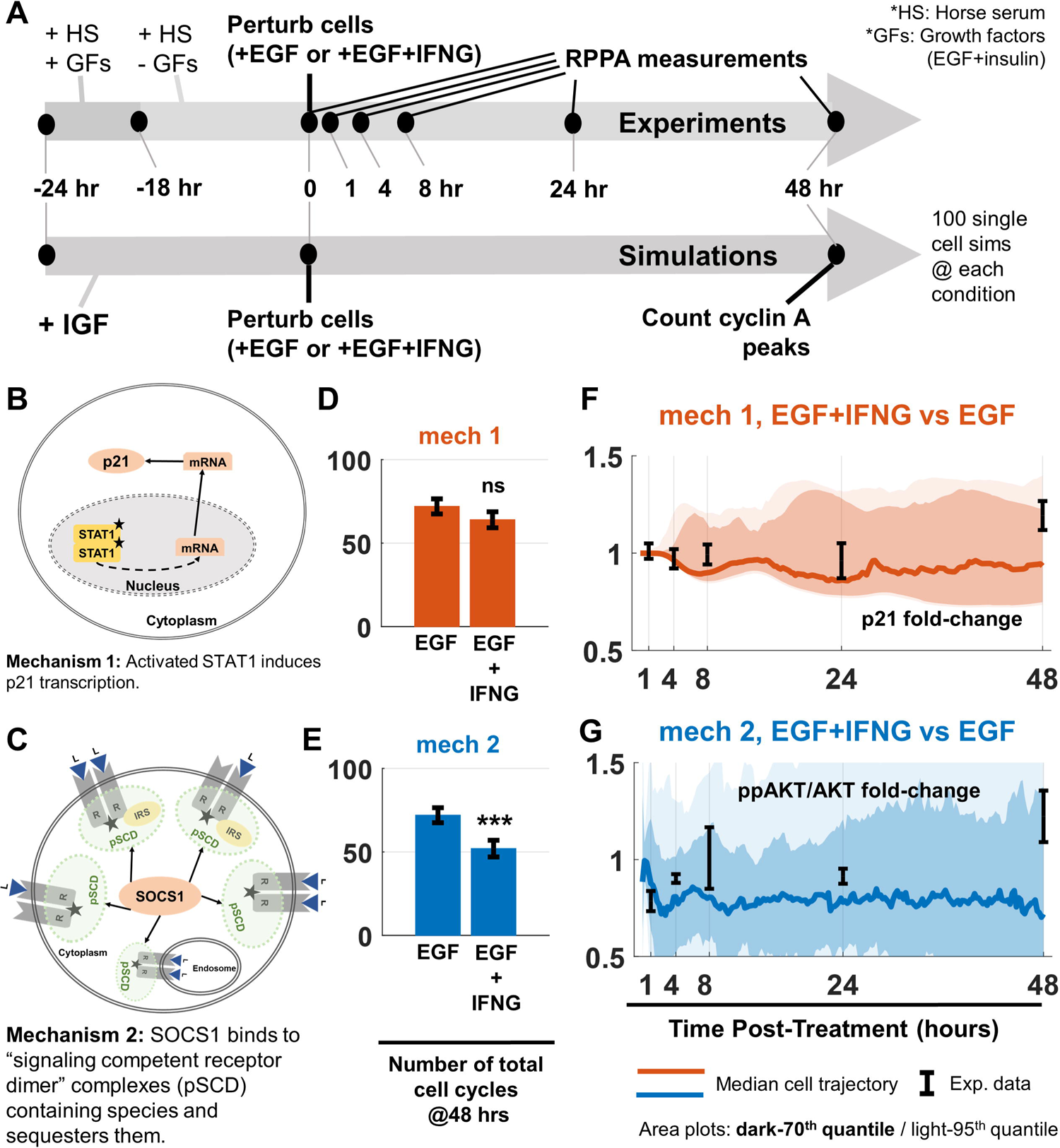
Study of SPARCED-I model enabled us to propose how IFNγ treatment decreases cell proliferation induced by EGF. (A) Experimental setup and simulation workflow for SPARCED-I model variant analysis. See Methods. (B) Candidate mechanism schematic 1: activated STAT1 induces p21 transcription. (C) Candidate mechanism schematic 2: SOCS1 sequesters activated ligand-receptor complexes. (D) Simulation of 100 stochastic cells for SPARCED-I-mech1 and (E) SPARCED-I-mech2 models showed that SOCS1 sequestration of activated receptor complexes can better explain the IFNγ effect on cell proliferation inhibition, compared to experiments shown in Fig. 4A. The bar plots show the total number Cyclin A peaks calculated at specified time points for the starting 100 cells. (F) Normalized p21 levels do not show a significant change when IFNγ is included in addition to the EGF. (G) Normalized ppAKT levels show a significant decrease after IFNγ treatment, when EGF+IFNγ Mechanism 2 simulations are compared to EGF alone case. RPPA data are shown in black error lines, from three independent replicates. Colored dark lines represent median cell trajectories from simulations, dark and light-colored regions represent 70^th^ and 95^th^ quantiles, respectively. Significance tested using two-sided two-sample t-test, where * indicates p-value < 0.05, *** indicates p-value < 0.001. (F-G) Data from synapse.org/#!Synapse:syn12526172.

The second mechanism involves the negative regulator of IFNγ signaling, SOCS1. SOCS1 protein has different binding domains, including SH2 domains (82, 83). SH2 domains bind to phosphorylated tyrosine residues on other proteins (84, 85). It is proposed that SOCS1 not only binds to activated IFNγ receptors, but also to many other activated receptor complexes with free phosphorylated tyrosine residues (mech 2, Fig. 5C) (83,86,87). Thus, IFNγ-induced SOCS1 protein can bind to growth factor-activated receptor complexes (or the so-called signaling competent dimers – pSCD) and prevent further downstream signaling by sequestration. This mechanism was modeled by adding SOCS1 binding to activated receptor complexes (pSCDs) reactions in the `Ratelaws` input file. The SPARCED-I mech2 model contained 1302 species (348 new) and 3584 reactions (1044 new) (Supp. File 25). GRB2 proteins also contain SH2 domains that bind to tyrosine phosphorylated receptors (pSCDs) and the rate constants of SOCS1 interaction with all these complexes are taken as the average of such parameters of GRB2 complexes.

Before evaluating these model variants, the initial conditions must be set. The SPARCED-I model initial conditions are based on serum and growth factor starved MCF10A cells. However, the experiments with IFNγ (Fig. 4A) were done in media with horse serum. Horse serum upregulates ppAKT levels by four-fold (Figs. S16 and S17), possibly thorough IGF/IGF1R pathway (88–90). Including 0.02 nM IGF meets this basal activity increase constraint, and therefore is included in simulations prior to and during simulated EGF and IFNγ treatments (see Methods and Figs. S16-19).

After creating the SPARCED-I Mechanism 1 and 2 variants and defining the simulation conditions, we stochastically simulated both models with IGF treatment for 24 hours for 100 different single cells to provide a baseline. Then, either EGF or EGF + IFNγ are added for an additional 48 hours for each cell (Fig. 5A). As a simulation metric for cell proliferation, we counted and summed the number of cyclin A peaks in each of the 100 starting cells for the 48 hours (Fig. 5D and E-orange and blue bars). Mechanism 2 leads to a significant decrease in the number of cell cycles in IFNγ treated cells, whereas Mechanism 1 did not cause a change in EGF and EGF+IFNγ conditions.

Exploring further, we saw that Mechanism 1 did not lead to significant increases in p21 protein levels (area plots in Fig. 5F). While it is unexpected that the model predicted no significant increase in p21 levels despite a clearly added mechanism, if the parameter value for STAT1 induction of p21 transcription is changed enough (see Supplemental Equation 12 in (40)), simulations do indeed show IFNγ treatment induces p21 (Fig. S20). However, experimental RPPA data from the LINCS Consortium confirm that total p21 levels do not significantly change over the course of 24 hours in EGF+IFNγ treated cells compared to EGF alone stimulation (error bars in Fig. 5F, synapse.org/#!Synapse:syn12526172). The experimental 48 hour time point showed elevated p21 levels, which is different from simulations. We conclude that the original parameter choices are appropriate and that the putative p21 mechanism is therefore unlikely.

We investigated responses of simulated cells with Mechanism 2 to explore why fewer simulated cells enter the cell cycle (Fig. 5E). We saw a decrease in MAPK activation in IFNγ treated cells, supported by experimental data, but in both Mechanisms 1 and 2 (Fig. S21). Then, looking at AKT responses, we saw lower in silico AKT phosphorylation level in IFNγ treated cells of only Mechanism 2, compared to only EGF stimulated cells (Fig. 5G area plots). The greatest decrease occurs at around one hour after stimulation and stays decreased for 48 hours. We compared this simulation result to experimental RPPA data (synapse.org/#!Synapse:syn12526172), which also show decreased ppAKT levels at 1-4 hours post-ligand treatment (Fig. 5G error bars). The variability in the 8 hour measurement was large, and the 24 hour measurement showed still reduced ppAKT. By 48 hours, ppAKT is upregulated in the experimental data, which the model simulations do not account for. Is the somewhat subtle decrease in AKT activity for ∼1 day enough to prevent cell cycle entry in MCF10A cells? Bouhaddou et al. showed that a slight change in time integrated AKT dynamics can tilt the cell cycle progression decisions significantly (see Fig. S11 and (40) for further details) and here we saw ppAKT levels show a statistically significant decrease after IFNγ treatment, which may explain fewer proliferative cells. Taken together, these results suggest that: (1) the SPARCED model formalism can be used for simple implementation of large-scale mechanistic model-based hypotheses testing, (2) STAT1 induction of p21 expression levels is an unlikely mechanism in this MCF10A cell context, (3) crosstalk between IFNγ and EGF pathways may occur through SOCS1 sequestration of activated receptor domains, and (4) SOCS1-induced inhibition of AKT activation may contribute to proliferation suppression.

## DISCUSSION

Here, we have re-created one of the largest mechanistic models in the literature, using our new python-based creation and simulation pipeline. Enabling large-scale, single-cell modeling as a data integration tool, our pipeline is based on structured and easy to modify input text files and uses Jupyter notebooks or scripts (for scaled cloud-computing) to create and simulate model files. It also enables easier model alteration (species/rate law or parameter value changes), omics data integration, and model variant vs. hypotheses testing. Our exemplar model, called SPARCED, is available online on GitHub (github.com/birtwistlelab/SPARCED). SPARCED can serve as a basis for creating context-specific (personalized) model variants, studying virtual cell population responses, and as a building block towards whole-cell-scale models.

First, we showcased the use of SPARCED model by changing the cellular context of the model from MCF10A breast epithelial cells to U87 glioblastoma cells, by only replacing parameter values in three/four input files. Although we used previously calculated values from the Bouhaddou2018 model to show reproducibility of the subsequent analyses, it is notable that SPARCED pipeline correctly creates and simulates a large-scale mechanistic model file only from an altered set of text-based input files. Additionally, we provided a new version of the Initialization script (Supp. File 16) that utilizes another cell-line specific input file (Supp. File 18) to calibrate the model initial conditions. The initialization allows distribution of total protein and mRNA level omics data across all model species and estimates data-driven, cell-line specific translation rate constants. As a customizable set of steps, the initialization sustains user defined phenotypic responses, like cells not going into apoptosis or cell cycle without growth stimulation. Importantly, the procedure accommodates mRNA input alone (without proteomics data) and calculates total protein levels using gene-level mRNA-to-protein ratios from the default MCF10A values. The output of the initialization procedure are species concentrations and parameter values deposited into a new SBML file, which is exported with a new name and re-compiled using AMICI.

Secondly, we investigated two candidate mechanisms to explain how IFNγ can inhibit EGF-induced cell proliferation. By creating and testing two mechanisms (by only changing the input files), we hypothesized that IFNγ-induced SOCS1 sequestration of activated receptors is a more likely mechanism than IFNγ induction of cell cycle inhibitor protein p21. Besides these two mechanisms tested, there are others in the literature, like the positive feedback of STAT1 inducing STAT1 and IRF1 transcription (78), or the inhibition of Bcl-2 by STAT1 (77). However, here, we only focused on demonstrating the capabilities of SPARCED pipeline to easily create and test model variants to help explain one of our experimental observations, where testing all possible mechanisms was out of our scope.

Many existing big models are constructed in complicated and hard-coded ways and are not available in standard modeling formats, like SBML. For instance, the Bouhaddou2018 model we used as our starting point was custom coded in MATLAB with tens of different script files with thousands of lines. Although the model performance was optimized for its topology, alteration and expansion of the model was extremely difficult. However, models, especially the large-scale and clinically relevant mechanistic models, must become easy to formulate, understand, and disseminate for reproducibility, re-useability, and applicability in clinical decision making. Here, the model construction pipeline and the SPARCED model contributes to this need by being built upon structured and annotated input files, by using open-source packages, and by being available publicly on GitHub.

One key advantage of the SPARCED model format is its potential compatibility with RBM. The reactions and species created by RBM software can be incorporated (manually or programmatically) into the SPARCED model input files. Although existing RBM software can export models in SBML format and enable multiple features, the SPARCED models enable single rate parameter changes and inclusion/exclusion of individual rate laws at the input file level. Then, the SPARCED-nf pipeline can be used to study large-scale variant analysis or to do parameter scanning. One main goal of the AMICI package (91) is enabling large-scale parameter estimation, and our choice to use this package was to enable such future endeavors when needed. Combining this idea to test consistency across multiple datasets, users can search for best-fit models or pinpoint discrepant datasets given the model topology (28).

Another advantage of the SPARCED model will be its ability to integrate multiple omics datasets into a large-scale mechanistic model, creating a “personalized” model variant reflective of another cellular or patient context (26, 92). With the *Initialization* procedure linked to our pipeline, users can incorporate mRNA, copy number variation (CNV), and even proteomics data from established databases like CCLE (93), TCGA (94–96), HPA (97–99), and Cellosaurus (100) into the input files programmatically (Fig. S14) and test changing the initial conditions of the model using the same network structure (Fig. 4).

The SPARCED model encodes intrinsic stochasticity of total protein levels and mRNA numbers in its hybrid simulation mode, making it unique (together with the Bouhaddou2018 model) to offer stochastic as well as deterministic simulation settings. Although other tools such as COPASI offer hybrid (deterministic + stochastic) simulation settings, our approach always treats the gene expression module as stochastic, and the events modelled as Poisson processes. COPASI uses next-reaction-method (101) for the part it determines as stochastic based on molecule numbers of the interacting species. However, as the developers stated, such implementations tend to be inefficient and take prolonged simulation wall-times. The single-cell capability of SPARCED allows one to capture some important aspects of cell line and tumor heterogeneity compared to an average cell condition (the way many mechanistic models are built). Users can leverage this feature to simulate virtual populations and study a cell population response to drug treatment, which is often a single-cell readout as are most cellular phenotypes. However, such simulation settings require larger computational resources and thus model compatibility for high performance computing environments. The SPARCED model is built to be compatible with cloud computing, where it can be used to simulate thousands of single cells with single job execution (see Methods).

There are, of course, remaining challenges. One such task is to explore simulations of *spatially aware* single cells. Currently, the SPARCED model captures intrinsic heterogeneity of cells (by having stochastic gene switching and mRNA birth/death events) but these cells cannot “talk” to each other. In the future, by having scenarios where spatial orientation of cells are recorded and the secreted or stimulated molecules are shared between them, we can better capture tissue microenvironment and heterogenous pharmacokinetics (102, 103). Related to this first task, the second challenge is to create and simulate scenarios with multiple cell types (i.e. models trained on data from different subtypes of cells) or defining events to capture differentiation of cells. For example, one may be able to use single cell RNAseq data to train SPARCED-like models to enable tissue-level simulations with the critical cell types in the proper geometric locations. This overall vision would enable spatially aware, single-cell level, large-scale mechanistic models trained on individual patient data for *in silico* drug screening. The pipeline presented here is an important step towards this goal.

Another challenge to achieve using large-scale models is a whole-cell level mechanistic model for mammalian cells (56, 104). With our approach, the SPARCED model can be enlarged using other small-scale models for pathways and mechanisms not currently included in the model. By utilizing the unit testing approach, one can then verify the model performance and get larger, more comprehensive models. The open-source framework presented here increasingly facilitates community contribution for model context-change and parameter tuning based on new experimental conditions.

Our introduced method of large-scale mechanistic model construction, and the SPARCED model as a basis, will enable researchers to more easily create and manipulate new model versions, test different mechanisms of action to interpret experimental observations, and change the model’s cellular context. The models created by our pipeline can incorporate multiple (omics) datasets, providing non-“black-box” data integration and modeling. These SPARCED models additionally provide single-cell level simulations, compatibility with cloud computing, and human-interpretable & annotated model files in SBML format. The SPARCED model now can more easily be re-used as one of the largest mammalian-cell mechanistic model in the literature and serves a primer role in creation of context-specific, hypotheses testing, and expandable models. In conclusion, the SPARCED model format contributes towards important foundations of reusable big models, paving the way towards personalized mechanistic models for data integration.

## MATERIALS AND METHODS

### Computational Methods

#### The Bouhaddou2018 model

The Bouhaddou2018 model (Fig. 1A) is one of the largest single-cell mechanistic models for mammalian cell signaling regulating proliferation and death. The first version of the model used as a test case in this work was written in MATLAB (The MathWorks, Inc.) (40). The model is a hybrid of deterministic and stochastic modules. The deterministic module describes the concentration dynamics of 774 proteins, protein complexes, and post-translationally modified species through 2449 reactions using the Sundials CVODEs package for simulation (105). The stochastic module describes gene state (active/inactive) and mRNA birth/death dynamics for 141 genes. The deterministic and stochastic modules exchange information every 30 simulated seconds. In short, the current levels of select protein states can induce changes in gene activation/deactivation and/or mRNA transcription/decay rates. The newly updated mRNA copy numbers change nascent protein translation rates in the deterministic module. See (40) for further details.

#### The SPARCED model

We converted the Bouhaddou2018 model into a Python + SBML (59) format (Fig. 1B). The deterministic module is ultimately encoded in an SBML file (.xml) whereas the stochastic module is written in Python. A foundational and important feature of this recoding effort is that the SBML file is generated from a small set of simple structured input text files (Fig. 1C) via Python scripts. Introduction of such structured input files and associated Jupyter notebooks enables simple alteration of model structure and/or parameter values, for example turning on/off certain interactions. The input files also enable rigorous annotation of model features using, for example, ENSEMBL (106) and HGNC (107) identifiers, which is seldom done in such mechanistic modeling.

#### Input files

There are six SPARCED model input text files (tab separated values), each with a defined structure as detailed below. The user can change these files to create and compile a new model. `OmicsData`: This file (Supp. File 1) includes the gene copy number, mRNA copy number, and proteomic data. This input file also contains rate constants for the stochastic module and initialization procedure. Each row of the file corresponds to one gene and the columns are different data types. The first column is gene name (HGNC identifiers), the second column is gene copy number, the third column is mRNA molecule copy number per cell (mpc), the fourth and fifth columns are rate constants of gene inactivation and activation respectively (s^-1^), the sixth column is constitutive transcription rate constants (molecules per second), the seventh column is maximal transcription rate constants (molecules per second), the eighth column is mRNA degradation rate constants (s^-1^), the ninth column is protein copy number (mpc), the tenth column is protein half-life parameters (seconds), and finally the eleventh column is the translation rate constants (s^-1^). These latest set of rate constants are from literature and provided for genes for which our omics input lacked protein level data. All the rate constants are taken from the Bouhaddou2018 model. Users can add new rows to this file, using RNAseq data to estimate mRNA levels for the genes to be added (40). When adding genes (rows) to the model, a reasonable starting point for rate constants (or other values), in the absence of any other data, is to use median values from the genes/parameters currently in the model.

1. `Species`: This file (Supp. File 2) contains information about the species in the deterministic module. Each row corresponds to one species (protein, protein complex, post-transcriptionally modified species). Transcripts (in nM) are also included in this file because they are regarded as species with updated concentrations in the stochastic module every 30 seconds and are used in translation rate laws. The first column is the species name. Names can be arbitrary so long as they are unique in the model. Importantly, the name list needs to match the first column in the `StoichiometricMatrix` file described below. The second column is the species home compartment. The home compartment of a species defines its cellular localization. A species can reside in a compartment defined in the `Compartments` input file: currently Cytoplasm, Mitochondria, Nucleus, or Extracellular. The third column is initial condition in nM units, with respect to the home compartment volume. These values are taken from the Bouhaddou2018 model, post-initialization. The fourth column is a comma separated list of ENSEMBL gene identifiers corresponding to gene products present in the species.
2. `Ratelaws`: This file (Supp. File 3) has a row for each reaction in the deterministic module. The first column is the unique (arbitrary) name of each reaction. Currently, we named each reaction based on the related sub-module (e.g. vA1-87 for Apoptosis and vC1-104 for Cell Cycle). The number and order of rows in this file should match the columns in the `StoichiometricMatrix` input file defined below. The second column in this file contains the home compartments for the reactions. The designated compartments should be one defined in the `Compartments` input file: currently Cytoplasm, Mitochondria, Nucleus, or Extracellular. The home reaction compartments define the effective search volume for each reaction and is used to rescale concentrations when appropriate. Note that both species and reactions have home compartments defined, where a species can participate in a reaction defined in a different compartment. For instance, the EGF binding to EGFR reaction occurs in extracellular space (volume V_e_), where EGF’s home compartment is the extracellular space and EGFR’s home compartment is cytoplasm (V_c_). A volumetric correction for EGFR concentration in this rate law is done by multiplying by the ratio of V_c_/V_e_. The third column can have either a number or a reaction formula. If it is a number, it means the corresponding reaction is mass-action type, and the number is the rate constant for that reaction in units of nM and seconds. Note that the reactants and products are defined in the `StoichiometricMatrix` input file. If the third column is a formula, it means the reaction will follow that rate law, and the next set columns in that row are the values of each parameter defined in the formula in the third column, again in units of nM and seconds. The rate law can include any species name described in the Species input file. The parameter names in the rate law should start with “*k*” and be unique in that formula. We distinguish multiple parameter names with an underscore and ascending list of numbers (e.g. kA_1, kA_2). During model generation, all parameter names in this file are re-named in an ascending order based on the number of rate laws. The full list of parameter name/value pairs are outputted into a new file (`ParamsAll`) for user reference.
3. `StoichiometricMatrix`: This file (Supp. File 4) defines the reaction stoichiometry, and therefore the reactants and products of model reactions. The rows correspond to the `Species` input file and the columns correspond to the rows in the `Ratelaws` input file. Here, the species and rate law names should match the names defined in Supp. Files 2 and 3. Each element (starting at the second row and second column index) has a stoichiometric coefficient (typically − 2, −1, 0, 1, or 2), where negative sign indicates reactants, and positive sign implicates products of a reaction.
4. `GeneReg`: This file (Supp. File 5) describes transcriptional activation and inhibition interactions, where rows correspond to genes (the same order as the first column of Supp. File 1) and columns to species that are defined as activators or repressors of transcriptional activity. The first column is gene name (HGNC format). There are currently seven more columns in this file, each corresponding to one species defined as an activator or a repressor (e.g. p53 induces p21 transcription or AP1 inhibits cFOS transcription). A single value of zero indicates no effect. A non-zero entry in row i and column j denotes that species j regulates gene i transcription. The non-zero entries have the form “A; B”, where “A” is the hill coefficient and “B” is the half-maximal concentration of the species “j” effect. To simplify the input file structure, we use positive values of “A” to denote activation, and negative “A” values to denote inhibition. This file is used by the stochastic module script to update mRNA levels. To add additional transcriptional regulators (activators or repressors) into the SPARCED model, users should add as many columns as new regulator species and populate the columns with corresponding rate constants.
5. `Compartments`: This file (Supp. File 6) contains the names of compartments in the model (first column), the volume of the compartment in liters (second column), and the corresponding GO-term of the compartment (third column). The compartment names should match the compartment names listed in `Species` and `Ratelaws` input files.
6. `Observables`: This file (Supp. File 7) contains information about model observables. Each observable corresponds to the compartmental-volume-corrected summation of all formats of a protein. There are 102 observables defined (columns) for the model species (rows) in this file. The entries are either 1 (the species in the row is part of the observable in the column) or 0 (otherwise). The “`createModel`” Jupyter notebook (Supp. File 8) uses this file to define an observables variable as an input for the `AMICI` model compiler.
7. `Initializer (Optional)`: This file (Supp. File 16) contains information used for model initialization. Species concentrations (columns 1-2), mRNA level adjustments (columns 3-4), parameter values (columns 5-7), observables to exclude from translation rate adjustments (column 8), and single parameter scan range (columns 9-11) are populated for each step of initialization. The steps used here are shown to work to get a good starting point for serum starved MCF10A cells, which do not undergo apoptosis or enter cell cycle without growth factor stimulation, in deterministic simulation mode.

#### Dependencies

1. `Docker`: All model dependencies and runtime environments are *Dockerized* into a downloadable image for self-contained model execution. To run the SPARCED model using Jupyter notebooks (SPARCED-jupyter), Docker must be installed. Then, by downloading the docker image, built on the Ubuntu-18.04 operating system with `python3` installed (hub.docker.com/repository/docker/birtwistlelab/sparced-notebook), users can run the Jupyter notebooks defined below in any web-browser within the docker container. The Docker image includes system utilities required for the simulation package `AMICI` (67,69).

### Jupyter notebook 1: model creation

The input files described above are processed by the “`createModel`” Jupyter notebook (68) (Supp. File 8 and converted into an Antimony (63) text file (Fig. 1B and Supp. File 9). This intermediate step and file provide an additional means to model input, fine-tuning, and alteration for experienced users. It can be explored via any text editor and it lists all elements of the model: species, rate laws, parameters, compartments, and corresponding values (Supp. File 9). This text file is then converted into an SBML (.xml) file, using `libantimony` in the same script. The Antimony format does not, to our knowledge, support addition of structured annotations, so the annotations (species and compartments) are added to the newly created SBML file using `libsbml` and the model input files (Supp. Files 2 and 6). Finally, the annotated SPARCED model file (.xml) is generated (Supp. File 10).

### Deterministic module

The model SBML file forms the basis of the deterministic module. When run deterministically, the model does not account for stochastic gene switching and mRNA transcription events (see next section). The default parameters and concentrations of the SPARCED model correspond to an average, serum-starved cell state in deterministic mode.

### Stochastic module

In addition to the deterministic module, the SPARCED model includes a stochastic module. The stochastic module describes gene states (active/inactive) and mRNA birth/death events for genes (currently 141 of them). The deterministic and stochastic modules exchange information every 30 simulated seconds. The current levels of select protein states can induce changes in gene activation/deactivation and/or mRNA transcription/decay rates. The newly updated mRNA copy numbers change nascent protein translation rates in the deterministic module. See (40) for further details.

The stochastic module constitutes two short Python scripts. At the start of each simulation, one of the scripts (`RunPrep.py`) reads in the `OmicsData` input file, processes parameter values, and sets the initial transcript levels and stochastic module rate constants. The second script (`SGEmodule.py`) uses information from `RunPrep.py` and species concentrations from the deterministic module to simulate mRNA transcription/degradation and gene activation/inactivation events. One output of the second script is the new concentrations of mRNAs, which is updated in the deterministic module to calculate rates of translation for the next 30 second simulation. The second output is the state of all gene copies (active or inactive).

### Jupyter notebook 2: model initialization

Making use of the created model file and the `Initializer` input file, total protein abundance data are converted protein and protein complex starting concentrations by adjusting translation rate constants. The “`InitializeModel`” notebook (Supp. File 16) also verifies that the simulated cells behave as expected in serum-starved state. It is possible to modify the steps of initialization to confer new basal behavior (such as cycling) or a mutational effect (loss of PTEN in U87 cells), as introduced by *Initialization* protocol in Bouhaddou2018 model. Running this file is optional and recommended only when new models are created for new cell contexts.

### Jupyter notebook 3: model simulation

Supp. File 11 includes an example of simulation setup and input parameters of the SPARCED model. The notebook “`runModel`” imports a user specified model SBML file and stochastic module scripts to run simulations using the `AMICI` package (69, 108). `AMICI` is an interface for `Sundials CVODEs` solvers that converts SBML files into executable C code for fast simulation. Other required input parameters for model simulation include: a flag to specify if the simulations are fully deterministic (flag=1) or hybrid (flag=0), the total simulation time in hours (th), input ligand concentrations, and a flag (1 or 0) to indicate if results should be exported.

### The SPARCED-I model

#### SPARCED-I model creation

We created a new enlarged version of the SPARCED model called SPARCED-I. We merged Yamada2003 model (73) of interferon-gamma receptor (IFNGR) signaling into SPARCED. The expansion included addition of 34 new species, 8 corresponding genes, and ∼70 reactions. New genes (IFNG, IFNGR, JAK2, STAT1, SHP2, SOCS1, PPN, PPX) corresponding to the proteins in the model are added as new rows and mRNA levels are inserted into the `OmicsData` input file. Gene copy numbers are taken as two (40). Each new protein, protein complex, and mRNA species are inserted into the `Species` and `StoichiometricMatrix` input files as new rows. Each new reaction is inserted into the `Ratelaws` input files as rows, and into the `StoichiometricMatrix` input file as new columns. The new genes are added as rows to the `GeneReg` file. The final SPARCED-I model has 954 species and 2540 reactions.

#### Parameter estimation of the SPARCED-I model

After expansion of the SPARCED model with IFNGR pathway and setting the initial species levels from MCF10A cells, SPARCED-I model showed ultrafast receptor and STAT1 activation dynamics inconsistent with biological observations. Thus, we selected six rate constant parameters for calibration based on substantial sensitivity for STAT1 activation dynamics. The parameters calibrated are: (i) STAT1 binding to activated receptor complexes, (ii) nuclear translocation rate of STAT1* dimers, (iii) SOSC1 mRNA translation rate constant, (iv) SOCS1 protein degradation rate, (v) STAT1 unbinding rate from active receptor-SOCS1 complexes, and set the EIF4E-dependent translation rate constant of SOCS1 to zero (see (40) for details on the EIF4E effect on translation). We fit the chosen parameters individually, while keeping the values of “best fit” at each step. The range of variation was set at ±2 in log10 scale. The SPARCED-I model was then run 1000 simulated hours, without any ligand stimulation, for equilibration.

#### Context estimation of the SPARCED-I model

The SPARCED model initial conditions are based on serum and growth factor starved MCF10A cells, grown in standard tissue culture plates. However, the experiments involving IFNγ were done in media with horse serum and collagen-coated tissue culture plates. The details of both experimental procedures are explained below. “Bridge” experiments showed that the horse serum upregulates active AKT (ppAKT) levels four-fold prior to growth factor stimulation. We modeled increased basal levels of active AKT with IGF1 treatment for 24 hours, and found that 0.02 nM IGF1 was consistent with the above experimental observations (Figs. S16 and S17). Therefore, we simulate the SPARCED-I model with 0.02 nM IGF1 treatment for 24 hours prior to EGF or EGF+IFNγ stimulation.

#### Cell proliferation estimation in SPARCED-I model

The Bouhaddou2018 model used cyclin A levels as a proxy for cell cycle entry in each cell (40). Here, we used the same MATLAB scripts (`findpeaks.m`) as the Bouhaddou2018 model to identify the peaks in cyclin A trajectories. A peak is defined as a hill-like shape, with a minimum prominence of 4 nM compared to surrounding basal levels. For each cell, the number of occurrences of such peaks are counted within 0-48 hours of simulation. Then, the peak counts in all 100 starting cells are summed to find the total number cell cycles in the population.

### SPARCED-nf (nextflow, model version running on Nautilus-Kubernetes cluster)

The SPARCED model can be ported to high-performance cloud computing infrastructures for large-scale simulation, in the SPARCED-nf variant. Specifically, we ran SPARCED-nf simulations using the Pacific Research Platform, a distributed network of academic computing resources organized as a Kubernetes cluster (64, 66). The prevalence of Kubernetes on both democratized and commercial cloud compute networks makes the model portable, allowing users to run large-scale jobs on distributed supercomputers on a wide range of platforms (65).

#### Dependencies

1. `Docker`: As is the practice with Kubernetes-compatible workflows, all model dependencies and runtime environments are *Dockerized* into a downloadable image for self-contained model execution. This means when a job for SPARCED-nf is launched on the Kubernetes cluster, it will download the Docker image for SPARCED-nf and execute the model within that container. The Docker image for SPARCED-nf is built on the Ubuntu-18.04 operating system with python3 installed, as well as a few minor system utilities required for `AMICI`. The image can be found at https://hub.docker.com/repository/docker/birtwistlelab/sparced.
2. `Nextflow` (nf): In this cloud-scalable version of the model, the Jupyter notebooks have been converted into python source code and re-modularized for greater parallel-simulation efficiency. The process of creating and executing the model is handled entirely by `Nextflow`, a workflow-management application and language for building resilient pipelines. When SPARCED-nf is launched, `Nextflow` begins by creating a head pod on the cluster to coordinate each of the jobs needed to run the model (Fig. S1). The head pod creates smaller jobs that each download the containerized dependencies from Dockerhub, pull the model source files from the SPARCED-nf GitHub repository, and run the assigned process. Once the model has completed execution, the output files are saved to a section of the Kubernetes cluster called the persistent volume claim (PVC), where they remain stored in the cloud for user download.

### SPARCED-nf model simulation set-up

SPARCED-nf uses the same tab-separated-value input files as SPARCED. For SPARCED-nf to build and execute, the files are copied into the aforementioned PVC for workflow access. This is done with `kube-runner` (https://github.com/SystemsGenetics/kube-runner), a submodule for automating common PVC tasks with Kubernetes’ `kubectl` tool. The `kube-load.sh` file is used to write new input to the PVC, and `kube-login.sh` is used to access and delete old input files from the cluster.

Along with its scalability, SPARCED-nf is also highly customizable. The `nextflow.config` configuration file is used to define the specifics of simulation scenarios.

1. `nextflow.config`: This configuration file has two main sections. In the first section (called K8), users define the Kubernetes namespace specifics and folder configurations. In the second section (called params), users customize runtime arguments for simulation settings. The available parameters are input_dir_name (the directory name of the input files), flag_deterministic (flag=1 for deterministic or flag=0 for hybrid simulations), sim_time (simulation time in hours), Vol_nuclear (volume of nuclear compartment in liters), Vol_cyto (volume of cytoplasmic compartment in liters), speciesVals (species names + initial concentration values to start from), ratelawVals (parameter names + values), and numCells (number of single cells if the simulations are hybrid). Importantly, the “speciesVals” and “ratelawVals” parameters allow users to pass in a formatted string to specify parameter sweeps. Using these in conjunction with the “numCells” parameter, the user can simulate thousands of cells in hundreds of different microenvironments in a single execution.
2. `SPARCED-nf:model_build`: Analogous to “`createModel.ipynb`” in SPARCED-jupyter model, this phase of the Nextflow pipeline constructs all necessary files for the model simulation.
3. `SPARCED-nf:split_from_params`: This is the major parallelizing step of SPARCED-nf. Having received the relevant model files from the last step, the workflow ingests the *speciesVals*, *ratelawVals*, and *numCells* arguments set by the user in the `nextflow.config`. Using the input files, it creates new input files to satisfy the user-specified parameter sweeps. Each new input file permutation is moved into its own new folder, and each such folder is duplicated *numCells* times.
4. `SPARCED-nf:model_run`: This final step of the Nextflow workflow is responsible for model execution and output generation. Each folder created in the previous step above serves as the unique runtime environment in this step. The model pulls assigned simulation input files associated with the folder. Each instance of this step is run in parallel across different simulation environments (Fig. S1B). Functionally, the code executed is very similar to the “`runModel.ipynb`” notebook and the model outputs are saved to the PVC.

When the models complete execution, each `SPARCED-nf:model_run` instance saves its output to a unique folder on the PVC. To download these folders to the local filesystem, users can employ `kube-save.sh` (from the `kube-runner` module).

### GitHub

The final model scripts, files, and information are available in Birtwistle Lab GitHub repository, <u>github.com/birtwistlelab/SPARCED</u>.

## Experimental Methods

### Cell culture and western blotting

*Cell culture:* MCF10A cells (acquired from LINCS Consortium/Gordon Mills and STR verified internally) are cultured in DMEM/F12 (Gibco #11330032) medium supplemented with 5% (by volume) horse serum (Gibco #16050122), 20ng/mL EGF (PeproTech #AF-100-15), 0.5 mg/mL hydrocortisone (Sigma #H-0888), 10μg/mL insulin (Sigma #I-1882), 100ng/mL cholera toxin (Sigma #C-8052), and 2mM L-Glutamine (Corning #25-005-CI). Cells were cultured at 37°C in 5% CO2 in a humidified incubator and passaged every 2-3 days with 0.25% trypsin (Corning #25-053-CI) to maintain subconfluency. Serum starvation medium is DMEM/F12 medium supplemented with 2mM L-Glutamine. Experimental starvation medium is DMEM/F12 medium supplemented with 5% (by volume) horse serum (Gibco #16050122), 0.5 mg/mL hydrocortisone (Sigma #H-0888), 100ng/mL cholera toxin (Sigma #C-8052), and 2mM L-Glutamine (Corning #25-005-CI).

*Tissue culture treated, non-collagen coated plates with full serum starvation:* The cells were seeded in full growth media at 150,000 cells/well in tissue culture treated six well plates (Corning # 08-772-1B). The next day, cells are washed once with 1X PBS (one phosphate buffered saline tablet (Sigma #P4417-100TAB) in 200 mL milli-Q water, autoclaved) and the media was exchanged to serum starvation media (DMEM/F12 medium, 2mM L-Glutamine) for 16-24 hours. Then, the cells were treated with vehicle control, EGF (10 ng/mL, PeproTech #AF-100-15), and HGF (40 ng/mL, R&D Systems #294-HGN-005) for 0, 5, and 60 minutes in a humidified, 5% CO2, 37°C incubator.

*Collagen-coated plates and growth-factor starvation only:* Collagen-coating mixture was prepared as follows: 7.5 mL diluent buffer (20% v/v glycerol, 10 mM EDTA, PBS), 1.5 mL Tris-HCL, 0.6 mL COL1 (Cultrex #3442-050-01), and 5.4 mL PBS. 950 μL coating mix was added into each well of a six-well plate. After making sure that the entire well surface was covered, the plates were incubated one hour at room temperature. After incubation, any remaining liquid is aspirated and discarded. The wells were washed twice with sterile PBS and left lid-open under a sterile laminar flow hood until wells were fully dry (∼one hour). Upon replacement of the plate lid, the plates were stored in a benchtop desiccator at room temperature for a minimum of 3 days before use. Then, MCF10A cells were seeded in full growth media at 150,000 cells/well. After being allowed to attach for 7-8 hours, the wells were washed once with PBS and the media was changed to full growth media without EGF and insulin for 18 hours. Then, the cells were treated with vehicle control, EGF (10 ng/mL), and HGF (40 ng/mL) as above for 0, 5, and 60 minutes.

*Cell lysis:* After growth factor treatment, the plates were removed from the incubator and put on ice. The media in the wells were aspirated and the wells were washed with PBS. 110 μL of freshly-prepared, ice-cold RIPA buffer (50mM Tris, pH 7-8 (Acros Organics #14050-0010), 150 mM NaCl (Fluka #71383), 0.1 % SDS (Fisher #46040CI), 0.5% sodium deoxycholate, 1% Triton-X-100, filter sterilized, stored at 4C) with protease & phosphatase inhibitors (1μg/mL aprotinin, 1μg/mL leupeptin, 1μg/mL pepstatin A, 10 mM β-glycerophosphate, and 1mM sodium orthovanadate) was added into each well, while gently rotating the plate to cover the full surface area. The plates were transferred to the cold room for 15-20 minutes, with slow rocking. The lysate was scraped off from the wells with a cell scraper. 100 μL of cell lysate from each well were transferred into labeled Eppendorf tubes on ice. Each tube was vortexed three times to homogenize cell debris, keeping other tubes on ice. All tubes were then centrifuged at 4°C for 15 minutes at 14000 rpm. 80 μL of the supernatant from each tube was transferred into new Eppendorf tubes on ice. These cleared lysate samples were stored at −80C for long-term storage or used immediately as below.

*Protein quantification:* Total protein quantification was done using the BCA-Pierce 660 Assay (Thermo Scientific #23225). As the reference, BSA stock (Thermo Scientific #23209) was used according to the manufacturer protocol. In short, 10 μL of samples and BSA standards were loaded into wells, in triplicate, in a 96-well plate (Corning #3370). 150 μL BCA Protein Assay Reagent was loaded into each non-empty well. The plate was covered and incubated at room temperature for 5 minutes. The absorbance at 660 nm was measured on a plate reader (BioTek #Epoch2). The average reading of blank wells was subtracted from all other readings, and then average readings were calculated. The standard curve was fitted by a polynomial using blank-corrected mean values of each standard condition versus its BSA concentration. The fitted curve was used to determine the protein concentration in each sample.

*Immunoblotting and quantification:* The lysates were put on ice and the amount of sample to load into each well was calculated using the total protein concentrations determined above (3 μq loaded here). Each sample was mixed with 2X Sample Buffer (950 μL of Laemmli’s Buffer (BioRad #161-0737), 50 μL beta-mercaptoethanol (Fisher #03446I-100)) in a 1:1 ratio and transferred into a new Eppendorf tube. The sample solutions were heated at 95°C for 5min on the heating plate and then briefly spun in benchtop microcentrifuge to return any condensation to the bottom of the tube. 10% acrylamide gels were prepared, and samples were loaded into the wells. A pre-stained protein ladder (LI-COR #928-70000) was loaded in the first and last wells. The gel was run in SDS running buffer (100 mL 10X Tris-Glycine-SDS buffer (IBI Scientific #IB01160) + 900 mL milli-Q water) at constant 220V until the dye front runs off the gel. Then, wet-transfer to nitrocellulose membrane (0.45µm pore size, VWR #10063-173) was done using cold transfer buffer (3.03g Tris-base (Acros Organics #14050-0010), 14.4g glycine (Acros Organics #220910050), 100 mL methanol (BDH #BDH1135-4LP), volume adjusted to 1 Liter with milli-Q water) and running the cassette with ice block at constant 100V for one hour. 1X TBST (Tris-Buffered Saline, 0.1% Tween) was prepared: 100 mL of 10X TBS solution (24 g Tris-base, 88 g NaCl (Fluka #71383), adjusted pH to 7.6, adjusted final volume to 1 Liter with milli-Q water, autoclaved), 1 mL Tween-20 (Fisher #BP337-100), and milli-Q water until final volume of 1 Liter (∼900 mL). When the transfer was finished, the membrane was blocked using BSA-TBST blocking buffer (2.5 g bovine serum albumin (Fisher# BP1600-100), 50 mL 1X TBST) for 45min at room temperature. The blocking buffer was discarded, and the membrane is incubated in primary antibody solution (1:1000 dilution, 10 μL primary antibody in 10 mL 5% BSA-TBST blocking buffer) overnight in cold room. The primary antibodies used were: AKT_pS473 (Cell Signaling #4060; 1:1000), AKT (Cell Signaling #2920; 1:1000), ERK_pT202_pY204 (Cell Signaling #4370; 1:1000), ERK (Cell Signaling #4696; 1:1000), alpha-tubulin (Novus #NB100-690, 1:1000), and β-actin (LI-COR #926-42212, 1:1000). After primary antibody incubation, membranes were washed three times for 15 min each with 1X TBST at room temperature, with gentle rocking. Then, the membranes are incubated with LI-COR secondary antibodies in 10 ml TBST blocking buffer for 45 minutes (anti-rabbit 800CW, LI-COR #926-32211 or anti-mouse 680LT, LI-COR #925-68070; 1:8000) at room temperature, with gentle rocking. Membranes was washed three times for 15 min each, with 1X TBST, on rocker. The imaging was done with a LI-COR Odyssey Infrared Imager, where bands were quantified using LI-COR Image Studio Lite v5.2 software (Figs. S16-19).

### Data availability

The LINCS datasets analyzed during the current study are available in the Synapse repository, synapse.org/#!Synapse:syn12526172. All other data are available from the authors upon reasonable request.

## ACKNOWLEDGMENTS

The authors acknowledge funding from the National Institutes of Health Grants R01GM104184 and U54HG008098-LINCS Center (MRB), U54CA209988 and U54HG008100-LINCS Center (LMH), the National Science Foundation Grant CC*-1659300 (FAF), and portions of this work were performed under the auspices of the U.S. Department of Energy by Lawrence Livermore National Laboratory under Contract DE-AC52-07NA27344 (RCB). CE was an NIH-LINCS Postdoctoral Fellow.

## AUTHOR CONTRIBUTIONS

Conceptualization: CE, MRB

Methodology: CE, LMH, FAF, MRB

Software: CE, EMB, AM, MMS, MB, RCB, WD

Validation: CE, EMB, AM, RCB, MRB

Formal Analysis: CE

Investigation: CE, SMG, LMH

Resources: LMH, FAF, MRB

Data Curation: CE, AM, WD, SMG

Writing – Original Draft: CE, MRB

Writing – Review & Editing: CE, MRB

Visualization: CE, EMB, MB

Supervision: CE, MRB

Project Administration: CE, MRB

Funding Acquisition: LMH, FAF, MRB

## SUPPLEMENTAL INFORMATION

**Supplemental Figure 1.** (A) SPARCED-nf implementation overview. (B) Details of SPARCED-nf pipeline.

**Supplemental Figure 2.** SPARCED model includes a stochastic gene expression module. Two isoforms of ERK gene (MAPK1 and MAPK3) are activated randomly (A) and leads to two distinct mRNAs (B). The ERK1 and ERK2 mRNAs are translated into a single ERK protein (C). The trajectories are obtained from a stochastic single cell simulation with no growth factor stimulation for 24 hours.

**Supplemental Figure 3.** (A) Ligand-receptor binding and Hill coefficients for each pair in MCF10A context. The simulations capture literature knowledge. (B) The dynamics of activated EGFR (membrane-bound and internalized) dimers are recaptured by the SPARCED model, compared to Bouhaddou2018 model.

**Supplemental Figure 4.** Signaling dynamics of ppERK, ppAKT, and pEIF4EBP1 induced by EGF, Insulin, or EGF+Insulin treatment for 6 hours. Serum-starved MCF10A cells are stimulated with EGF (0.01, 0.1, 1, and 10 nM), Insulin (0.17, 1.7, 17, and 1721 nM), or EGF+Insulin (0.01+0.17, 10+0.17, 0.01+1721, and 10+1721 nM).

**Supplemental Figure 5.** Signaling dynamics of ppERK and ppAKT induced by EGF, Heregulin (NRG1), HGF, PDGF, FGF, IGF, and Insulin treatment for 2 hours. Serum-starved MCF10A cells are stimulated with corresponding ligands at a dose range of 0.001 to 1000 nM.

**Supplemental Figure 6.** (A) p53 is activated in response to double (middle) or/and single (top/bottom) stranded DNA break damage. When DNA break repair mechanism is turned on (orange curves), p53 activity (or oscillatory behavior) dies down. (B) Single cells show different levels of p53 response to DNA damage. Increasing DNA damage amount (top to bottom) leads to higher number of activated p53 peaks. (C) the number of p53 pulses increases with increasing DNA damage, whereas pulse height and width remain relatively constant (results based on simulations shown in B). Plots show mean ± s.e.m.

**Supplemental Figure 7.** (A) Increasing TRAIL dose decreases the time it takes to die (ttd) for the average cell. Representative cells trajectories are shown, where the cells are simulated deterministically with different doses of TRAIL until they die (or up to 100 hours). The time of death is defined by the amount of cleaved PARP (cPARP, y-axis) surpassing the amount un-cleaved PARP. (B) Summary of ttd values for different TRAIL doses. (C) The fraction of surviving cells decreases as stimulated TRAIL dose increases. The red circles represent percentage of living cells when 20 stochastic single cells are simulated with specified TRAIL dosage for 5 hours. The black stars are experimental data from Bouhaddou 2018 model (D) Increasing ERK and AKT activity levels prolongs TRAIL induced time to death (blue curve), whereas increasing PUMA and NOXA expression levels decreases the time it takes for cells to die (red curve). Cells with specified alterations are compared to the cell stimulated with a low dose of TRAIL (black curve). cPARP levels are the proxy for cell death, where the cells go apoptosis when [cPARP]>[PARP].

**Supplemental Figure 8.** (A) Increasing Cyclin D mRNA levels induces proper cyclin-CDK complex progression and oscillations for cell cycle entry. Plots show Cyclin D, E, A, and B concentrations when basal (blue), 10X basal (dark orange), and 60X basal (light orange) levels of Cyclin D mRNA (CYCD) are simulated. (B) The number of ribosomes in the cell doubles around 20 hours. The cell is simulated with full growth condition (EGF=100 nM, NRG1=100 nM, HGF=100 nM, PDGF=100 nM, FGF=100 nM, IGF=100 nM, INS=100 nM).

**Supplemental Figure 9.** Inhibition of AKT and ERK pathways together synergistically increase cell death, in EGF and insulin stimulated cells. Serum-starved MCF10A cells are simulated with following conditions: (A) No stimulation, (B) EGF=20ng/mL + Insulin=10μg/mL, (C) EGF=20ng/mL + Insulin=10μg/mL + MEKi=10μM, (D) EGF=20ng/mL + Insulin=10μg/mL + AKTi=10μM, and (E) EGF=20ng/mL + Insulin=10μg/mL + MEKi=10μM + AKTi=10μM for up to 80 hours. The bar plots show mean ± s.e.m. of time to death for 30 cells. The ttd are captured by cPARP spikes.

**Supplemental Figure 10.** (A) Simulations where BIM-dependent or BAD-dependent mechanisms are switched off and percent death calculated in response to EGF + insulin at 48 hours. The results show that ERK and AKT inhibition induced cell death mechanisms are mostly BIM dependent, not BAD. Bars represent mean ± s.e.m. of 100 stochastic cell simulations. (B) EGF and insulin cooperatively induce cell cycle entry, with insulin inducing very little cell cycle entry alone. Cells are simulated with EGF (10nM), Insulin (1721nM), or EGF+Insulin (10nM+1721nM) for 30 hours and the percentage of cells entering S-phase are calculated. Cells are considered in S-phase when the sum of concentrations of Cyclin E, A, and B is greater than 20nM. Bars represent mean ± s.e.m. of 100 stochastic cell simulations.

**Supplemental Figure 11.** Activation of both ERK and AKT pathways are required for robust cell cycle entry. Time averaged ppERK and ppAKT levels correlate with Cyclin D levels. Basal levels of ppERK and ppAKT are increased (between 1X-20X) and each condition is simulated up to 6 hours. The time-averaged levels of ppERK and ppAKT are plotted against the time-averaged Cyclin D levels. Conditions representing EGF (10nM), Insulin (1721nM), and EGF+Insulin (10nM+1721nM) are shown with colored circles.

**Supplemental Figure 12.** SPARCED model recapitulates downstream pathway activation by ligands and ligand combination treatments. Experimental data and simulation results from MATLAB (lines) and SPARCED (circles) models with EGF (top) and EGF+Insulin (bottom) stimulation for 6 hours. Plots show double-phosphorylated ERK (ppERK), serine-phosphorylated AKT (pAKT), and phospho-EIF4EBP1 (pEIF4EBP1) levels. The numbers in gray shaded boxes represents numbering of conditions in Fig. 3A. Exp: Experimental data, Sim: Simulation.

**Supplemental Figure 13.** (A) Bar plots corresponding to the conditions shown in Fig. 2C. Gray bars are experimental or simulation data from Bouhaddou 2018 model and blue bars are simulation results of SPARCED model. Bars represent mean ± s.e.m. (B) Etoposide treatment alone induces lesser cell death compared to Etoposide + Growth Factor stimulation, shown in Fig. 3D-E. (C) Percentage of cell death of 100 cells shown in (B). Bars represent mean ± s.e.m.

**Supplemental Figure 14.** SPARCED model alteration guidelines. (A) Steps of model expansion and context change procedures are listed. Refer to Supp. File 13 for more details. Steps can be skipped if no changes are necessary. (B) The list of parameters and species values modified for SPARCED model context change from MCF10A cells to U87 cells. (C) SPARCED_U87 model simulations reproduce previous observations, where U87 cells show increased response and sensitivity to AKT inhibition. MEKi: MEK inhibitor, AKTi: AKT inhibitor. Bars represent mean ± s.e.m. of 100 single cell simulations for each condition.

**Supplemental Figure 15.** Comparison of initialized species concentrations for MCF10A (top) and U87 (bottom) cells. The Bouhaddou2018 model values are reproduced by the new initialization notebook. The concentrations values (black dots) are almost exact and identity line (dashed red) coincides with the linear fit line (solid black). SPARCED initialized values are on y-axis and B2018 initialized values are on x-axis.

**Supplemental Figure 16.** (A) The experimental setup and (B) results of exploring the effect of collagen-coating and serum starvation in MCF10A cells.

**Supplemental Figure 17.** (A) The quantification of western blots shown in Fig. S16. All data are normalized to condition 10 (cells in non-coated plates with full growth media, measurements at time 0 hr). At least three biological replicates. pAKT levels in growth factor starved cells in collagen-coated plates (#3) are four times higher than pAKT levels in serum+growth factor starved cells in non-coated plates (#11). No significant change in pERK levels between two conditions (#3 and #11). The order of bars is given according to the list on the left in panel (B). (B) The experimental conditions of western blot membranes in Figs. S16, S18, and S19.

**Supplemental Figure 18.** The full images of western blots (replicates 1 and 2) shown in Fig. S16.

**Supplemental Figure 19.** The full images of western blots (replicates 3 and 4) shown in Fig. S16.

**Supplemental Figure 20.** (A) p21 trajectories of SPARCED-I Mechanism 1 model simulations. Changing STAT1 regulation of p21 transcription induces higher p21 levels (orange, right) compared to original (orange, left) simulations. The p21 level increase does not occur in single EGF stimulation conditions (blue). (B) The total number of cyclin A peaks of 100 starting cells at 48 hours in SPARCED-I mech1 (gray) and modified SPARCED-I mech1 (orange) models.

**Supplemental Figure 21.** (A-B) Normalized pMAPK levels show a significant change when IFNγ is included in addition to the EGF, in both Mechanism 1 (A) and Mechanism 2 (B). (C) Normalized ppAKT levels do not show a significant decrease after IFNγ treatment, when EGF+IFNγ Mechanism 1 simulations are compared to EGF alone case. (D) Normalized ppAKT levels show a significant decrease after IFNγ treatment, when EGF+IFNγ Mechanism 2 simulations are compared to EGF alone case. RPPA data are shown in black error lines, from three independent replicates. Colored dark lines represent median cell trajectories from simulations, dark and light-colored regions represent 70th and 95th quantiles, respectively. Experimental data are from synapse.org/#!Synapse:syn12526172.

**Supplemental File 1.** OmicsData input file.

**Supplemental File 2.** Species input file.

**Supplemental File 3.** Ratelaws input file.

**Supplemental File 4.** Stoichiometric Matrix input file.

**Supplemental File 5.** Gene Regulation input file.

**Supplemental File 6.** Compartments input file.

**Supplemental File 7.** Observables input file.

**Supplemental File 8.** Model creation Jupyter notebook.

**Supplemental File 9.** SPARCED model Antimony file.

**Supplemental File 10.** SPARCED model SBML file.

**Supplemental File 11.** Model import and simulation Jupyter notebook.

**Supplemental File 12.** Parameter value replacements for U87 cell line SPARCED model.

**Supplemental File 13.** SPARCED model alteration steps.

**Supplemental File 14.** U87 cell line omics data and model parameter values from the Bouhaddou2018 model.

**Supplemental File 15.** Protein-to-mRNA ratios file.

**Supplemental File 16.** Model initialization Jupyter notebook.

**Supplemental File 17.** Model initialization Jupyter notebook for U87 cell line.

**Supplemental File 18.** Initialization input file.

**Supplemental File 19.** U87 cell line species input file.

**Supplemental File 20.** Gene Regulation input file for SPARCED-I model.

**Supplemental File 21.** Gene Regulation input file for SPARCED-I Mechanism 1 model.

**Supplemental File 22.** MCF10A RNA-seq and proteomics data.

**Supplemental File 23.** SPARCED-I model SBML file.

**Supplemental File 24.** SPARCED-I Mechanism 1 model SBML file.

**Supplemental File 25.** SPARCED-I Mechanism 2 model SBML file.

