## Supplemental Files and Figures for "A Scalable, Open-Source Implementation of a Large-Scale Mechanistic Model for Single Cell Proliferation and Death Signaling": SPARCEDpaperF_SuppFigs.pdf

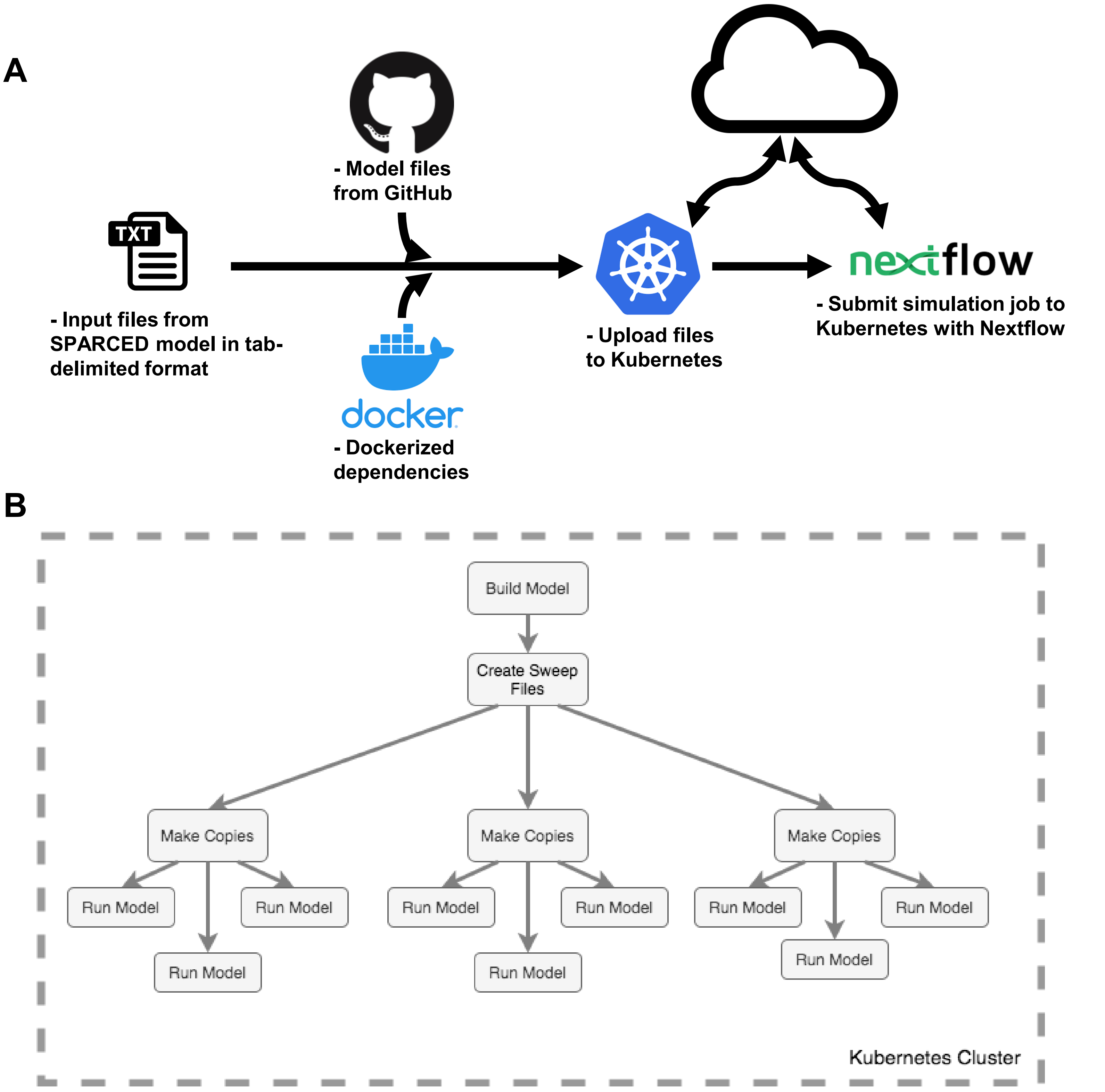


**Figure S1.** (A) SPARCED-nf implementation overview. (B) Details of SPARCED-nf pipeline.

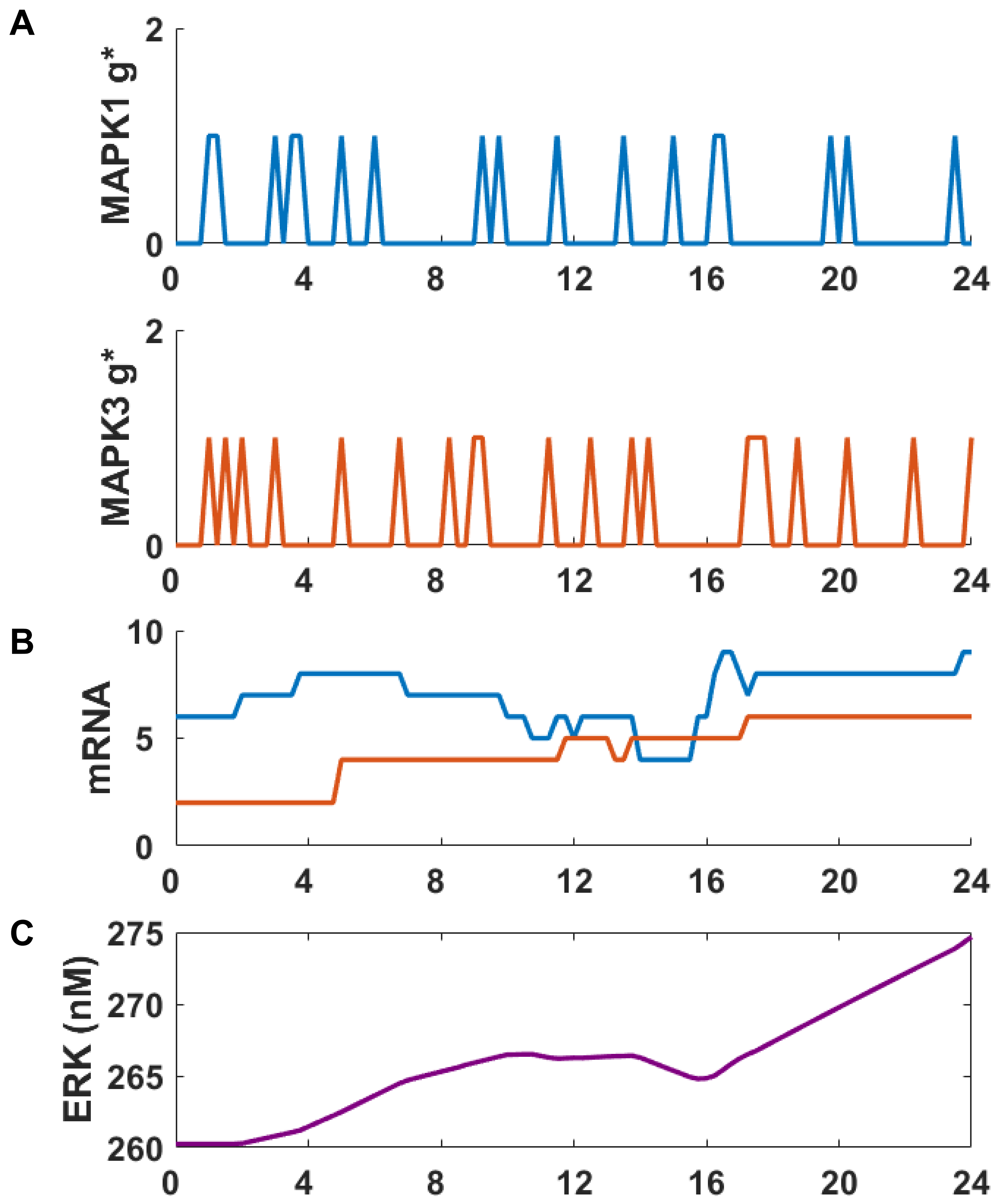


**Figure S2.** SPARCED model includes a stochastic gene expression module. Two isoforms of ERK gene (MAPK1 and MAPK3) are activated randomly (A) and leads to two distinct mRNAs (B). The ERK1 and ERK2 mRNAs are translated into a single ERK protein (C). The trajectories are obtained from a stochastic single cell simulation with no growth factor stimulation for 24 hours.

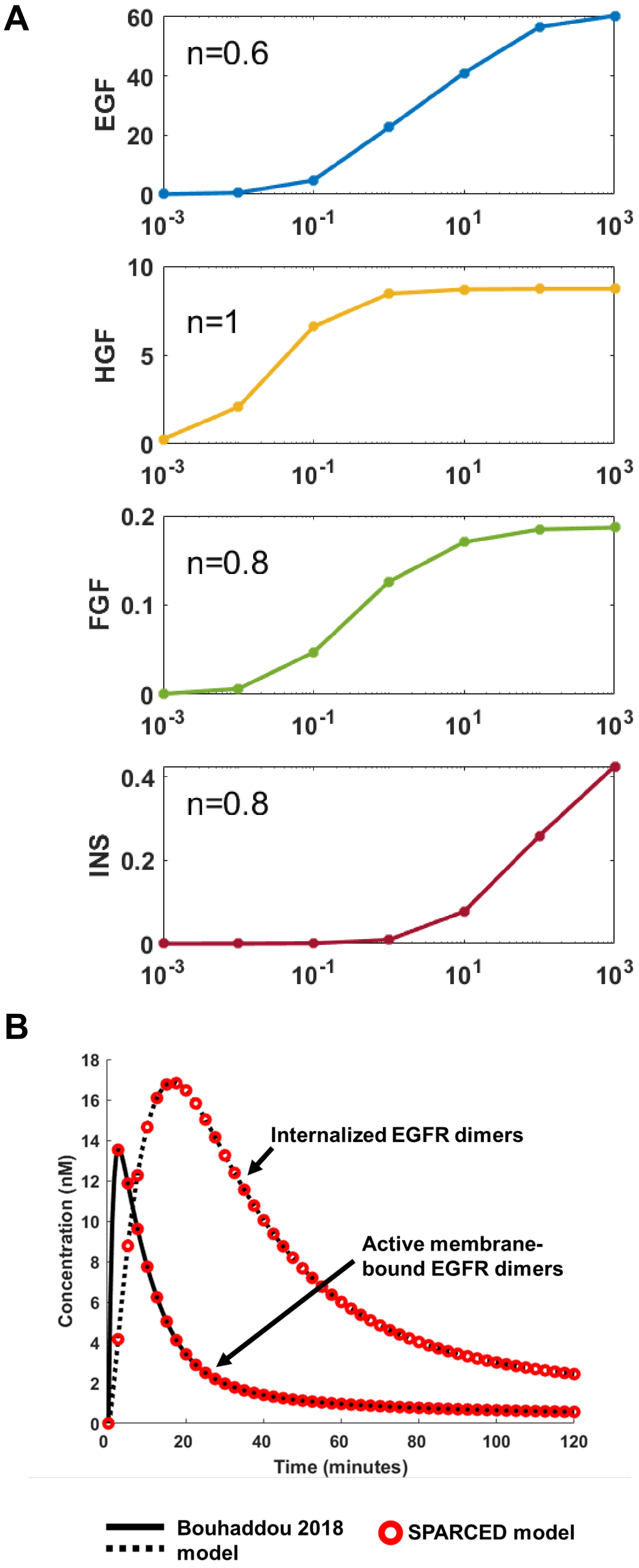


**Figure S3.** (A) Ligand-receptor binding and Hill coefficients for each pair in MCF10A context. The simulations capture literature knowledge. (B) The dynamics of activated EGFR (membrane-bound and internalized) dimers are recaptured by the SPARCED model, compared to Bouhaddou2018 model.

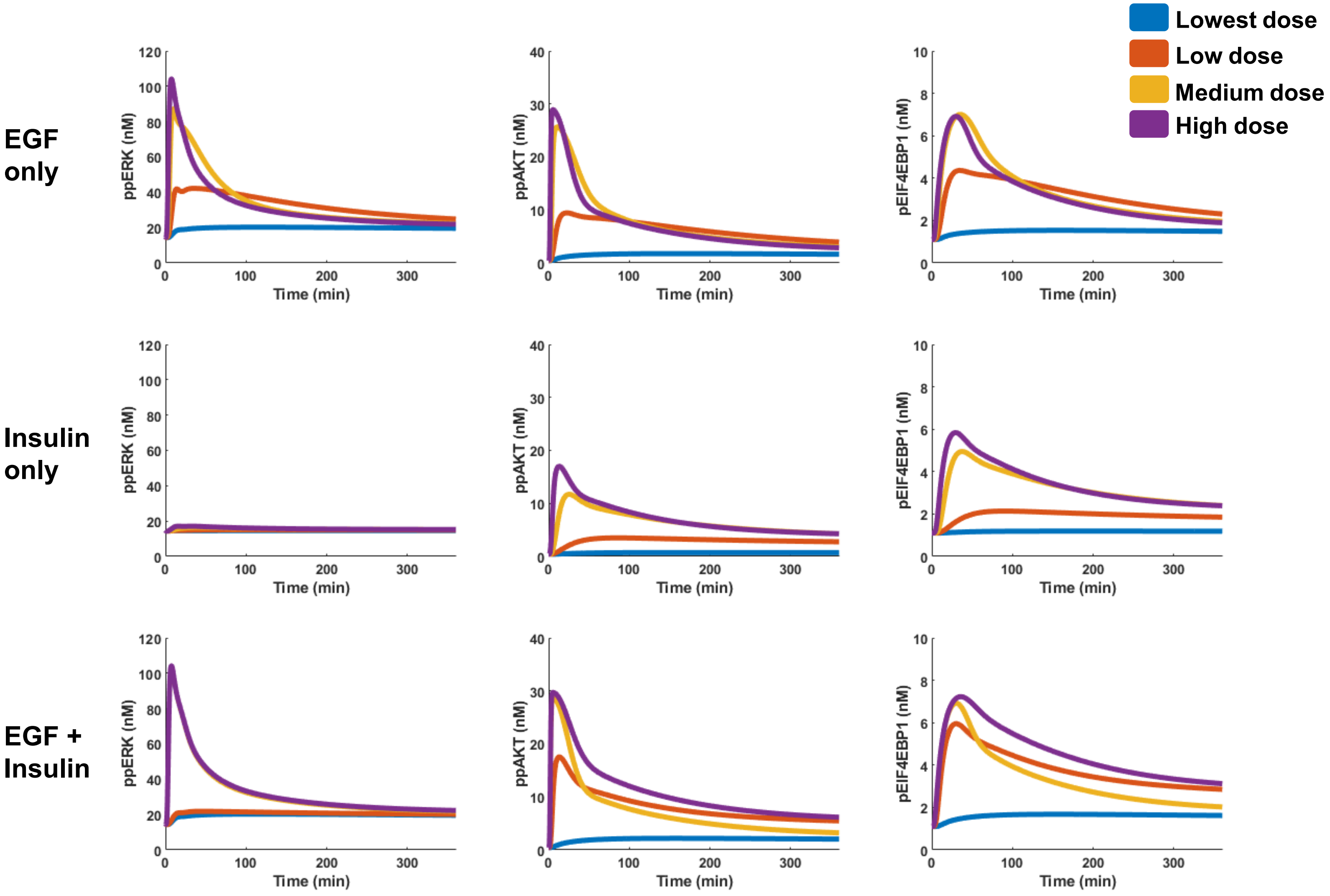


**Figure S4.** Signaling dynamics of ppERK, ppAKT, and pEIF4EBP1 induced by EGF, Insulin, or EGF+Insulin treatment for 6 hours. Serum-starved MCF10A cells are stimulated with EGF (0.01, 0.1, 1, and 10 nM), Insulin (0.17, 1.7, 17, and 1721 nM), or EGF+Insulin (0.01+0.17, 10+0.17, 0.01+1721, and 10+1721 nM).

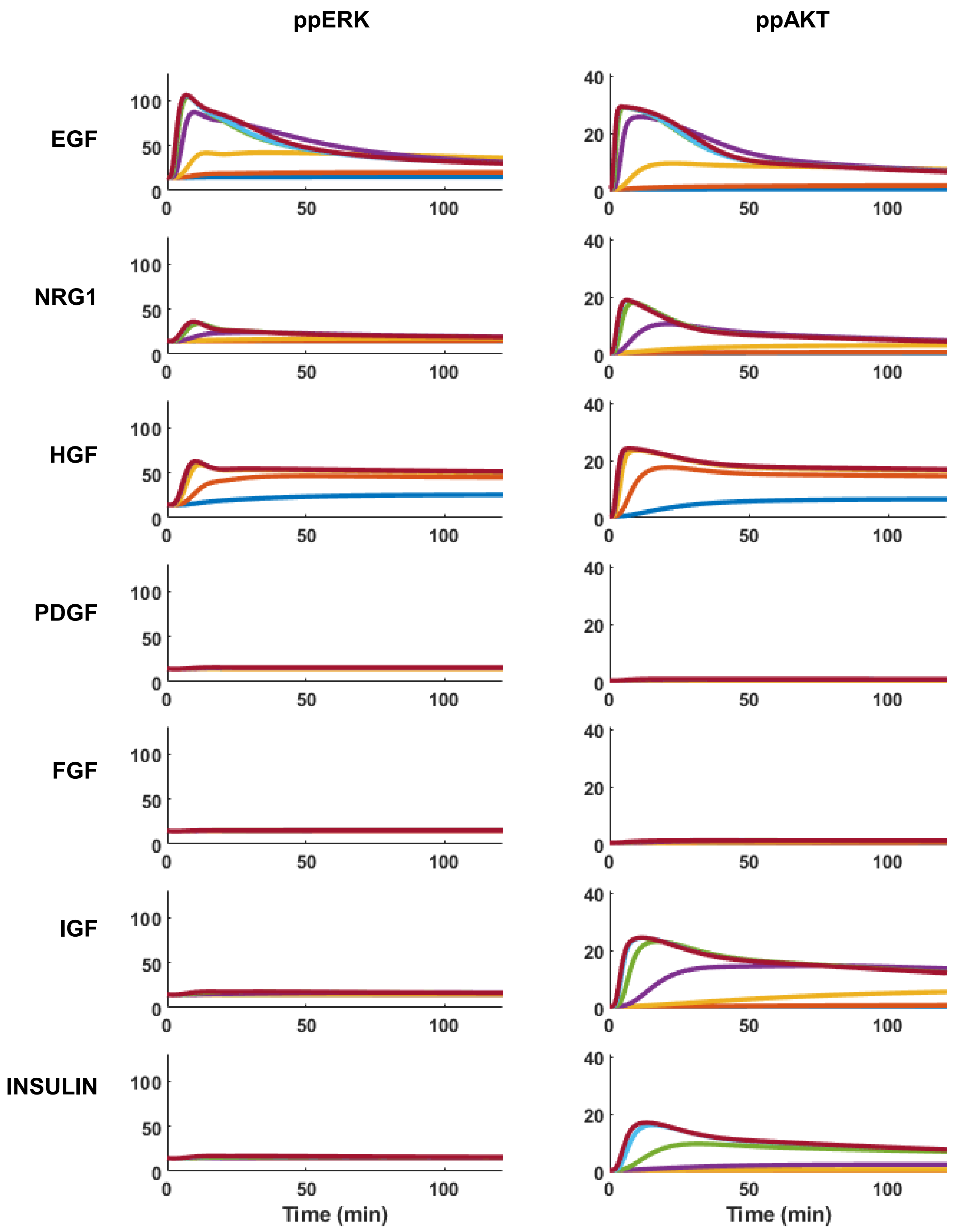


**Figure S5.** Signaling dynamics of ppERK and ppAKT induced by EGF, Heregulin (NRG1), HGF, PDGF, FGF, IGF, and Insulin treatment for 2 hours. Serum-starved MCF10A cells are stimulated with corresponding ligands at a dose range of 0.001 to 1000 nM.

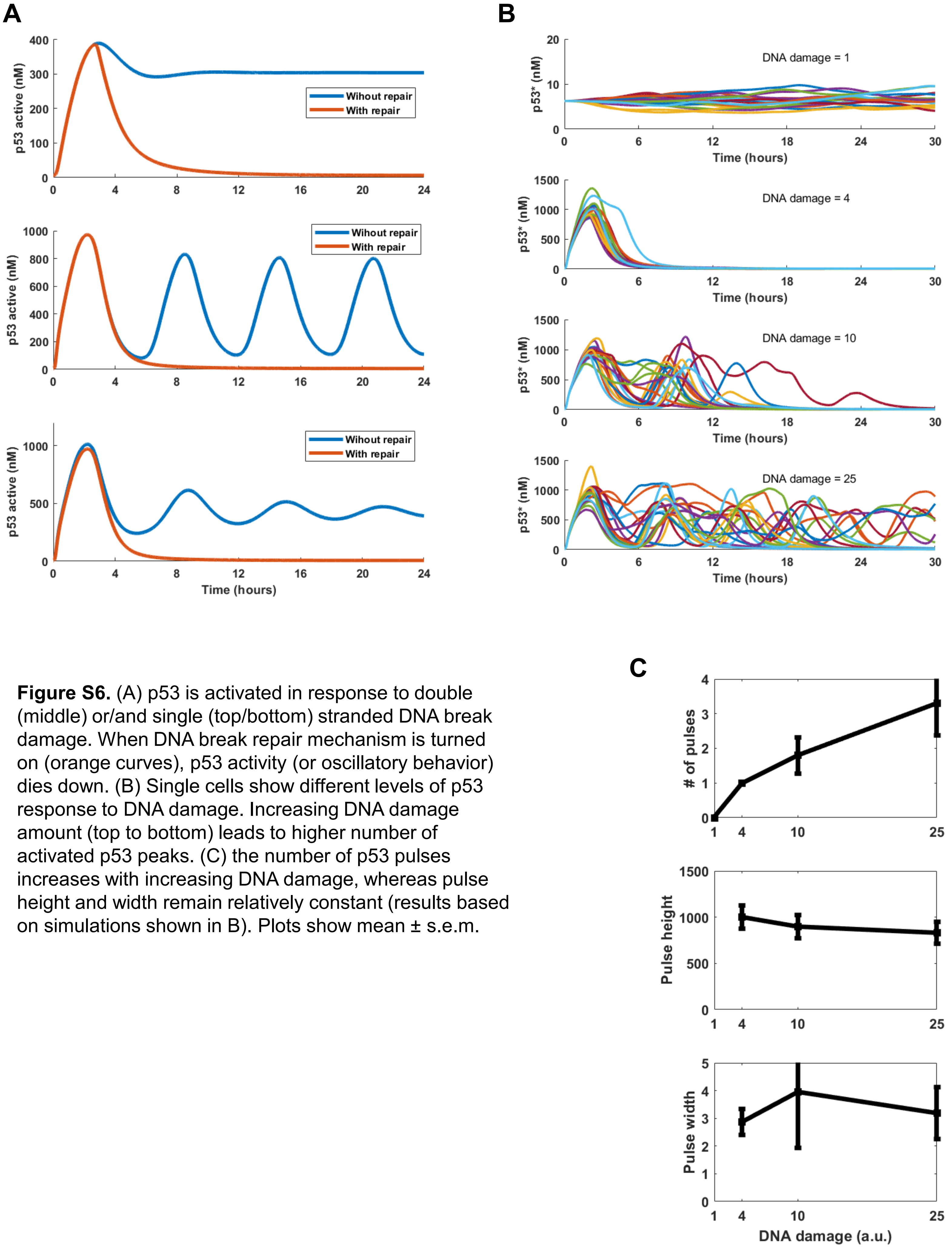


**Figure S6.** (A) p53 is activated in response to double (middle) or/and single (top/bottom) stranded DNA break damage. When DNA break repair mechanism is turned on (orange curves), p53 activity (or oscillatory behavior) dies down. (B) Single cells show different levels of p53 response to DNA damage. Increasing DNA damage amount (top to bottom) leads to higher number of activated p53 peaks. (C) the number of p53 pulses increases with increasing DNA damage, whereas pulse height and width remain relatively constant (results based on simulations shown in B). Plots show mean ± s.e.m.

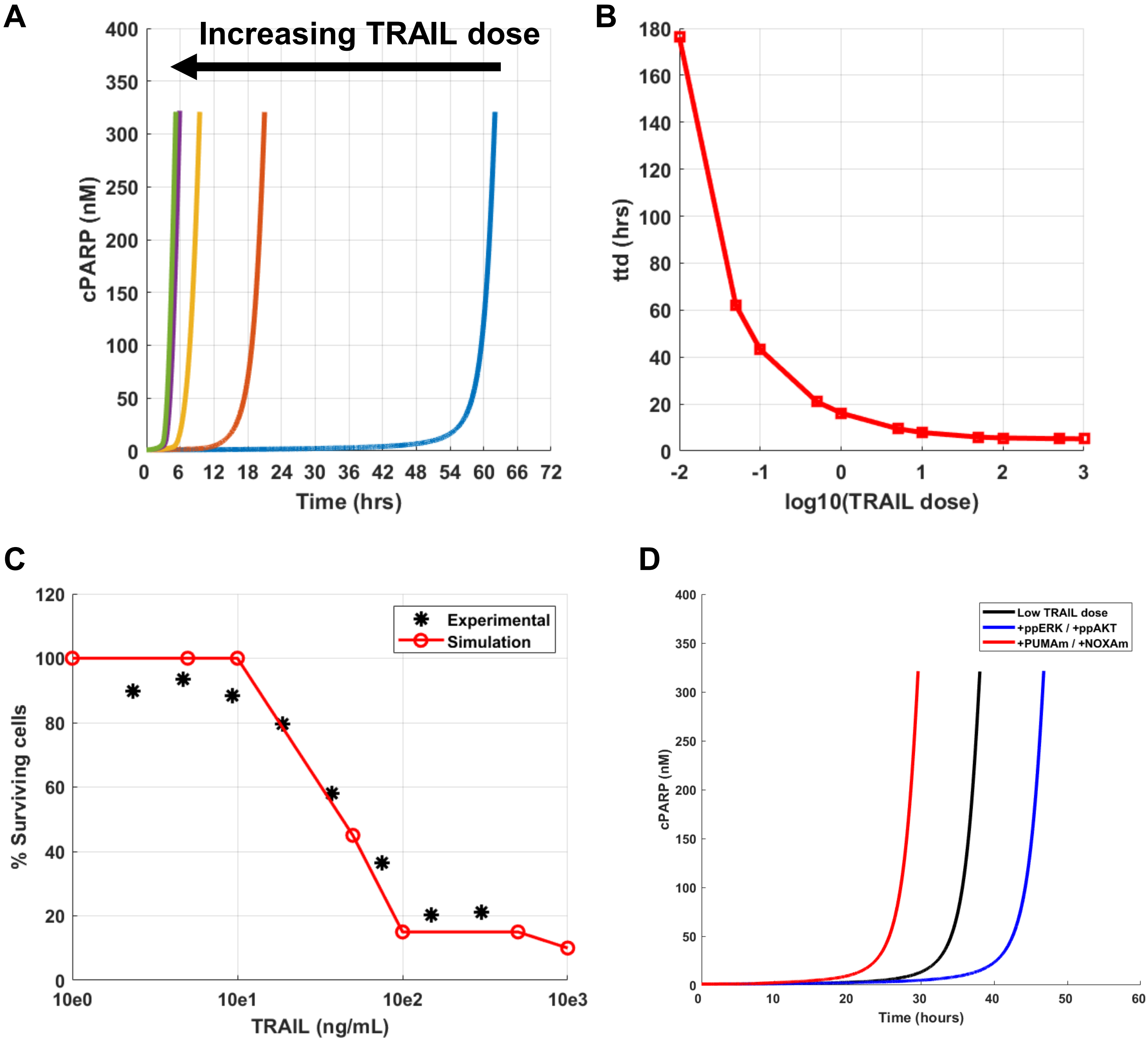


**Figure S7.** (A) Increasing TRAIL dose decreases the time it takes to die (ttd) for the average cell. Representative cells trajectories are shown, where the cells are simulated deterministically with different doses of TRAIL until they die (or up to 100 hours). The time of death is defined by the amount of cleaved PARP (cPARP, y-axis) surpassing the amount un-cleaved PARP. (B) Summary of ttd values for different TRAIL doses. (C) The fraction of surviving cells decreases as stimulated TRAIL dose increases. The red circles represent percentage of living cells when 20 stochastic single cells are simulated with specified TRAIL dosage for 5 hours. The black stars are experimental data from Bouhaddou 2018 model (D) Increasing ERK and AKT activity levels prolongs TRAIL induced time to death (blue curve), whereas increasing PUMA and NOXA expression levels decreases the time it takes for cells to die (red curve). Cells with specified alterations are compared to the cell stimulated with a low dose of TRAIL (black curve). cPARP levels are the proxy for cell death, where the cells go apoptosis when [cPARP]>[PARP].

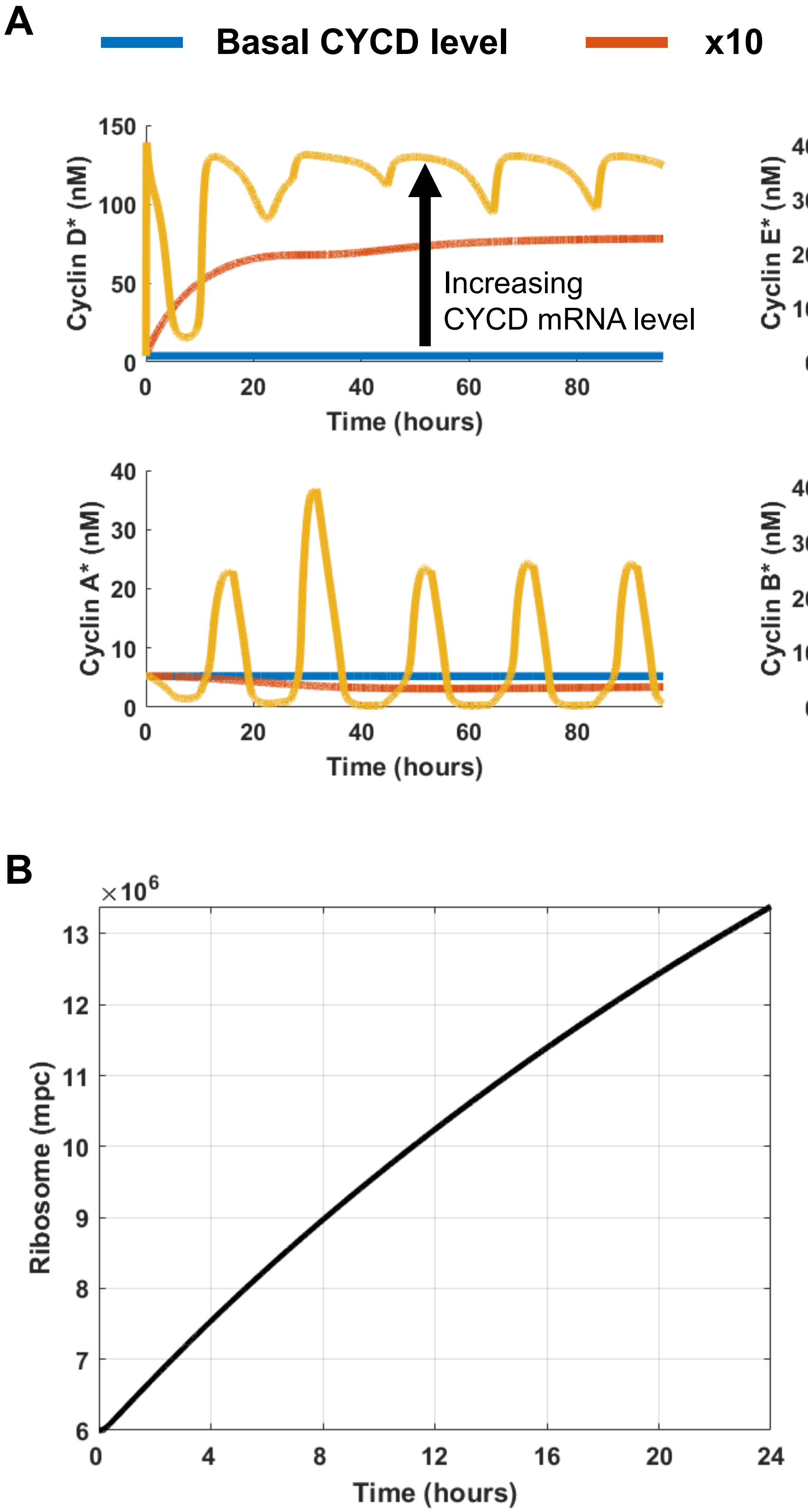


**Figure S8.** (A) Increasing Cyclin D mRNA levels induces proper cyclin-CDK complex progression and oscillations for cell cycle entry. Plots show Cyclin D, E, A, and B concentrations when basal (blue), 10X basal (dark orange), and 60X basal (light orange) levels of Cyclin D mRNA (CYCD) are simulated. (B) The number of ribosomes in the cell doubles around 20 hours. The cell is simulated with full growth condition (EGF=100 nM, NRG1=100 nM, HGF=100 nM, PDGF=100 nM, FGF=100 nM, IGF=100 nM, INS=100 nM).

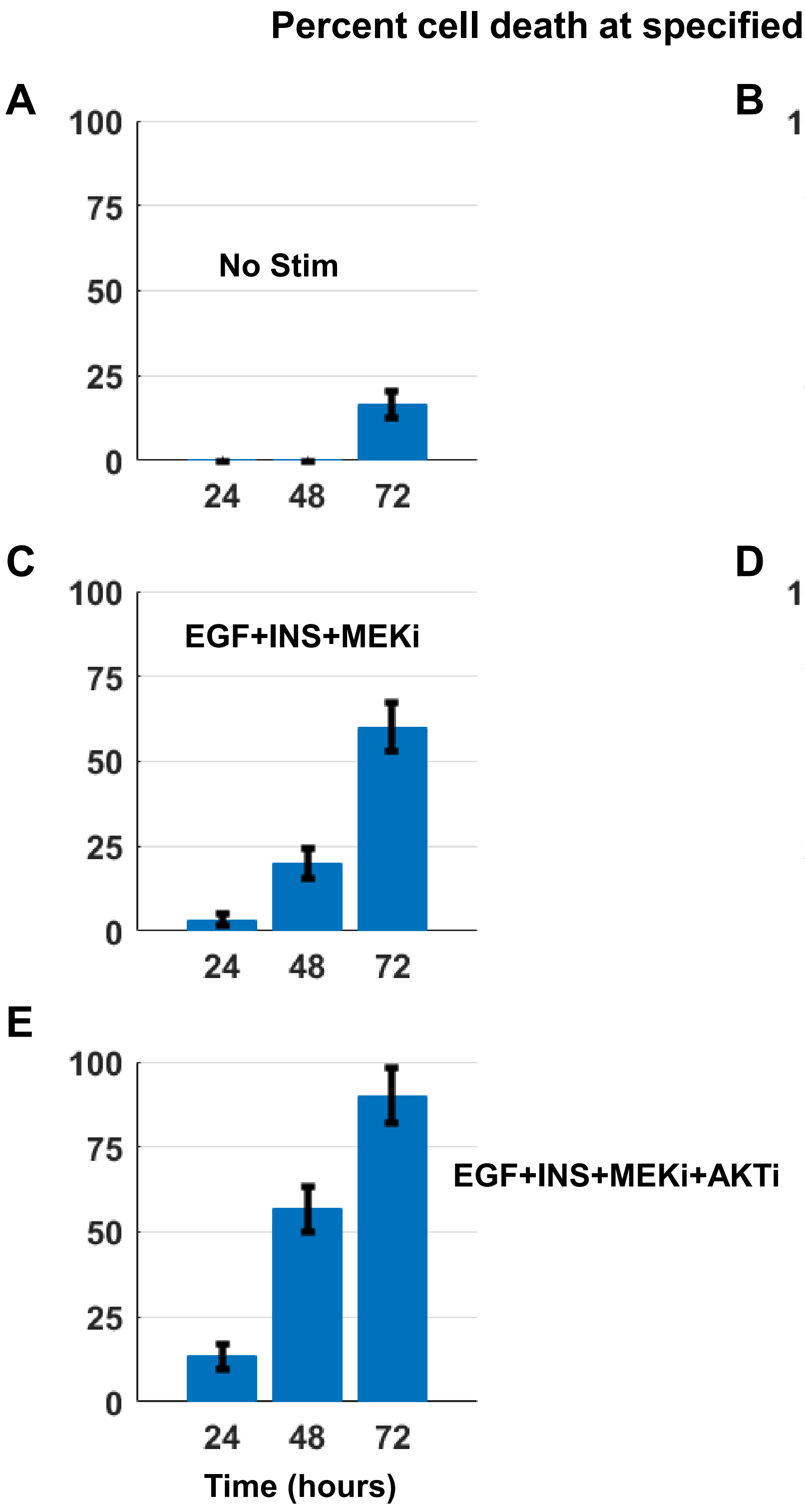


**Figure S9.** Inhibition of AKT and ERK pathways together synergistically increase cell death, in EGF and insulin stimulated cells. Serum-starved MCF10A cells are simulated with following conditions: (A) No stimulation, (B) EGF=20ng/mL + Insulin=10µg/mL, (C) EGF=20ng/mL + Insulin=10µg/mL + MEKi=10µM, (D) EGF=20ng/mL + Insulin=10µg/mL + AKTi=10µM, and (E) EGF=20ng/mL + Insulin=10µg/mL + MEKi=10µM + AKTi=10µM for up to 80 hours. The bar plots show mean ± s.e.m. of time to death for 30 cells. The ttd are captured by cPARP spikes.

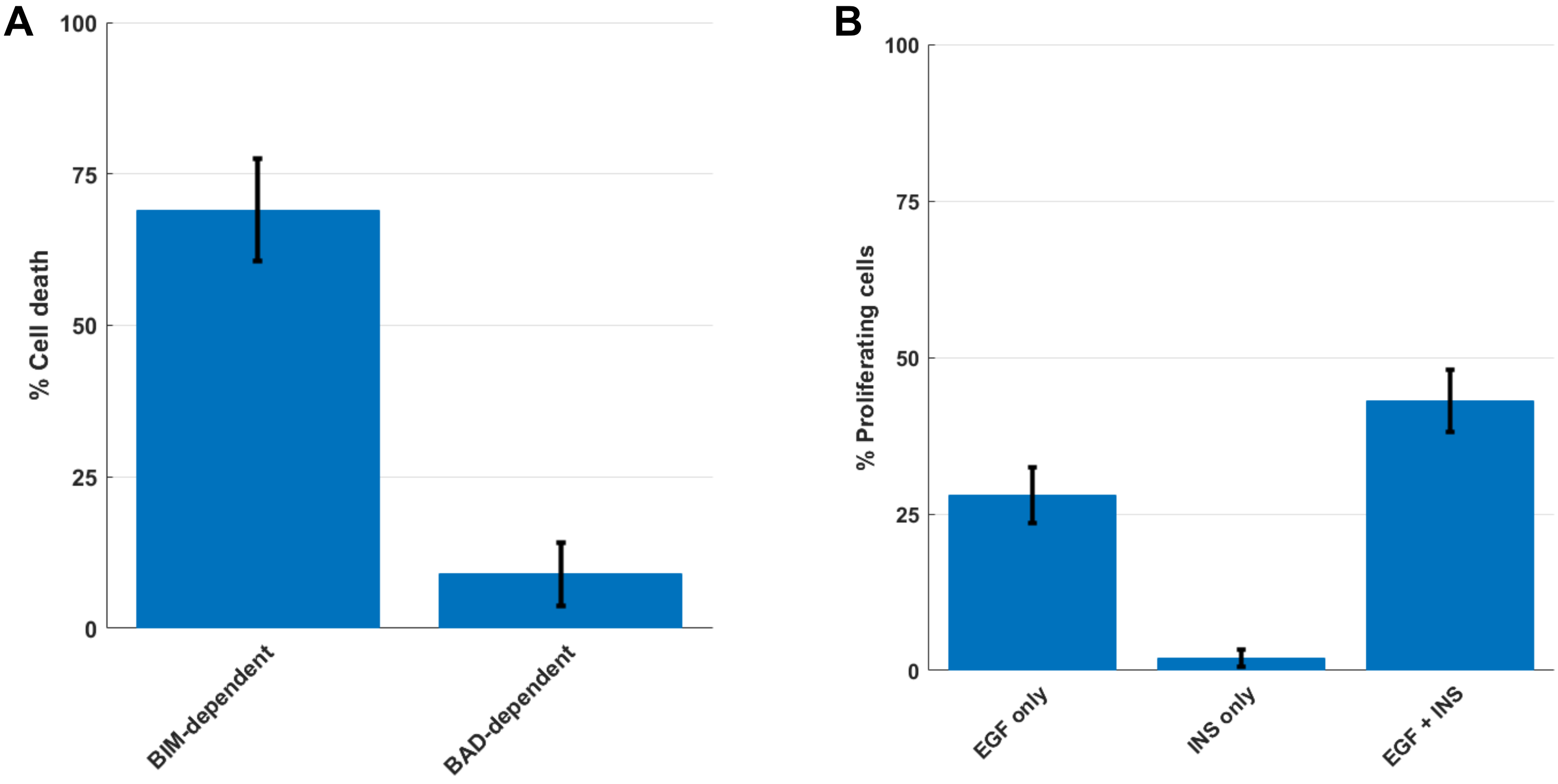


**Figure S10.** (A) Simulations where BIM-dependent or BAD-dependent mechanisms are switched off and percent death calculated in response to EGF + insulin at 48 hours. The results show that ERK and AKT inhibition induced cell death mechanisms are mostly BIM dependent, not BAD. Bars represent mean ± s.e.m. of 100 stochastic cell simulations. (B) EGF and insulin cooperatively induce cell cycle entry, with insulin inducing very little cell cycle entry alone. Cells are simulated with EGF (10nM), Insulin (1721nM), or EGF+Insulin (10nM+1721nM) for 30 hours and the percentage of cells entering S-phase are calculated. Cells are considered in S-phase when the sum of concentrations of Cyclin E, A, and B is greater than 20nM. Bars represent mean ± s.e.m. of 100 stochastic cell simulations.

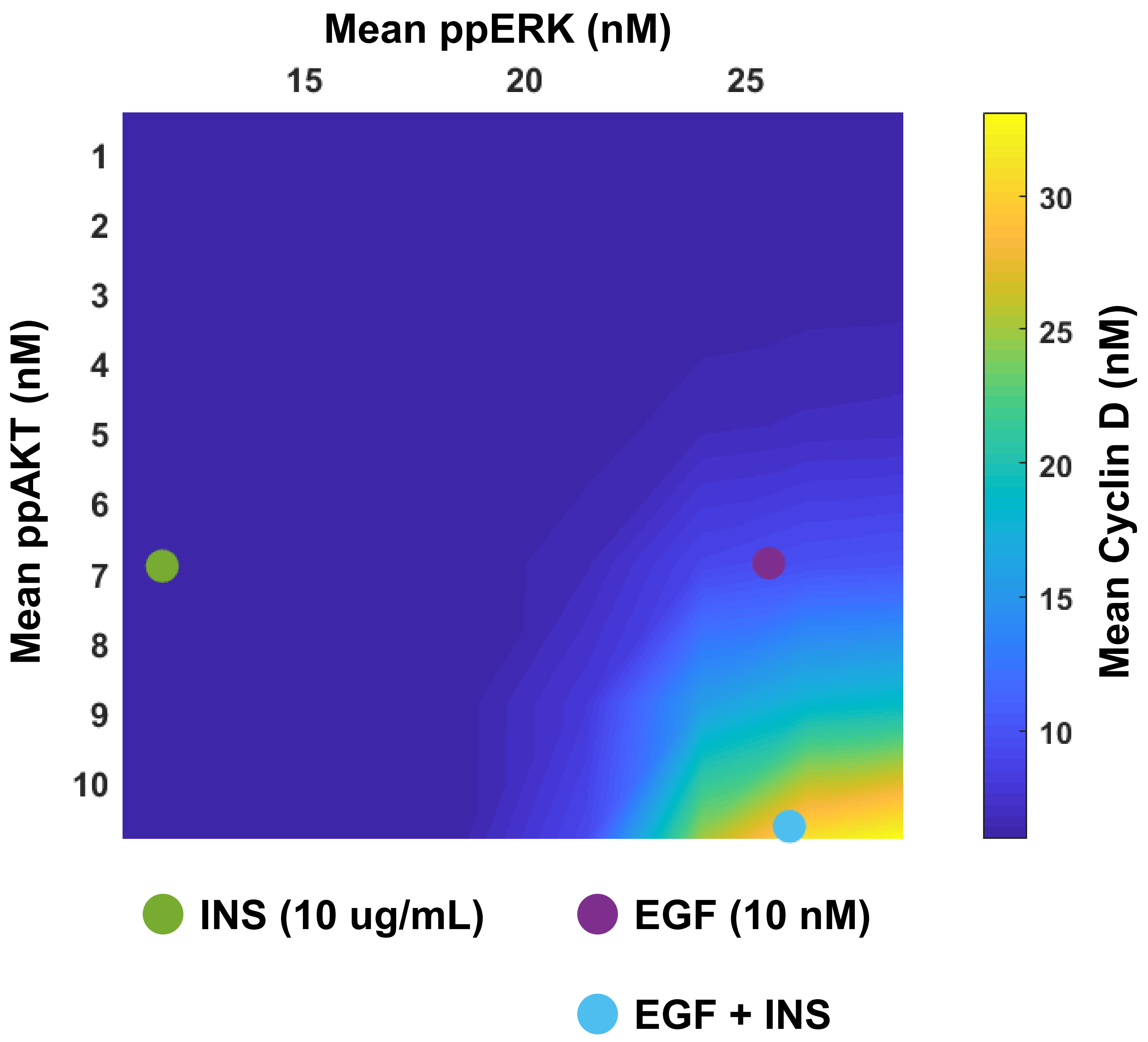


**Figure S11.** Activation of both ERK and AKT pathways are required for robust cell cycle entry. Time averaged ppERK and ppAKT levels correlate with Cyclin D levels. Basal levels of ppERK and ppAKT are increased (between 1X-20X) and each condition is simulated up to 6 hours. The time-averaged levels of ppERK and ppAKT are plotted against the time-averaged Cyclin D levels. Conditions representing EGF (10nM), Insulin (1721nM), and EGF+Insulin (10nM+1721nM) are shown with colored circles.

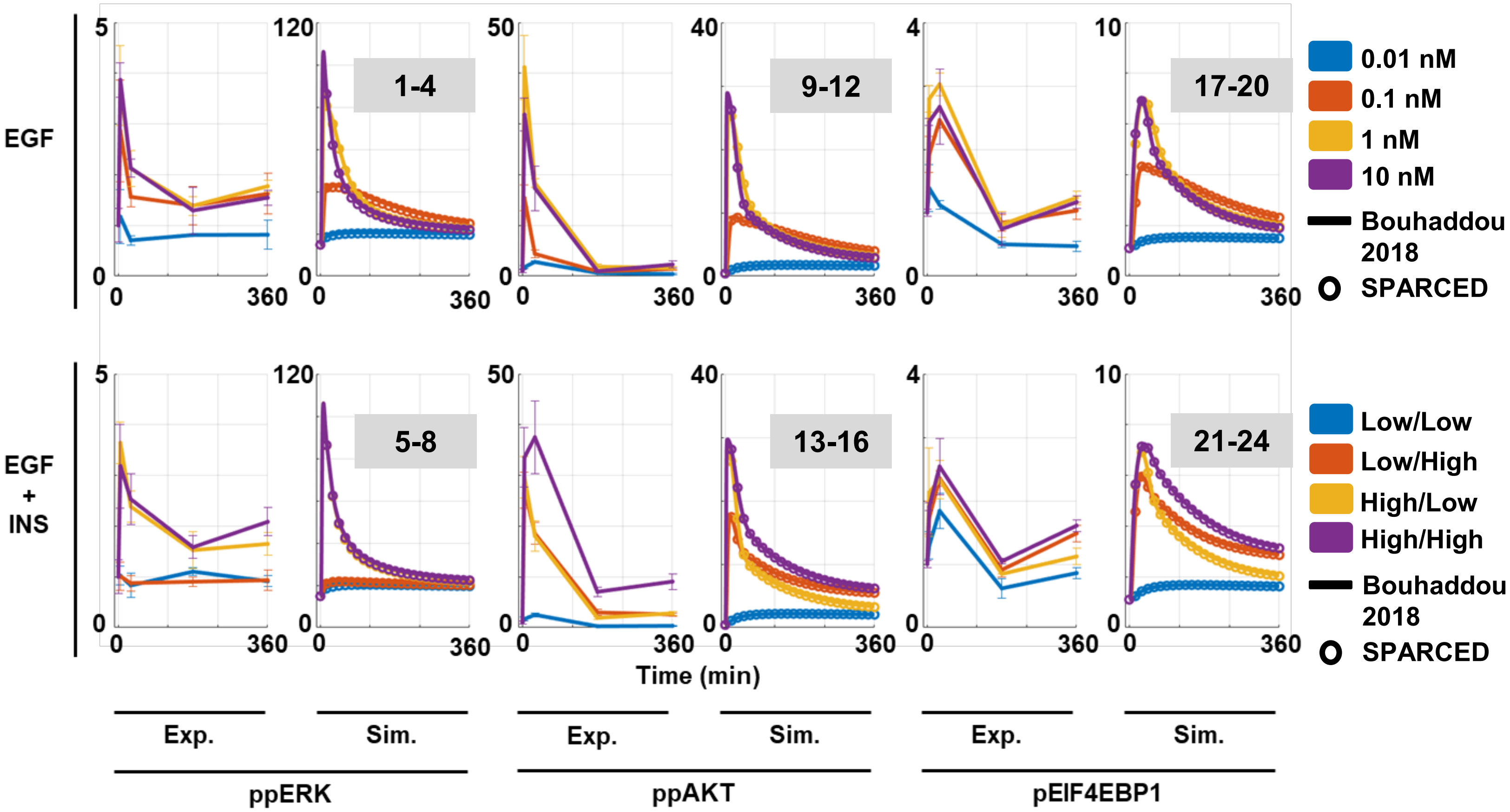


**Figure S12. SPARCED model recapitulates downstream pathway activation by ligands and ligand combination treatments.** Experimental data and simulation results from MATLAB (lines) and SPARCED (circles) models with EGF (top) and EGF+Insulin (bottom) stimulation for 6 hours. Plots show double-phosphorylated ERK (ppERK), serine-phosphorylated AKT (pAKT), and phospho-EIF4EBP1 (pEIF4EBP1) levels. The numbers in gray shaded boxes represents numbering of conditions in Fig. 2A. Exp: Experimental data, Sim: Simulation.

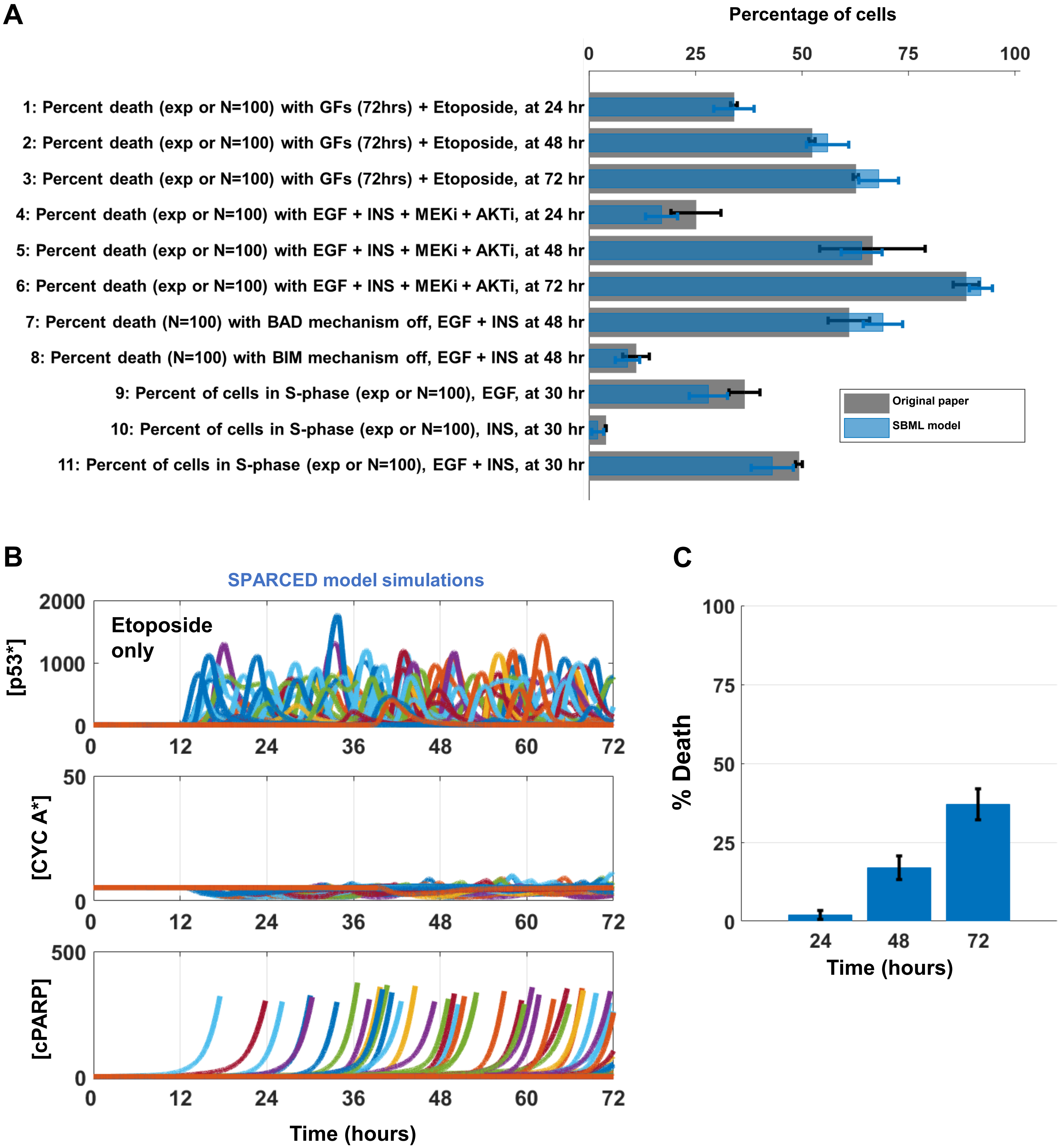


**Figure S13.** (A) Bar plots corresponding to the conditions shown in Fig. 2C. Gray bars are experimental or simulation data from Bouhaddou 2018 model and blue bars are simulation results of SPARCED model. Bars represent mean ± s.e.m. (B) Etoposide treatment alone induces lesser cell death compared to Etoposide + Growth Factor stimulation, shown in Fig. 3D-E. (C) Percentage of cell death of 100 cells shown in (B). Bars represent mean ± s.e.m.

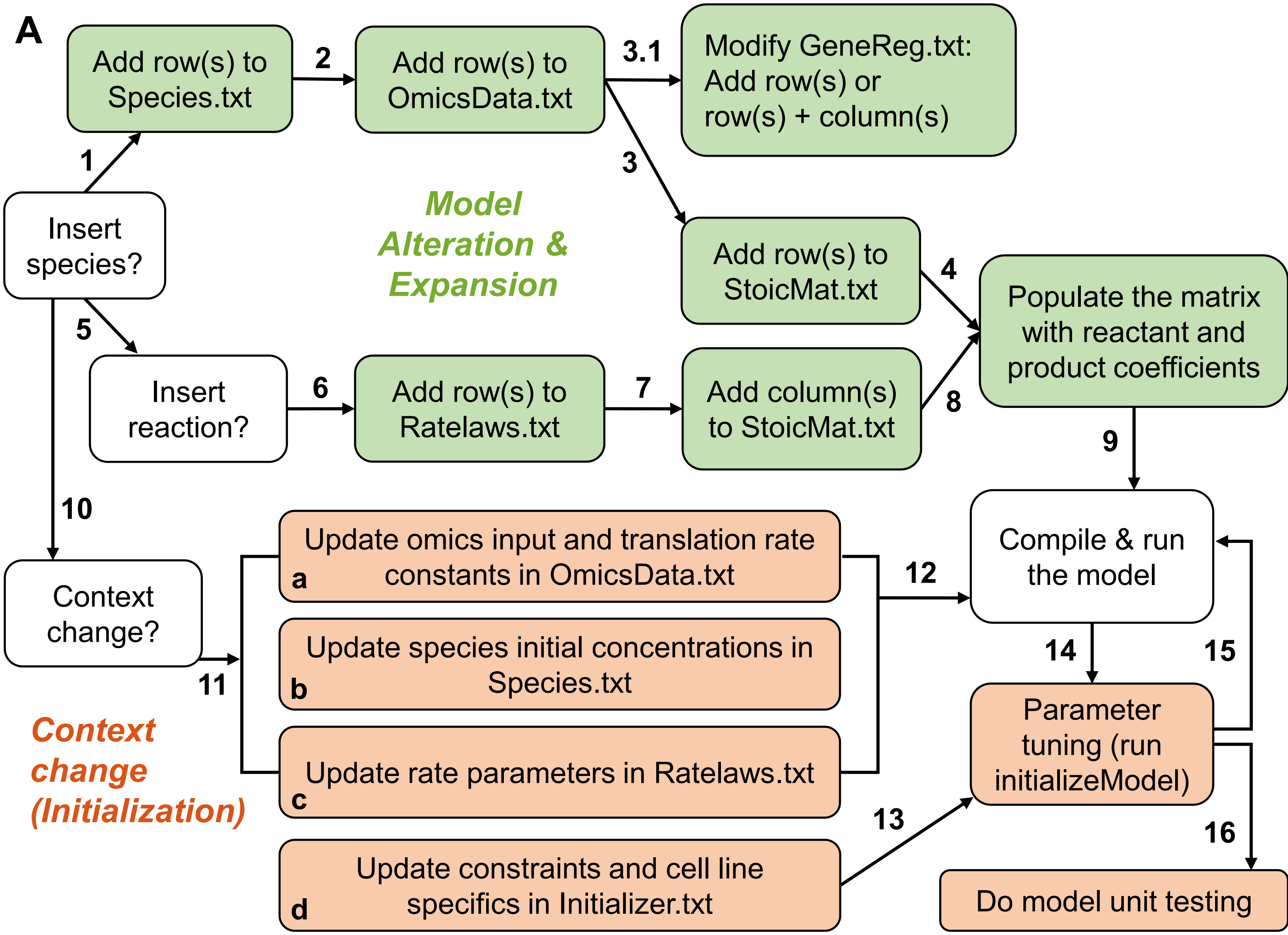


**B**

| Parameter category changed | Number of parameters changed (out of 3275 possible) | Input file |
|---|---|---|
| mRNA counts | 141 | OmicsData.txt |
| Gene copy numbers | 141 | OmicsData.txt |
| Constitutive mRNA translation rates | 141 | OmicsData.txt |
| Induced mRNA translation rates (from Bouhaddou2018) | 141 | Ratelaws.txt |
| Degradation rate constants (from Bouhaddou2018) | 46 | Ratelaws.txt |
| Species initial concentrations (from Bouhaddou2018) | 914 (out of 914) | Species.txt |

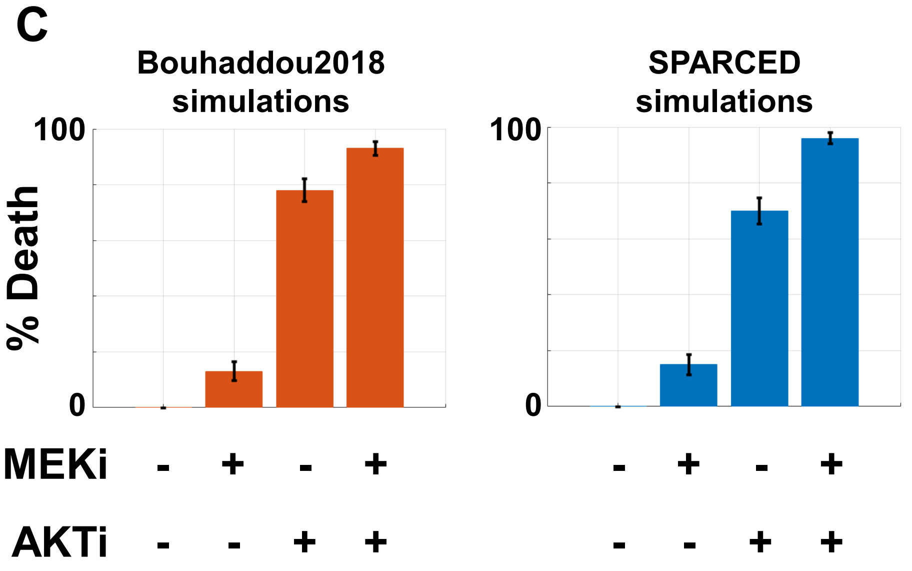


**Figure S14. SPARCED model alteration guidelines.** (A) Steps of model expansion and context change procedures are listed. Refer to Table 2 for more details. Steps can be skipped if no changes are necessary. (B) The list of parameters and species values modified for SPARCED model context change from MCF10A cells to U87 cells. (C) SPARCED_U87 model simulations reproduce previous observations, where U87 cells show increased response and sensitivity to AKT inhibition. MEKi: MEK inhibitor, AKTi: AKT inhibitor. Bars represent mean ± s.e.m. of 100 single cell simulations for each condition.

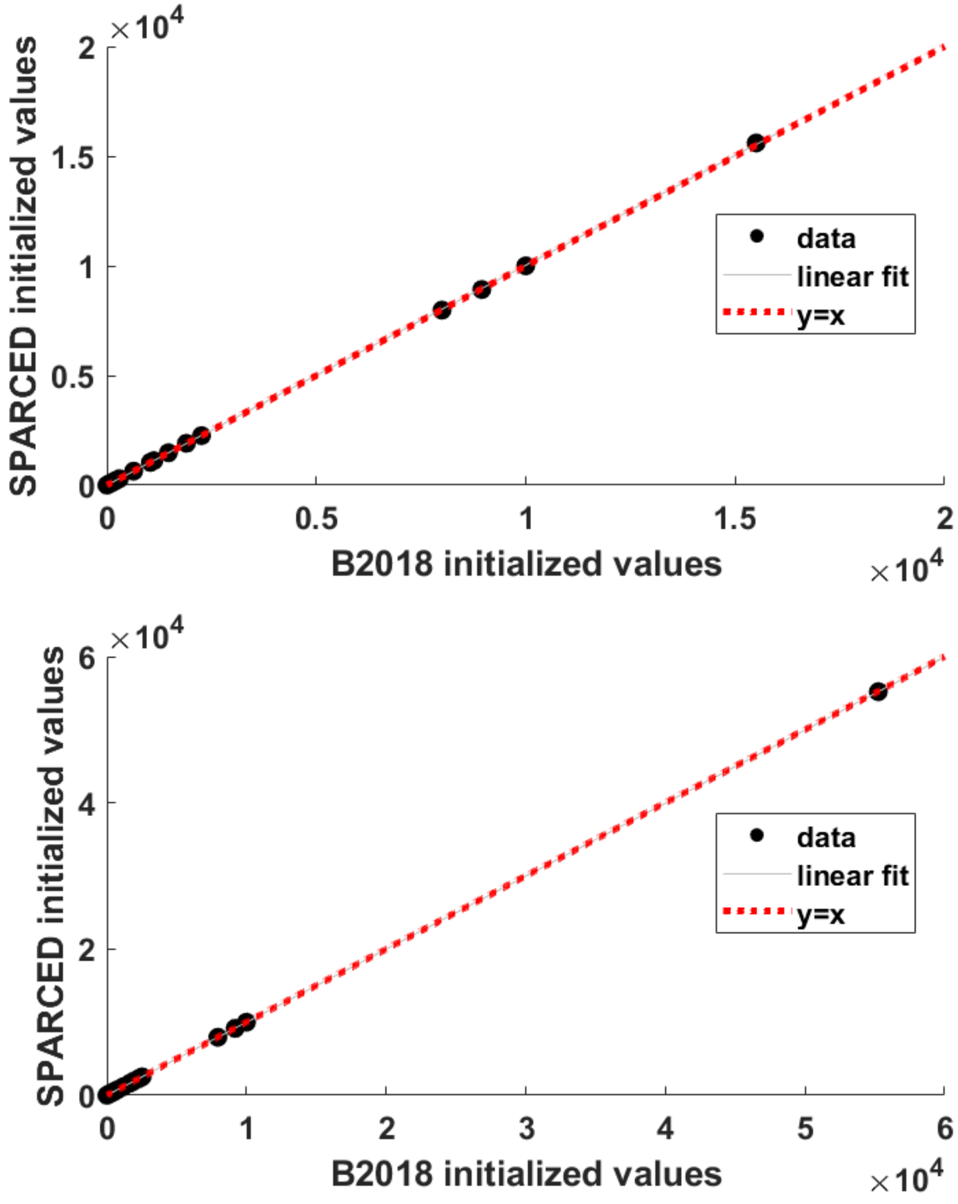


**Figure S15.** Comparison of initialized species concentrations for MCF10A (top) and U87 (bottom) cells. The Bouhaddou2018 model values are reproduced by the new initialization notebook. The concentrations values (black dots) are almost exact and identity line (dashed red) coincides with the linear fit line (solid black). SPARCED initialized values are on y-axis and B2018 initialized values are on x-axis.

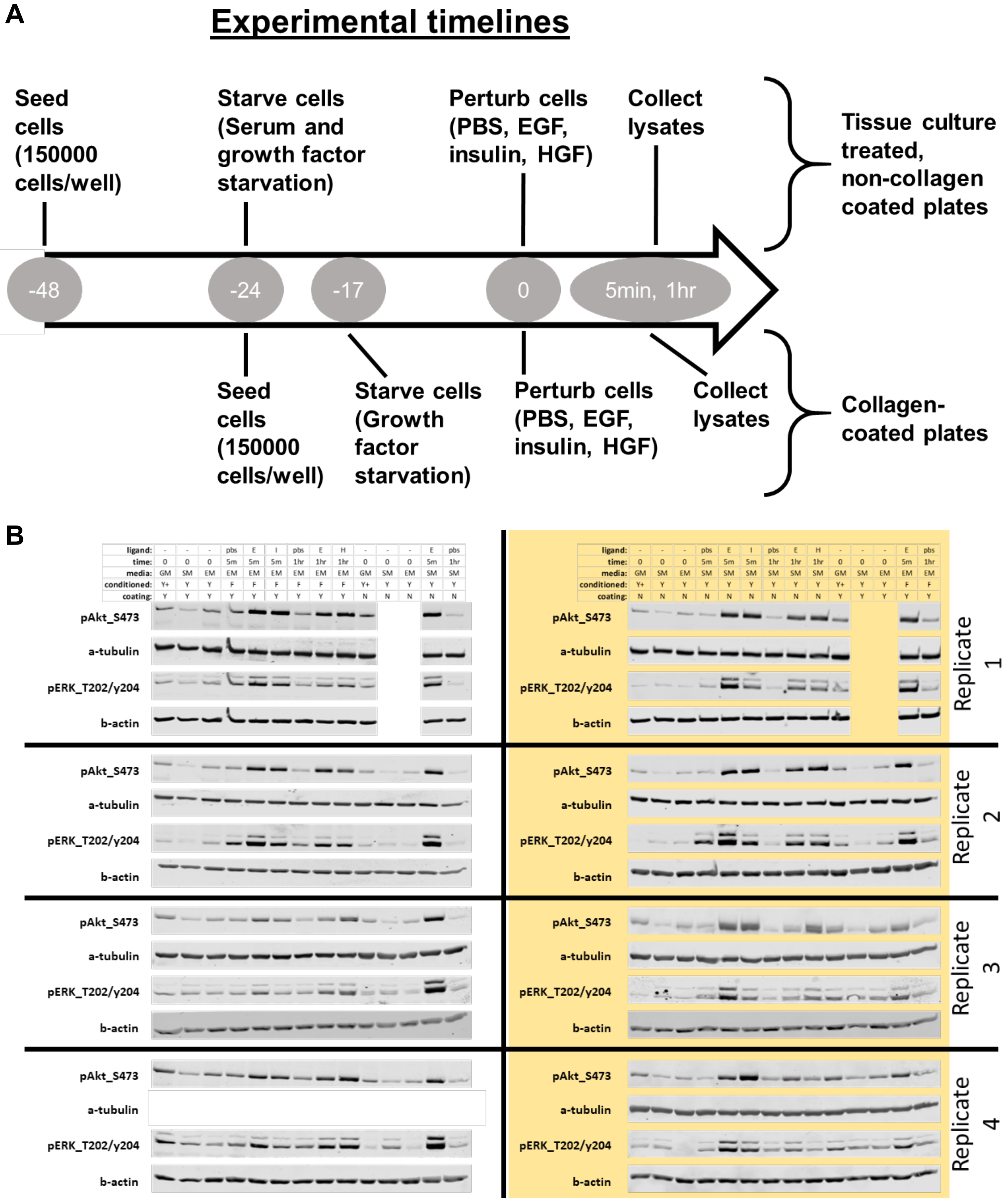


**Figure S16.** The experimental setup and results of exploring the effect of collagen-coating and serum starvation in MCF10A cells.

A

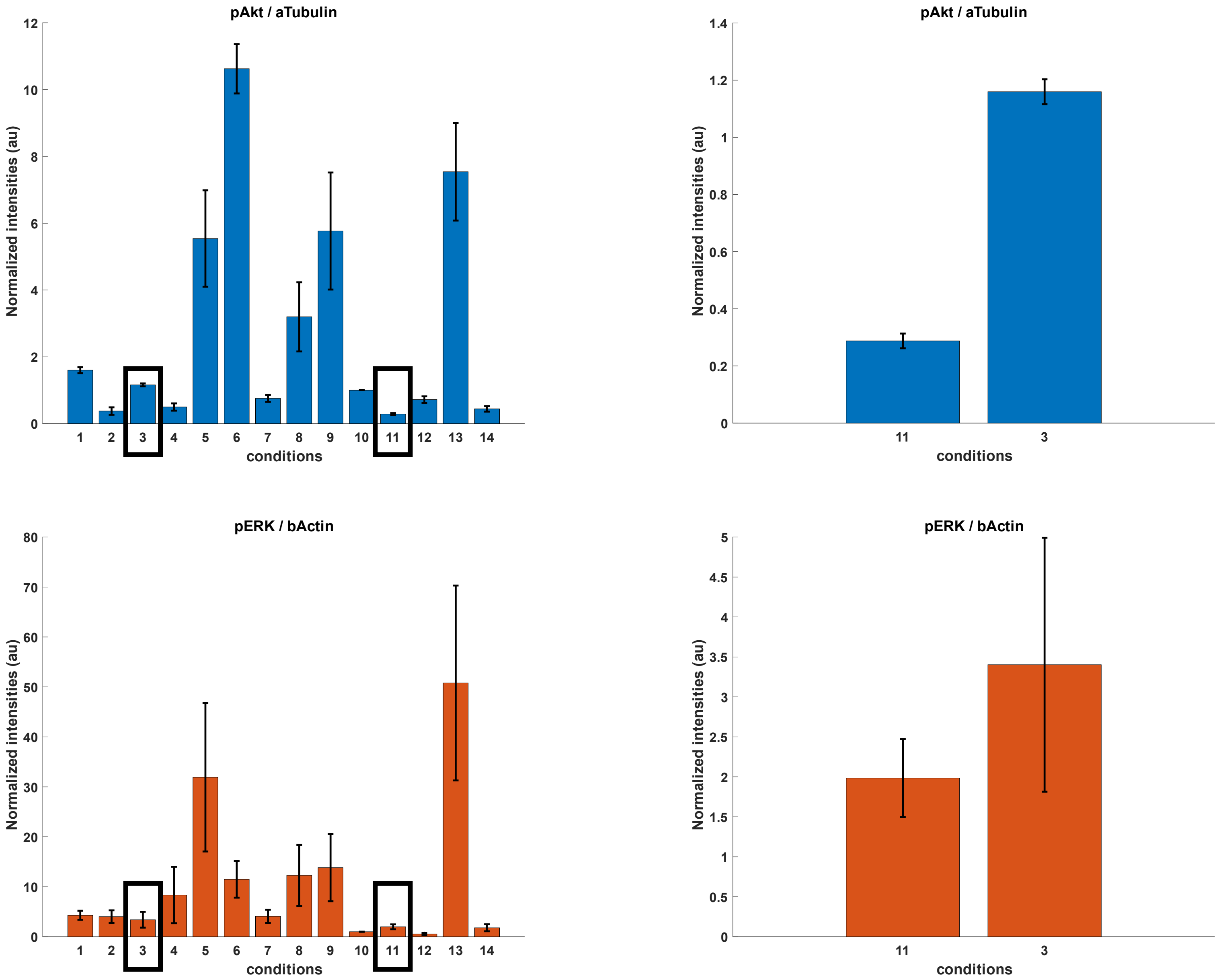


B

| Lane | coating | media | time | ligand | coating | media | time | ligand |
|---|---|---|---|---|---|---|---|---|
| **1** | Y | GM | 0 | - | N | GM | 0 | - |
| **2** | Y | SM | 0 | - | N | SM | 0 | - |
| **3** | Y | EM | 0 | - | N | EM | 0 | - |
| **4** | Y | EM | 5m | PBS | N | SM | 5m | PBS |
| **5** | Y | EM | 5m | EGF | N | SM | 5m | EGF |
| **6** | Y | EM | 5m | INS | N | SM | 5m | INS |
| **7** | Y | EM | 1hr | PBS | N | SM | 1hr | PBS |
| **8** | Y | EM | 1hr | EGF | N | SM | 1hr | EGF |
| **9** | Y | EM | 1hr | HGF | N | SM | 1hr | HGF |
| **10** | N | GM | 0 | - | Y | GM | 0 | - |
| **11** | N | SM | 0 | - | Y | SM | 0 | - |
| **12** | N | EM | 0 | - | Y | EM | 0 | - |
| **13** | N | SM | 5m | EGF | Y | EM | 5m | EGF |
| **14** | N | SM | 1hr | PBS | Y | EM | 1hr | PBS |

**Figure S17.** (A) The quantification of western blots shown in Fig. S18. All data are normalized to condition 10 (cells in non-coated plates with full growth media, measurements at time 0 hr). At least three biological replicates. pAKT levels in growth factor starved cells in collagen-coated plates (#3) are four times higher than pAKT levels in serum+growth factor starved cells in non-coated plates (#11). No significant change in pERK levels between two conditions (#3 and #11). The order of bars are given according to the list on the left in panel (B). (B) The experimental conditions of wester blots membranes in Fig. S18.

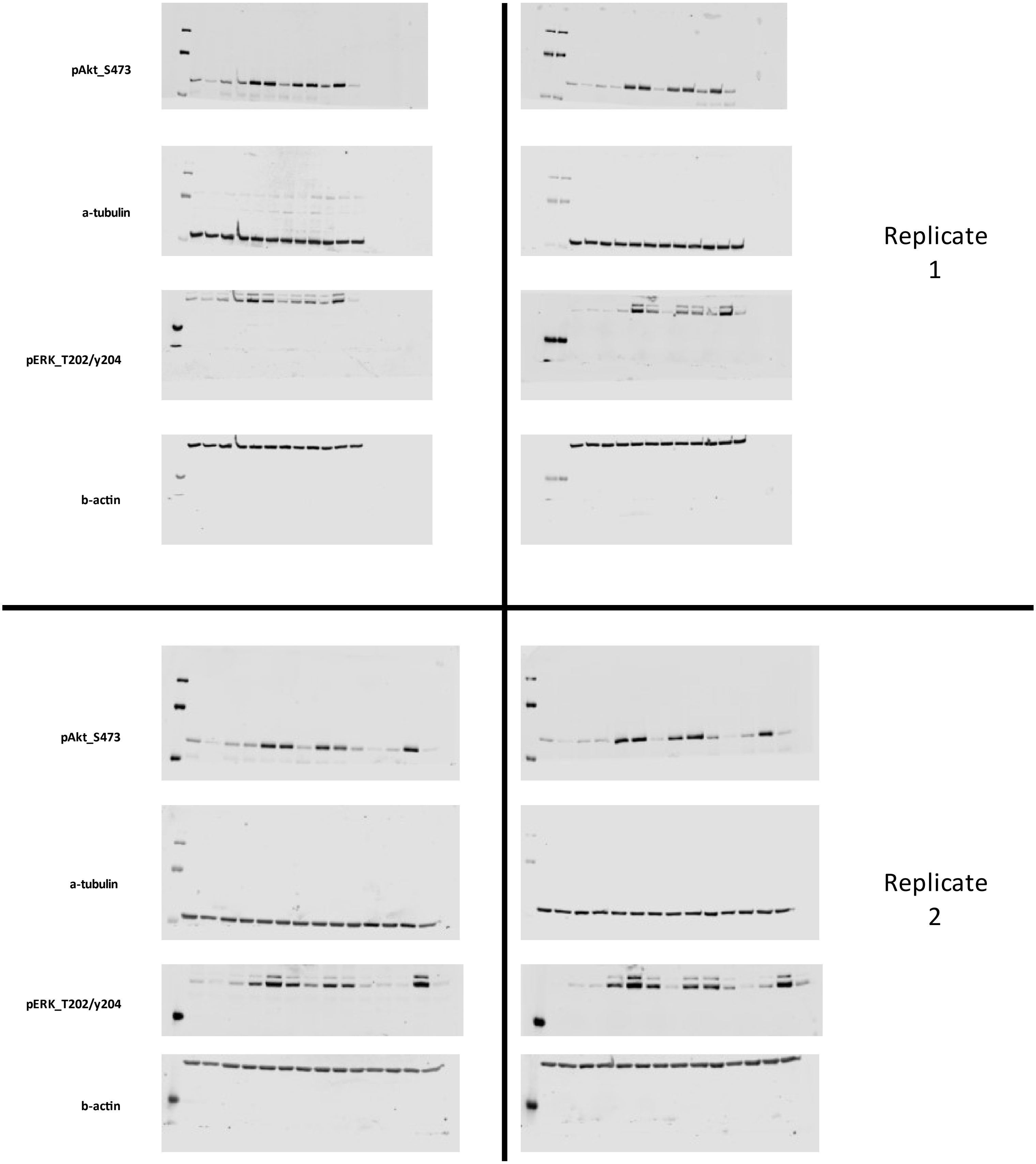


**Figure S18.** The full images of western blots (replicates 1 and 2) shown in Fig. S16.

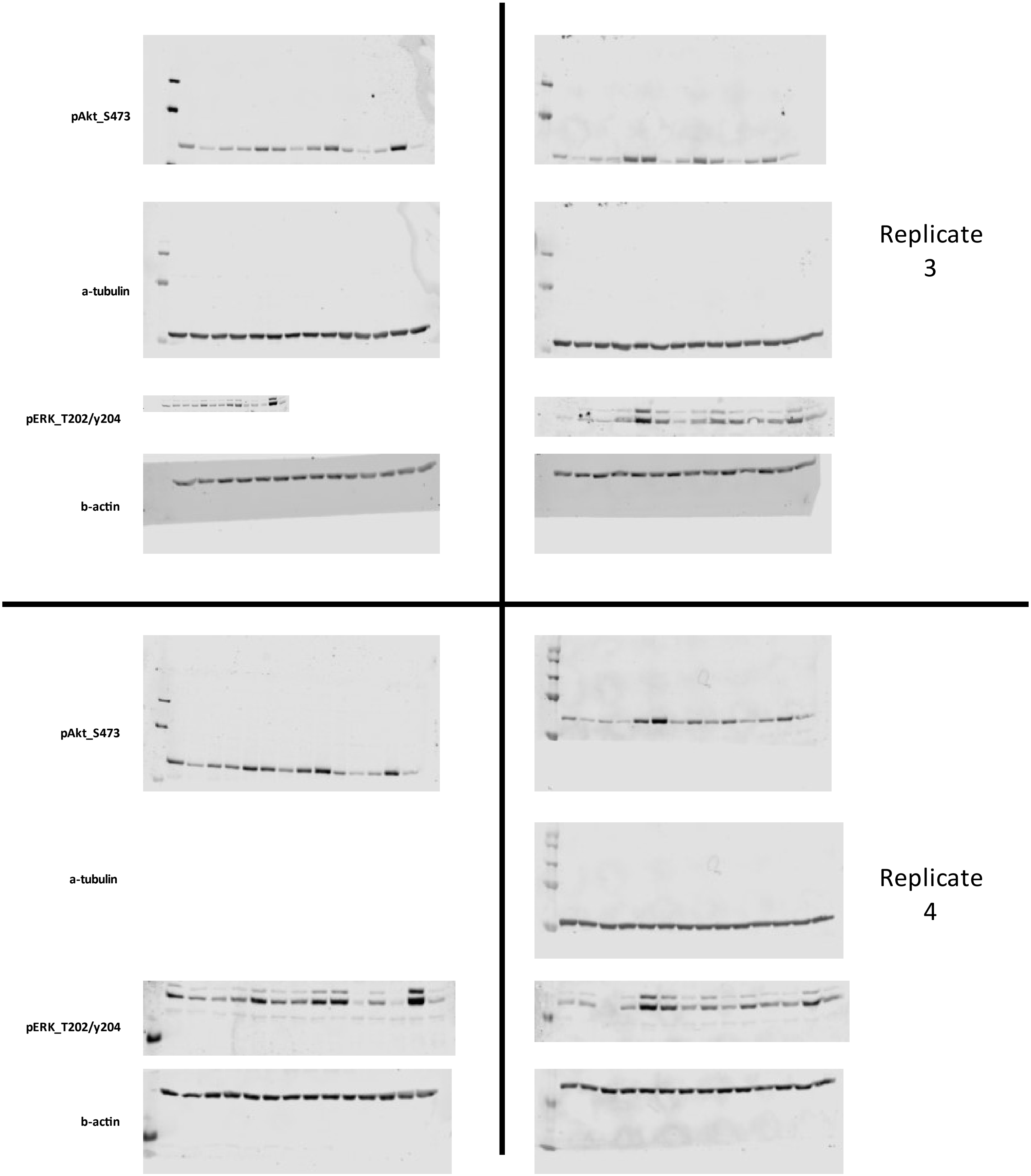


**Figure S19.** The full images of western blots (replicates 3 and 4) shown in Fig. S16.

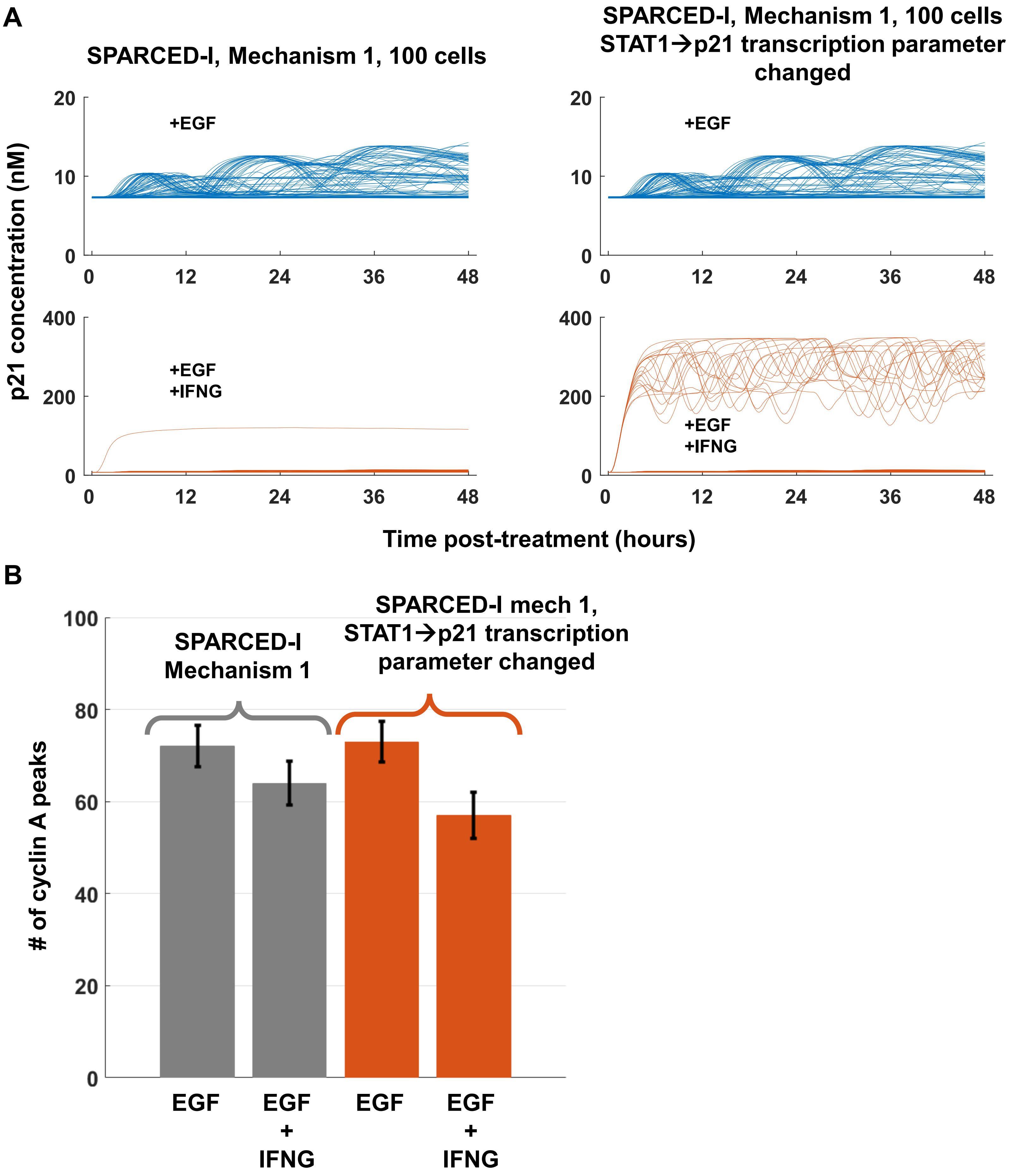


**Figure S20.** (A) p21 trajectories of SPARCED-I Mechanism 1 model simulations. Changing STAT1 regulation of p21 transcription induces higher p21 levels (orange, right) compared to original (orange, left) simulations. The p21 level increase does not occur in single EGF stimulation conditions (blue). (B) The total number of cyclin A peaks of 100 starting cells at 48 hours in SPARCED-I mech1 (gray) and modified SPARCED-I mech1 (orange) models.

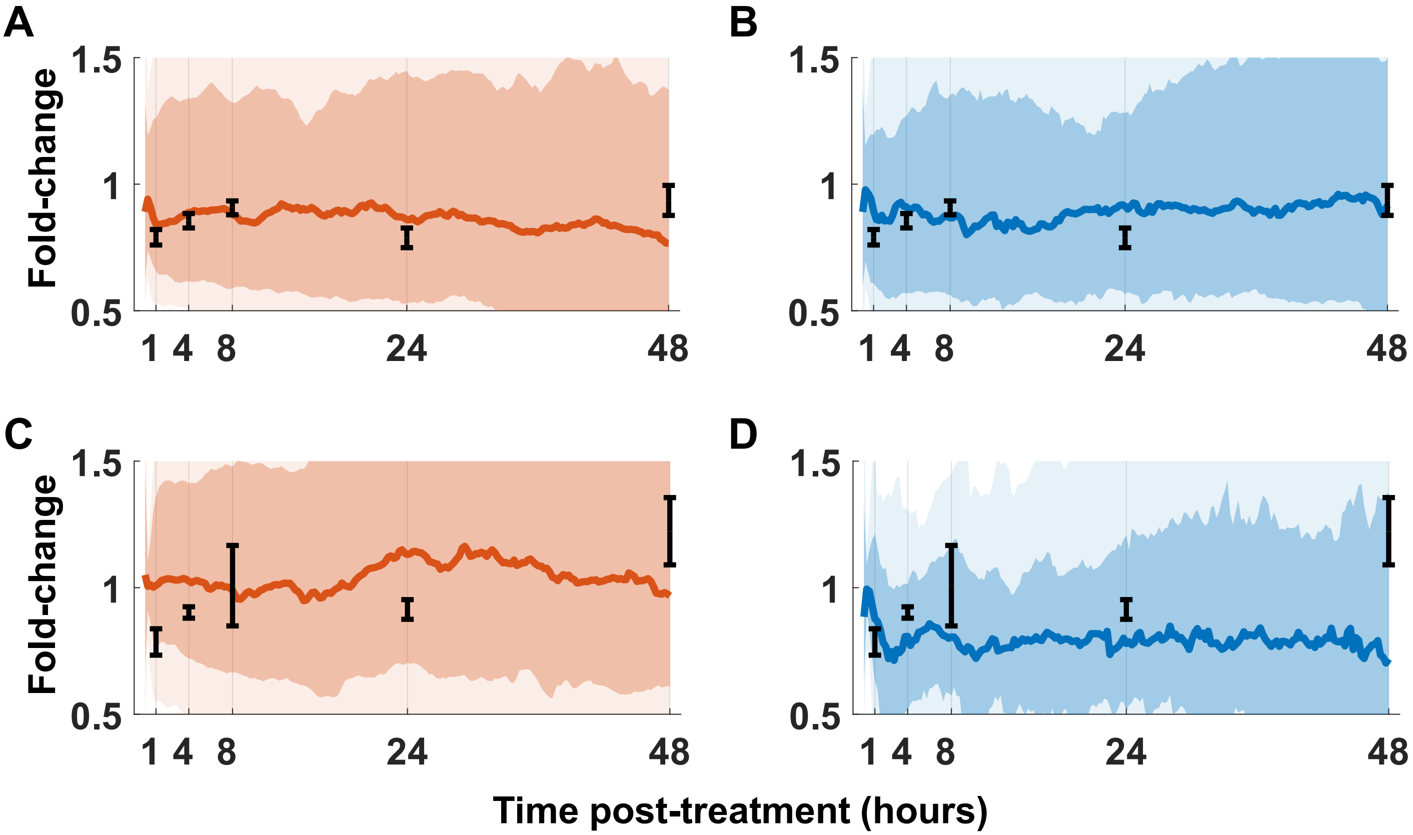


**Figure S21.** (A-B) Normalized pMAPK levels show a significant change when IFNG is included in addition to the EGF, in both Mechanism 1 (A) and Mechanism 2 (B). (C) Normalized ppAKT levels do not show a significant decrease after IFNG treatment, when EGF+IFNG Mechanism 1 simulations are compared to EGF alone case. (D) Normalized ppAKT levels show a significant decrease after IFNG treatment, when EGF+IFNG Mechanism 2 simulations are compared to EGF alone case. RPPA data are shown in black error lines, from three independent replicates. Colored dark lines represent median cell trajectories from simulations, dark and light-colored regions represent 70th and 95th quantiles, respectively. Experimental data from synapse.org/#!Synapse:syn12526172.
