## Supplemental Files and Figures for "A Scalable, Open-Source Implementation of a Large-Scale Mechanistic Model for Single Cell Proliferation and Death Signaling": SuppFile8_createModel.ipynb

```
# %% [markdown]
# SPARCED Model Creation

This file transforms the input files into Antimony and SBML model files. Here, we do not execute any simulation. 

# %% [markdown]
### These are the "default" input file names

They are structured, tab-separated text files, located in the "input files" folder by default.

# %% [code]
# Input file name definitions
fileComps = 'Compartments.txt' # input
fileSpecies = 'Species.txt' # input
fileStoic = 'StoicMat.txt' # input
fileRatelaws = 'Ratelaws.txt' # input
fileParamsOut = 'ParamsAll.txt' # output: Lists all parameter names, rxn names, values

# %% [markdown]
### Import required packages and scripts

# %% [code]
import sys
import os
sys.path.append(os.getcwd()[0:os.getcwd().rfind('/')]+'/bin')

import libsbml
import importlib
import amici
import numpy as np
import re
import pandas as pd
from antimony import *
from modules.copyDir import copyDirectory

# Optional packages to import
import amici.plotting
import matplotlib.pyplot as plt

# %% [markdown]
### Copy the input files from the "input_files" folder to the current folder

This step can be skipped if the input files are already in the working directory

# %% [code]
#copy input files over to current directory
current_dir = os.getcwd()
input_data_folder = current_dir[0:current_dir.rfind('/')+1]+'input_files'
copyDirectory(input_data_folder, os.getcwd()+"/")

# %% [markdown]
### Define the model (file) name here

First, the Antimony model file will be created using the input files. Users can add information per the antimony file structure. A default explanation for the SPARCED model is inserted here.

# %% [code]
# Antimony model name and information text
fileModel = open('SPARCED.txt','w') # file name
fileModel.write("# PanCancer Model by Birtwistle Lab \n") # some explanation
fileModel.write("model SPARCED()\n") # model name

# %% [markdown]
### Define the model SBML file name here

The SBML file will be created using the Antimony file. Define its name here. 
For consistency, we use the same name for the Antimony file, SBML file, and the model name.

# %% [code]
# SBML file name
sbml_file = 'SPARCED.xml'
# Name of the model that will also be the name of the python module
model_name = sbml_file[0:-4] 
# Directory to which the generated model code is written
model_output_dir = model_name 
# The AMICI package will create this folder while compiling the model, refer to AMICI github page for more details

# %% [markdown]
## Input file processing

Below are steps of input file processing and writing the Antimony file.

# %% [markdown]
### First, read-in the "Compartments" and "Volumes"

# %% [code]
# Initializing compartment and volume lists
compartments = []
volumes = []

# Create/write compartments
compartment_sheet = np.array([np.array(line.strip().split("\t")) for line in open(fileComps)])

#read in each line minus the header row of compartments file
for row in compartment_sheet[1:]:
    compartments.append(row[0])
    volumes.append(row[1])

# %% [markdown]
#### Set (write) the Compartments's names and assing the Volume values

# %% [code]
fileModel.write("\n  # Compartments and Species:\n") # Antimony Compartments/Species module title
for idx in range(len(compartments)):
    compName = compartments[idx]
    fileModel.write("  Compartment %s; " % (compName))
fileModel.write("\n")

# %% [markdown]
### Second, read-in the "Species" and related information

# %% [code]
# Write species and assign compartments
species_sheet = np.array([np.array(line.strip().split("\t")) for line in open('Species.txt', encoding='latin-1')])

# %% [code]
#read in each line minus the header row of species file
species_compartments = [] # Create a list of "home" compartments for each species in the model
for row in species_sheet[1:]:
    species_compartments.append(row[1]) 
species_compartments = np.array(species_compartments)

# %% [markdown]
#### Write the species names and their compartments in the Antimony format

# %% [code]
#Write each species to model txt file
fileModel.write("\n")
for idx,val in enumerate(species_sheet[1:]):
    fileModel.write("  Species ")
    fileModel.write("%s in %s" % (val[0], species_compartments[idx]))
    fileModel.write(";\n")

# %% [markdown]
### Third, read-in the "Reactions" and "StoicMat" files to create reactions of the model

# %% [code]
# Write reactions
fileModel.write("\n\n  # Reactions:\n") # Antimony Reactions module title

#reads in file from excel and gets rid of first row and column (they're data labels)
stoic_sheet = np.array([np.array(line.strip().split("\t")) for line in open('StoicMat.txt')])

#creates associated ratelaw data list
ratelaw_sheet = np.array([np.array(line.strip().split("\t")) for line in open('Ratelaws.txt')])
ratelaw_data = np.array([line[1:] for line in ratelaw_sheet[1:]])

#gets first column minus blank space at the beginning, adds to stoic data list
stoic_columnnames = stoic_sheet[0]
stoic_rownames = [line[0] for line in stoic_sheet[1:]]
stoic_data = np.array([line[1:] for line in stoic_sheet[1:]])

# %% [markdown]
#### Create the reations of the model based on the stoichiometric input and (if available) ratelaw input

# %% [code]
# builds the important ratelaw+stoic lines into the txt file 
paramnames = []
paramvals = []
paramrxns = []
paramidxs = []
for rowNum, ratelaw in enumerate(ratelaw_data):
    reactants = []
    products = []
    formula="k"+str(rowNum+1)+"*"

    for i, stoic_rowname in enumerate(stoic_rownames):
        stoic_value = int(stoic_data[i][rowNum])
        if stoic_value < 0:
            for j in range(0,stoic_value*-1):
                reactants.append(stoic_rowname)
                formula=formula+stoic_rowname+"*"
        elif stoic_value > 0:
            for j in range(0,stoic_value):
                products.append(stoic_rowname)

    if "k" not in ratelaw[1]:
        # the mass-action formula
        formula=formula[:-1]
        #the parameter
        paramnames.append("k"+str(rowNum+1))
        paramvals.append(np.double(ratelaw[1]))
        paramrxns.append(ratelaw_sheet[rowNum+1][0])
        paramidxs.append(int(0))
    else:
        # specific formula (non-mass-action)
        formula = ratelaw[1]
        j = 1
        params = np.genfromtxt(ratelaw[2:], float) # parameters
        params = params[~np.isnan(params)]
        if len(params) == 1:
            paramnames.append("k"+str(rowNum+1)+"_"+str(j))
            paramvals.append(float(ratelaw[j+1]))
            paramrxns.append(ratelaw_sheet[rowNum+1][0])
            paramidxs.append(int(0))
            pattern = 'k\D*\d*'
            compiled = re.compile(pattern)
            matches = compiled.finditer(formula)
            for ematch in matches:
                formula = formula.replace(ematch.group(),paramnames[-1])
        else:
            for q,p in enumerate(params):
                paramnames.append("k"+str(rowNum+1)+"_"+str(j))
                paramvals.append(float(ratelaw[j+1]))
                paramrxns.append(ratelaw_sheet[rowNum+1][0])
                paramidxs.append(q)
                pattern1 = 'k(\D*)\d*'+'_'+str(j)
                compiled1 = re.compile(pattern1)
                matches1 = compiled1.finditer(formula)
                for ematch in matches1:
                    formula = formula.replace(ematch.group(),paramnames[-1])
                j +=1
    if ratelaw[0] == 'Cytoplasm':
        valcomp = 5.25e-12
    elif ratelaw[0] == 'Extracellular':
        valcomp = 5.00e-5
    elif ratelaw[0] == 'Nucleus':
        valcomp = 1.75e-12
    elif ratelaw[0] == 'Mitochondrion':
        valcomp = 3.675e-13
    #don't include reactions without products or reactants
    if products == [] and reactants == []:
        pass
    else:
        fileModel.write("  %s: %s => %s; (%s)*%.6e;\n" % (stoic_columnnames[rowNum], " + ".join(reactants), " + ".join(products), formula, valcomp))

# Export parameters for each reaction, with corresponding order within the ratelaw and its value
params_all = pd.DataFrame({'value':paramvals,'rxn':paramrxns,'idx':paramidxs},index=paramnames)
params_all.to_csv(fileParamsOut,sep='\t',header=True, index=True)

# %% [markdown]
### Next, we set the "Initial Conditions" for Compartments, Species, and Parameters

# %% [markdown]
#### The Compartment initial conditions

# %% [code]
# Write compartment ICs
fileModel.write("\n  # Compartment initializations:\n")
for idx in range(len(compartments)):
    fileModel.write("  %s = %.6e;\n" % (compartments[idx], np.double(volumes[idx])))
    fileModel.write("  %s has volume;\n" % (compartments[idx]))

# %% [markdown]
#### The Species initial conditions

# %% [code]
# Write species ICs
fileModel.write("\n  # Species initializations:\n")
for idx, val in enumerate(species_sheet[1:]):
    fileModel.write("  %s = %.6e;\n" % (val[0],np.double(val[2])))

# %% [markdown]
#### The Parameter (of reactions) initial conditions

# %% [code]
# Write parameter ICs
fileModel.write("\n  # Parameter initializations:\n")
count = 0
for param in paramnames:
    fileModel.write("  %s = %.6e;\n" % (param, np.double(paramvals[count])))
    count += 1

# %% [markdown]
### Other declarations supported by Antimony

# %% [code]
# Write other declarations
constantVars = ['Cytoplasm','Extracellular','Nucleus','Mitochondrion']

fileModel.write("\n  # Other declarations:\n")
fileModel.write("  const")
for constVar in constantVars[:-1]:
    fileModel.write("  %s," % (constVar))
#last item in row needs semicolon
fileModel.write("  %s;\n" % (constantVars[-1]))

# %% [markdown]
### The Unit Definitions

We define the time, volume, and substance units

# %% [code]
# Write unit definitions
fileModel.write("\n  # Unit definitions:")
fileModel.write("\n  unit time_unit = second;")
fileModel.write("\n  unit volume = litre;")
fileModel.write("\n  unit substance = 1e-9 mole;")
fileModel.write("\n  unit nM = 1e-9 mole / litre;")
fileModel.write("\n")

# %% [markdown]
### End of the Antimony file, close the file

# %% [code]
# End the model file
fileModel.write("\nend")
# Close the file
fileModel.close()

# %% [markdown]
## The Antimony file import and conversion to SBML format


# %% [code]
# load model and convert to SBML
if loadFile("SPARCED.txt") == 1:
    print("Success loading antimony file")
else:
    print("Failed to load antimony file")
    exit(1)

if writeSBMLFile("SPARCED.xml","SPARCED") == 1:
    print("Success converting antimony to SBML")
else:
    print("Failure converting antimony to SBML")
    exit(1)

# %% [markdown]
#### The initial SBML file is imported again, to add annotations for the Compartments and Species

Currently, the Antimony file format does not support Annotation definitions. So, we re-process the SBML file and add Annotations for the Compartments (GO terms) and Species (HGNC or ENSEMBL identifiers).

# %% [markdown]
#### Import the SBML file and get the model handle

# %% [code]
# create interaction components
sbml_reader = libsbml.SBMLReader()
sbml_doc = sbml_reader.readSBML(sbml_file)
sbml_model = sbml_doc.getModel()

# %% [markdown]
#### The Species annotations are set using the last column of the Species input file

# %% [code]
# Set species annotations
for idx,row in enumerate(species_sheet[1:]):
    Annot=""
    for col in range(4,(len(row))):
        aa=str(row[col].strip())
        if aa=="nan" or aa == "":
            break
        else:
            Annot=Annot+" "+row[col]
    sbml_model.getSpecies(row[0]).setAnnotation(Annot)

# %% [markdown]
#### The Compartment annotations are set using the last column of the Compartments input file

# %% [code]
# Set compartment annotations
for row in compartment_sheet[1:]:
    sbml_model.getCompartment(row[0]).setAnnotation(row[2])

# %% [markdown]
#### Export the finalized (annotated) SBML file

# %% [code]
# Write with the same name or use the next section instead of below lines
writer = libsbml.SBMLWriter()
writer.writeSBML(sbml_doc, sbml_file)

# %% [markdown]
## The Model Compilation

# %% [markdown]
### Import the SBML file and compile using the AMICI package

# %% [code]
# prepares to use interaction components to synthesize model
sys.path.insert(0, os.path.abspath(model_output_dir))
model_name = sbml_file[0:-4]
model_output_dir = model_name

sbml_reader = libsbml.SBMLReader()
sbml_doc = sbml_reader.readSBML(sbml_file)
sbml_model = sbml_doc.getModel()

# Create an SbmlImporter instance for our SBML model
sbml_importer = amici.SbmlImporter(sbml_file)

# %% [markdown]
#### Setting AMICI converter arguments

We set all the rate parameters as constant for faster compilation. It is also a good idea to define "Observables" for easier comparison to experimental data. An observable in our model corresponds to the totality of a protein sepcies, including all of its post-translationally modified and unmodified states, corrected for compartmental volume differences and reported based on cytoplasmic volume. We defined 102 observables using the "Observables.txt" input file in `AMICI` required format.

# %% [code]
#sets important constants for model build
constantParameters = [params.getId() for params in sbml_model.getListOfParameters()]

# %% [code]
# creates observables.
ObsMat = pd.read_csv('Observables.txt', sep='\t',header=0, index_col=0)
Vc = float(compartment_sheet[compartment_sheet[:,0]=='Cytoplasm',1])

species_names = np.array([species_sheet[i][0] for i in range(1,len(species_sheet))])
Vol_species = np.array([species_sheet[i][1] for i in range(1,len(species_sheet))])
Vol_species = [float(compartment_sheet[compartment_sheet[:,0]==Vol_species[i],1][0]) for i in range(len(Vol_species))]
Vol_species = pd.Series(Vol_species, index=species_names)

formula_obs = []
for obs in ObsMat.columns:
    sp_obs = ObsMat.index[np.nonzero(np.array(ObsMat.loc[:,obs]>0))[0]]
    sp_obs_id = np.nonzero(np.array(ObsMat.loc[:,obs]>0))[0]
    Vr = Vol_species/Vc
    Vf = Vr*ObsMat.loc[:,obs].values
    if len(sp_obs) == 1:
        formula_i = sp_obs[0]+'*'+str(Vf[sp_obs_id][0])
    elif len(sp_obs) == 2:
        formula_i = str(sp_obs[0]+'*'+str(Vf[sp_obs_id][0])+'+'+sp_obs[1]+'*'+str(Vf[sp_obs_id][1]))
    elif len(sp_obs) > 2:
        formula_i = ''
        for j in range(len(sp_obs)-1):
            formula_i = formula_i+sp_obs[j]+'*'+str(Vf[sp_obs_id][j])+'+'
        formula_i = formula_i+str(sp_obs[-1])+'*'+str(Vf[sp_obs_id][-1])
    formula_obs.append(formula_i)

observables = {}
obs_names = list(ObsMat.columns)
for i in range(len(obs_names)):
    observables[obs_names[i]] = {}
    observables[obs_names[i]]['formula'] = formula_obs[i]

# %% [markdown]
#### Compile the model and create required C++ dependencies for simulation. 

Creates a sub-folder with the same name as the model (i.e. SPARCED)

# %% [code]
# The actual compilation step by AMICI, takes a while to complete for large models
sbml_importer.sbml2amici(model_name,
                         model_output_dir,
                         verbose=False,
                         observables=observables,
                         constantParameters=constantParameters)

# %% [markdown]
## The model creation is now complete! Enjoy...
```
