## Supplemental Files and Figures for "A Scalable, Open-Source Implementation of a Large-Scale Mechanistic Model for Single Cell Proliferation and Death Signaling": SuppFile11_runModel.ipynb

```
# %% [markdown]
# SPARCED Model Simulation

# %% [markdown]
This is how to simulate an existing SPARCED model, consisting of an SBML file and scripts for random sampling of mRNA counts for stochastic single-cell simulations. Below is an illustration of how a single-cell is simulated, with required inputs and arguments.

# %% [markdown]
## First, import the required packages

# %% [code]
import libsbml
import importlib
import amici
import amici.plotting
import numpy as np
import matplotlib.pyplot as plt
import pandas as pd
from datetime import datetime
import scipy.stats
import argparse
import sys
import os
sys.path.append(os.getcwd()[0:os.getcwd().rfind('/')]+'/bin')
from modules.RunSPARCED import RunSPARCED

# %% [markdown]
## Define the model SBML file name

# %% [code]
# SBML model we want to import
sbml_file = 'SPARCED.xml'
model_name = sbml_file[0:-4]
# Directory to which the generated model code was written by AMICI during model creation/compilation
model_output_dir = model_name

# %% [markdown]
## The required user input

Users should define if the simulation should be "deterministic for an average-cell" or "stochastic single-cell". Please refer to our manuscript for details and explanitions. In short, a "deterministic" run assumes an average cell response and does not randomly sample gene activation/inactivation and mRNA birth/death events. All simulations are deterministic. For a "stochastic" run, these events are assumed as Poission processes and each simulation represents a single-cell response. 

# %% [code]
# The flag for simulation type: deterministic=1, stochastic(hybrid)=0
flagD = 1
# The simulation time in hours
th = 1
# Input ligand concentrations (in order): EGF, Her, HGF, PDGF, FGF, IGF, INS
STIMligs = [100.0,100.0,100.0,100.0,100.0,100.0,1721.0] # in nM, in extracellular volume
# File name prior for saving the simulation trajectory and gene states
nmoutfile = 'GrowthStim_'

# %% [markdown]
#### Other required inputs

Below parameters are not usually changed. 

# %% [code]
ts = 30 # The time frame at which stochastic gene module and deterministic SBML module exhange/update information
cellNumber = 0 # used for filename to save

# %% [markdown]
### Species initial conditions 

We read-in the "Species" input file values here to set the initial conditions for simulations. The ligand concentrations are updated based on the user input above.

# %% [code]
sys.path.insert(0, os.path.abspath(model_output_dir))
species_sheet = np.array([np.array(line.strip().split("\t")) for line in open('Species.txt', encoding='latin-1')])

species_initializations = []
for row in species_sheet[1:]:
    species_initializations.append(float(row[2]))
species_initializations = np.array(species_initializations)
species_initializations[np.argwhere(species_initializations <= 1e-6)] = 0.0
species_initializations[155:162] = STIMligs

# %% [markdown]
### Model import

The existing model SBML is imported by AMICI and "solver"arguments are set.

# %% [code]
model_module = importlib.import_module(model_name)
model = model_module.getModel()
solver = model.getSolver() # Create solver instance
solver.setMaxSteps = 1e10 
model.setTimepoints(np.linspace(0,ts,2)) # set timepoints

# %% [markdown]
## Model simulation

The simulation step. The `RunSPARCED.py` function is called to simulate the model with the given user input. In short, the `RunSPARCED` function simulates the SBML module deterministically for 30 seconds, updates the transcriptional regulators for mRNA regulation, the stochastic sampling of gene activation/inactivations and mRNA birth/death events occur, and the new mRNA levels are updated for the next 30sec deterministic simulation. For more details, please refer to our manuscript.

# %% [code]
xoutS_all, xoutG_all, tout_all = RunSPARCED(flagD,th,species_initializations,[],sbml_file,model)

# %% [markdown]
### Save the output

The simulation trajectory and active/inactive genes at every 30sec are written respectively in "nmoutfile_S" and "nmoutfile_G" files.

# %% [code]
# saves output
columnsS = [ele for ele in model.getStateIds()]
columnsG = [x for n, x in enumerate(columnsS) if 'm_' in x]
columnsG = columnsG[1:]
resa = [sub.replace('m_', 'ag_') for sub in columnsG]
resi = [sub.replace('m_', 'ig_') for sub in columnsG]
columnsG2 = np.concatenate((resa, resi), axis=None)
condsSDF = pd.DataFrame(data=xoutS_all,columns=columnsS)
condsSDF.to_csv(nmoutfile+'S_'+str(cellNumber)+'.txt',sep="\t")  
condsSDF = None
condsGDF = pd.DataFrame(data=xoutG_all,columns=columnsG2)
condsGDF.to_csv(nmoutfile+'G_'+str(cellNumber)+'.txt',sep="\t") 
condsGDF = None
cellNumber+=1

# %% [markdown]
### Plot examples

# %% [code]
fig = plt.figure(figsize=(8, 2), dpi=300, facecolor='w', edgecolor='k')
tt = tout_all/3600.0

ax1 = fig.add_subplot(121)
plt.plot(tt, xoutS_all[:,717], 'r', linewidth=3, markersize=12, label='free ppERK')
plt.plot(tt, xoutS_all[:,696], 'b', linewidth=3, markersize=12, label='free ppAKT')
plt.xlim([-1,th+1])
plt.legend()
ax1.set_ylabel('Concentration (nM)')
ax1.set_xlabel('Time (hr)')
plt.xticks(np.arange(0, th+1, step=4))  # Set label locations.

ax1 = fig.add_subplot(122)
plt.plot(tt, xoutS_all[:,715], 'r--', linewidth=3, markersize=12, label='free total ERK')
plt.plot(tt, xoutS_all[:,694], 'b--', linewidth=3, markersize=12, label='free total AKT')
plt.legend()
ax1.set_xlabel('Time (hr)')
plt.xlim([-1,th+1])
plt.xticks(np.arange(0, th+1, step=4))  # Set label locations.
```
