## Supplemental Files and Figures for "A Scalable, Open-Source Implementation of a Large-Scale Mechanistic Model for Single Cell Proliferation and Death Signaling": SuppFile17_initializeModel_U87.ipynb

```
# %% [markdown]
# SPARCED Model Initialization

The model initialization procedure calibrates the parameter values (i) to make sure the tranlation rate constants induce a steady-state and (ii) to sustain certain phenotypic observations, such as serum-starved cells should not enter cell cycle or apoptosis.  

Each new cell line or cellular context (tumor microenvironement) should follow a custom `Initilization` procedure, based on user choices. So, below code should be handled as a guide on how to do this. 

Here, we made sure this script reproduces Bouhaddou2018 results. 

# %% [markdown]
## 1) Pre-processing and model preparation

Below are some adjustments and manual settings to match earlier model results.

# %% [code]
import pandas as pd
import numpy as np
import re
import libsbml
import os
import sys
import importlib
import amici
import amici.plotting

# %% [markdown]
### Load the model file, input files, and initialization specs

To start running `Initilization`, the user should first run `createModel` notebook and define the model SBML file name below. 

# %% [code]
#%% load model and input files
sbml_file = "SPARCED_U87.xml"
model_name= sbml_file[0:-4]
model_output_dir = model_name
sys.path.insert(0, os.path.join(model_output_dir))
model_module = importlib.import_module(model_name)
model = model_module.getModel()

sbml_reader = libsbml.SBMLReader()
sbml_doc = sbml_reader.readSBML(sbml_file)
sbml_model = sbml_doc.getModel()
params_model = []
[params_model.append(p.getId()) for p in sbml_model.getListOfParameters()]

# Ratelaws input file
ratelaw_sheet = np.array([np.array(line.strip().split("\t")) for line in open('Ratelaws.txt')])
ratelaw_data = np.array([line[1:] for line in ratelaw_sheet[1:]])

# OmicsData input file
gene_params = pd.read_csv('OmicsData_U87.txt', sep='\t', index_col=0, header=0)
model_genes = gene_params.index
mExp_mpc = gene_params['Exp RNA'].copy()

# Species input file
ICf = pd.read_csv('Species_U87.txt',header=0,index_col=0,sep='\t')

# Compartments input file
cell_params = pd.read_csv('Compartments.txt', header=0, index_col=0, sep='\t')
Vc = cell_params.loc['Cytoplasm','volume']
Vn = cell_params.loc['Nucleus','volume']
Vm = cell_params.loc['Mitochondrion','volume']
Ve = cell_params.loc['Extracellular','volume']
volumeofcell = Vc + Vn

# %% [markdown]
#### Define and calculate translation related parameters

Here, we define and calculate protein degradation rate constants. For definition of these parameter choices, please refer to Bouhaddou2018 paper.

# %% [markdown]
##### Here, notice the difference from 10A initialization procedure, where now we only have RNAseq data for U87 cells and use protein2mRNA ratios from 10A cells when needed.

# %% [code]
gene_params['kTLd'] = np.log(2)/gene_params['Protein_half_life_lit_h']/3600
gene_params['kTLd'][np.isnan(gene_params['kTLd'].values)] = np.log(2)/gene_params['Protein_half_life_Schwan_h'][np.isnan(gene_params['kTLd'].values)]/3600
gene_params['kTLd'][np.isnan(gene_params['kTLd'].values)] = 0

# Use protein2mRNA ratios
ratios_p2m = pd.read_csv('ratios.txt',sep='\t',header=0,squeeze=True,usecols=['Protein_mRNA']).values
ratios_p2m[101] = 0.0 #PTEN

gene_params['kTL_nat_cells'] = ratios_p2m*gene_params['kTLd']
gene_params['kTL_nat_cells'][np.isnan(gene_params['kTL_nat_cells']) | np.isinf(gene_params['kTL_nat_cells'])] = 0
gene_params['kTL_nat'] = gene_params['kTL_nat_cells']
gene_params['kTL_nat'][np.where(gene_params['kTL_nat']==0)[0]] = gene_params['kTLnatLit_s'][np.where(gene_params['kTL_nat']==0)[0]]
gene_params['kTL_nat'][np.where(gene_params['kTL_nat']==0)[0]] = gene_params['kTLnatSchwan_s'][np.where(gene_params['kTL_nat']==0)[0]]
gene_params['kTL_nat'][np.array([list(model_genes).index(x) for x in ['CCND1', 'CCND2', 'CCND3']])] = gene_params['kTL_nat'][np.array([list(model_genes).index(x) for x in ['CCND1', 'CCND2', 'CCND3']])].values * 5

xp_mpc = gene_params['kTL_nat']*mExp_mpc/gene_params['kTLd']
xp_mpc[np.isnan(xp_mpc)] = 0

ratios_kTL=pd.read_csv('ratios.txt',sep='\t',header=0,squeeze=True,usecols=['kTL_kTLd']).values
ratios_kTL[list(model_genes).index('PTEN')] = 0.0 # U87 cells are PTEN-null
xp_mpc = ratios_kTL*mExp_mpc
xp_mpc[np.isnan(xp_mpc)] = 0

# %% [markdown]
#### Read-in Initializer.txt file

The details of what to modify or keep constant are defined in the `Initializer.txt` input file. We will use different columns from that file at different stages of the `Initialization` procedure. For definitions of each column, please refer to out paper.

# %% [code]
# Read-in genes with modified mRNA levels
Step1_mrna = pd.read_csv('Initializer.txt',sep='\t',squeeze=True,usecols=['Step1_mrna','Step1_mrna_mpc'],index_col='Step1_mrna')
Step1_mrna = Step1_mrna[Step1_mrna.index.notnull()]
for gene_symbol in Step1_mrna.index:
    gene_params.loc[gene_symbol,'Exp RNA'] = Step1_mrna[gene_symbol]

# Finding reactions
reactions_all = [ratelaw_sheet[i][0] for i in range(len(ratelaw_sheet))]
vTL_pattern = re.compile("vTL\d+")
# Find translation ratelaws from the input
reactions_vTL = list(filter(vTL_pattern.match, reactions_all))
params_all = pd.read_csv('ParamsAll_U87.txt', header=0, index_col=0, sep='\t')
def params_getid(rxn,idx):
    param_id = str(params_all.index[np.logical_and(params_all['rxn'].values==rxn, params_all['idx'].values==int(idx))][0])
    return param_id

# Find the piece of stoichiometric matrix with translation ratelaws only
numberofgenes = len(model_genes)
S_PARCDL = pd.read_csv('StoicMat.txt', header=0, index_col=0, sep='\t')
S_TL = S_PARCDL.loc[:,S_PARCDL.columns.isin(reactions_vTL)]

# Read-in observables of the model - from the createModel notebook output
ObsMat = pd.read_csv('Observables.txt', header=0, index_col=0, sep='\t')
NumObs = len(ObsMat.columns)

kTL_id = []
kTL_default = []
k50E_id = []
k50E_values = []
kTL_genes = []
for rowNum, ratelaw in enumerate(ratelaw_data):  
    if "kTL" in str(ratelaw[1]):
        kTL_i = "k"+str(rowNum+1)+"_1"
        kTL_id.append(kTL_i)
        kTL_gene = re.search(r'm_\w+', ratelaw[1]).group()[2:]
        kTL_genes.append(kTL_gene)
        params_i = np.array(list(ratelaw[2:]))
        params_i = np.array([float(params_i[k]) for k in range(len(params_i))])
        params_i = params_i[~np.isnan(params_i)]
        kTL_value = params_i[0]
        kTL_default.append(float(kTL_value))
        a = len(params_i)
        if "EIF4E" in str(ratelaw[1]):
            if 'MDM2pro' in str(ratelaw[1]):
                k50E_id_i = "k"+str(rowNum+1)+"_3"
            elif 'CCND1' in str(ratelaw[1]) or 'CCND2' in str(ratelaw[1]) or 'CCND3' in str(ratelaw[1]):
                k50E_id_i = "k"+str(rowNum+1)+"_5"
            else:
                k50E_id_i = "k"+str(rowNum+1)+"_2"
            k50E_id.append(k50E_id_i)
            k50E_value_i = params_i[-1]
            k50E_values.append(k50E_value_i)   
    
# Define/read-in observables to exclude from fitting 
obs2exclude = pd.read_csv('Initializer.txt', sep='\t', usecols=['Step1_obs_excl'])
obs2exclude = obs2exclude.values[obs2exclude.notnull()]
np.append(obs2exclude,'PTEN')

kTLest = gene_params['kTL_nat'].values.copy()
kTLest[list(model_genes).index('PTEN')] = 0.0 #PTEN

mrna_id = []
mrna_filter = filter(lambda a: a.startswith('m_'), list(ICf.index))
for m in mrna_filter:
    mrna_id.append(m)

species_sheet = np.array([np.array(line.strip().split("\t")) for line in open('Species_U87.txt', encoding='latin-1')])
compartment_sheet = np.array([np.array(line.strip().split("\t")) for line in open('Compartments.txt')])
species_names = np.array([species_sheet[i][0] for i in range(1,len(species_sheet))])
# Volumes array of the species in the model
VxPARCDL = np.array([species_sheet[i][1] for i in range(1,len(species_sheet))])
VxPARCDL = [float(compartment_sheet[compartment_sheet[:,0]==VxPARCDL[i],1][0]) for i in range(len(VxPARCDL))]
VxPARCDL = pd.Series(VxPARCDL, index=species_names)

VxTL = np.ones(numberofgenes)*Vc
for i in range(np.shape(S_TL)[1]):
    if len(np.nonzero(S_TL.values[:,i])[0]) != 0:
        obs_ind = int(np.nonzero(S_TL.values[:,i])[0])
        VxTL[i] = VxPARCDL[obs_ind]
        
# protein levels in nanomolar 
pExp_nM = xp_mpc*1e9/(VxTL*6.023e23)
# model species concentrations
x0PARCDL = np.matmul(S_TL.values,pExp_nM)
x0PARCDL = pd.Series(x0PARCDL)
x0PARCDL.index = S_TL.index

mpc2nmcf_Vc=1E9/(Vc*6.023E+23)
mExp_nM=mExp_mpc*mpc2nmcf_Vc
mExp_nM = pd.Series(data=mExp_nM.values, index=mrna_id)
for m in mrna_id:
    x0PARCDL[m] = mExp_nM[m]

# Read-in species-species concentration pairs 
Step1sp = pd.read_csv('Initializer.txt', sep='\t',squeeze=True,usecols=['Step1_sp','Step1_val'],index_col='Step1_sp')
Step1sp = Step1sp[Step1sp.index.notnull()]
for sp in Step1sp.index:
    x0PARCDL[sp] = Step1sp[sp]

k50E_default = max(k50E_values)
x0 = x0PARCDL

# %% [markdown]
#### Define functions to handle model observables and to find corresponding species indices

# %% [code]
def get_observables(xout, VxPARCDL, Vc):    
    Obs = []  
    Vr = VxPARCDL/Vc
    for i in range(np.shape(ObsMat)[1]):
        Obs_i = np.sum(ObsMat.values[:,i]*xout*Vr.flatten())
        Obs.append(Obs_i)
    Obs = [0 if i <= 1e-6 else i for i in Obs]
    Obs = np.array(Obs)    
    return Obs
obs0 = get_observables(x0.values, VxPARCDL.values, Vc)
kTL_mod = np.ones(numberofgenes)*0.25
def obs2gene (obs_name):
    gene = model_genes[np.nonzero(np.matmul(ObsMat.values[:,list(ObsMat.columns).index(obs_name)],S_TL.values))[0]]
    return gene
def obs2gene_i (obs_name):
    gene_i = np.nonzero(np.matmul(ObsMat.values[:,list(ObsMat.columns).index(obs_name)],S_TL.values))[0]
    return gene_i
for m in obs2exclude:
    kTL_mod[obs2gene_i(m)] = 1.0
    kTLest[obs2gene_i(m)]
    for k in range(len(obs2gene_i(m))):
        kTLest[obs2gene_i(m)[k]] = model.getFixedParameterById(np.array(kTL_id)[obs2gene_i(m)][k])

# %% [markdown]
#### Read-in parameters with modified values

# %% [code]
Step1_par = pd.read_csv('Initializer.txt',sep='\t',squeeze=True,usecols=['Step1_par_rxn','Step1_par_idx','Step1_par_val'])
Step1_par = Step1_par[Step1_par.Step1_par_rxn.notnull()]
Step1_par_ids = [str(params_all.index[np.logical_and(params_all['rxn'].values==Step1_par.Step1_par_rxn[i], params_all['idx'].values==int(Step1_par.Step1_par_idx[i]))][0]) for i in range(len(Step1_par.Step1_par_rxn))]
[model.setFixedParameterById(Step1_par_ids[i],Step1_par.Step1_par_val[i]) for i in range(len(Step1_par_ids))]
[model.setFixedParameterById(k50E_id[k],0) for k in range(len(k50E_id))]

# %% [markdown]
#### Cyclin D
All three Cyclin D gene isoforms produce the same Cyclin D (Cd) protein species. Cyclin D controls the first entry point
to the cell cycle and its value is controled during Initialization procedure.

# %% [code]
Cd_genes = np.array([list(model_genes).index(x) for x in ['CCND1', 'CCND2', 'CCND3']])
Step1_Cd = pd.read_csv('Initializer.txt',sep='\t',squeeze=True,usecols=['Step1_Cd_par','Step1_Cd_val'],index_col='Step1_Cd_par')
Step1_Cd = Step1_Cd[Step1_Cd.index.notnull()]
kC173 = Step1_Cd['kC173']
kTL10_12 = kC173/sum(mExp_nM[Cd_genes])
[model.setFixedParameterById(Cd_kTL,kTL10_12) for Cd_kTL in np.array(kTL_id)[Cd_genes]]
kTLest[Cd_genes] = kTL10_12
kTL_initial = kTLest

# %% [markdown]
#### Model instance
Set model initial conditions

# %% [code]
# Set simulation time in seconds
ts = 1000*3600 
model.setTimepoints(np.linspace(0,ts,1000))

# set species initial conditions
x0 = x0PARCDL
model.setInitialStates(x0.values)

# model solver instance
solver = model.getSolver()
solver.setMaxSteps = 1e10

# %% [markdown]
### The main function `kTLadjustwhile`

This function is used multiple times during the Initialization procedure. It adjusts translation rate constnants and makes sure to reach steady-state values.

# %% [code]
def kTLadjustwhile(model,solver,x0, obs0, kTL_id, kTLest, kTL_mod, k50E_id, k50E_values, ObsMat, S_TL, flagE):
    if flagE == 0:
        [model.setFixedParameterById(k50E_id[k],0) for k in range(len(k50E_id))]
    elif flagE == 1:
        [model.setFixedParameterById(k50E_id[k],k50E_values[k]) for k in range(len(k50E_id))]
    model.setInitialStates(x0.values)
    [model.setFixedParameterById(kTL_id[k],kTLest[k]) for k in range(len(kTL_id))]
    m = len(ObsMat.columns)
    margin = 0.001
    # Repeat until convergence
    while m!=0:
        rdata = amici.runAmiciSimulation(model,solver)
        obs1 = rdata['y'][-1]
        error_fe = (obs0 - obs1)/obs0
        kTLf_obs = np.ones(len(obs0))
        for i in range(len(error_fe)):
            if ObsMat.columns[i] in obs2exclude:
                kTLf_obs[i] = 1
            elif obs0[i] == 0:
                kTLf_obs[i] = 0        
            elif error_fe[i] > margin and ~np.isinf(error_fe[i]):
                kTLf_obs[i] = 1/(1-error_fe[i])
            elif error_fe[i] < -1 * margin and ~np.isinf(error_fe[i]):
                kTLf_obs[i] = 1/(1-error_fe[i])
            elif error_fe[i] > -1 * margin and error_fe[i] < margin:
                kTLf_obs[i] = 1    
        kTLf = []
        for i in range(len(S_TL.columns)):
            a = np.nonzero(np.array(S_TL.iloc[:,i]))[0]
            if len(a) != 0:
                sp_ind = a[0]
                obs_ind = np.nonzero(np.array(ObsMat.iloc[sp_ind,:]))[0][0]
                kTLf.append(kTLf_obs[obs_ind])
            else:
                kTLf.append(1)
        kTLf = pd.Series(kTLf)
        kTLf = kTLf.transform(lambda x: 1 if np.isinf(x) or np.isnan(x) else x)
        kTLf = np.array(kTLf)
        kTLest = kTLest*(1+(kTLf-1)*kTL_mod)
        [model.setFixedParameterById(kTL_id[k],kTLest[k]) for k in range(len(kTL_id))]
        obs_notmatched = ObsMat.columns[~((error_fe > -1*margin) & (error_fe < margin) | (obs0==0))]
        obs_notmatched = obs_notmatched[~np.isin(obs_notmatched,obs2exclude)]
        m = len(obs_notmatched)
    kTLnew = kTLest
    [model.setFixedParameterById(kTL_id[k], kTLnew[k]) for k in range(len(kTL_id))]
    rdata_new = amici.runAmiciSimulation(model,solver)
    x_new = rdata_new['x']
    x1 = pd.Series(data=rdata_new['x'][-1], index=ObsMat.index)
    x1[x1.values<1e-6] = 0.0
    flagA = 0
    apop_def = x0['PARP']*.5 # Apoptosis event definition: If [cPARP]>[PARP], the cell dies!
    parp_all = x_new[:,list(ObsMat.index).index('PARP')]
    if sum(parp_all < apop_def) != 0:
        flagA = 1    
    return kTLnew, rdata_new, x1, flagA

# %% [markdown]
##### Below are some manual and detailed alterations. They are done to replicate Bouhaddou2018 model results. May not be necessary for other, new cell lines.

# %% [code]
EIF4Efree = float(x0['EIF4E'])
rhs = (100+EIF4Efree)/EIF4Efree
kTLnat = gene_params['kTL_nat'].values.copy()
kTL1_1 = (0.25/mExp_nM[list(model_genes).index('TP53')])*(100+EIF4Efree)/EIF4Efree
kTL2_1 = 0.2*1000/3600/mExp_nM[list(model_genes).index('MDM2')]*(100+EIF4Efree)/EIF4Efree
kTL2_2 = 2.5e-4/mExp_nM[list(model_genes).index('MDM2')]*(100+EIF4Efree)/EIF4Efree
kTL3_1 = 6.944e-5/mExp_nM[list(model_genes).index('PPM1D')]*(100+EIF4Efree)/EIF4Efree
rhs_genes = ['ATM','ATR','CDKN2A','MDM4','MSH6','BRCA2','MGMT']
for gene in rhs_genes:
    i = list(model_genes).index(gene)
    kTLest[i] = kTLnat[i]*rhs
kTLest[list(model_genes).index('CDKN2A')] = kTLest[list(model_genes).index('CDKN2A')]/3
kTLest[list(model_genes).index('MDM4')] = kTLest[list(model_genes).index('MDM4')]/3
kTLest[list(model_genes).index('TP53')] = kTL1_1
kTLest[list(model_genes).index('MDM2')] = kTL2_1
kTLest[list(model_genes).index('PPM1D')] = kTL3_1
model.setFixedParameterById(params_getid('vTL'+str(list(model_genes).index('MDM2')+1),1),kTL2_2)
kTL10_12_2 = 0.005/3600/sum(mExp_nM[Cd_genes])*rhs*17
for i in Cd_genes:
    model.setFixedParameterById(params_getid('vTL'+str(i+1),1), model.getFixedParameterById(params_getid('vTL'+str(i+1),1))*4.4)
kC104 = model.getFixedParameterById(params_getid('vC104',0))
kC104 = kC104*17
model.setFixedParameterById(params_getid('vC104',0),kC104)

# %% [markdown]
### Setting degredation rate constants

# %% [code]
kTLd = gene_params['kTLd']
kTLCd = np.zeros(len(ObsMat.columns))
for i,obs in enumerate(ObsMat.columns):
    kTLCd[i] = sum(kTLd[obs2gene(obs).values]*xp_mpc[obs2gene(obs)].values/sum(xp_mpc[obs2gene(obs)].values))
kTLCd[np.isnan(kTLCd)] = 0

# Read-in and exclude
kTLCd2exclude = pd.read_csv('Initializer.txt',sep='\t',squeeze=True,usecols=['Step1_sp_kTLCd'])
kTLCd2exclude = kTLCd2exclude.values[pd.notnull(kTLCd2exclude.values)]
for k in kTLCd2exclude:
    kTLCd[list(ObsMat.columns).index(k)] = 0

# Find degradation ratelaws
vXd_pattern = re.compile("v\D+d\d+")
reactions_vXd = list(filter(vXd_pattern.match, reactions_all))
reactions_vTLCd = list(filter(lambda x: ('vTLCd' in x),reactions_vXd))
reactions_vXd = list(filter(lambda x: ('vTLCd' not in x),reactions_vXd))

# Find the piece of stoichiometric with degradation reactions
S_Xd = S_PARCDL.loc[:,S_PARCDL.columns.isin(reactions_vXd)]

# Degradation constants array
kXd = np.zeros(np.shape(S_Xd)[1])
for kXdi in range(len(kXd)):
    ProtInd = np.nonzero(S_Xd.iloc[:,kXdi].values==-1)[0][0]
    Obs2Include = np.nonzero(ObsMat.iloc[ProtInd,:].values)[0]
    if len(Obs2Include) != 0:
        kXd[kXdi] = max(kTLCd[Obs2Include])
kXd = pd.Series(data=kXd,index=reactions_vXd)

# Manually setting some of the parameters as done in the Bouhaddou2018 model
kXdmod_1000 = ['Ractive']
for sp in kXdmod_1000:
    p = S_Xd.columns[np.argwhere(S_Xd.loc[sp].values==-1)[0][0]]
    kXd[p] = kXd[p]*1000

kXdmod_100 = ['C8','C3','C6','tBid','Baxactive']
for sp in kXdmod_100:
    p = S_Xd.columns[np.argwhere(S_Xd.loc[sp].values==-1)[0][0]]
    kXd[p] = kXd[p]*100

kXdmod_10 = ['pBIM','pBAD','pFOXO']
for sp in kXdmod_10:
    p = S_Xd.columns[np.argwhere(S_Xd.loc[sp].values==-1)[0][0]]
    kXd[p] = kXd[p]*10

# Read-in and exclude
kXd2exclude = pd.read_csv('Initializer.txt',sep='\t',squeeze=True,usecols=['Step1_sp_kXd'])
kXd2exclude = kXd2exclude.values[pd.notnull(kXd2exclude.values)]
kXd2remove = []
for k in kXd2exclude:
    kXd2remove.append(S_Xd.columns[np.argwhere(S_Xd.loc[k].values == -1)[0][0]])   
kXd = kXd[~np.isin(kXd.index,kXd2remove)]

# Set the above parameter values in the model instance
for i in range(len(kXd)):
    model.setFixedParameterById(params_getid(reactions_vXd[i],0),kXd[i])
for i in range(len(kTLCd)):
    model.setFixedParameterById(params_getid(reactions_vTLCd[i],0), kTLCd[i])

# %% [markdown]
## 2) Start Initialization Steps

# %% [markdown]
#### Fixed EIF4E, no ribosome

# %% [code]
kTLnew1, rdata_new, x1, flagA = kTLadjustwhile(model,solver,x0, obs0, kTL_id, kTLest, kTL_mod, k50E_id, k50E_values, ObsMat, S_TL, 0)

# %% [markdown]
#### Dynamic EIF4E, no ribosome

# %% [code]
kTLnew2, rdata_new, x2, flagA = kTLadjustwhile(model,solver,x1, obs0, kTL_id, kTLnew1, kTL_mod, k50E_id, k50E_values, ObsMat, S_TL, 1)

# %% [markdown]
#### Dynamic EIF4E, with ribosome back

# %% [code]
Step3_par = pd.read_csv('Initializer.txt',sep='\t',squeeze=True,usecols=['Step3_par_rxn','Step3_par_idx','Step3_par_val'])
Step3_par = Step3_par[Step3_par.Step3_par_rxn.notnull()]
Step3_par_ids = [params_getid(Step3_par.Step3_par_rxn[i],Step3_par.Step3_par_idx[i]) for i in range(len(Step3_par.Step3_par_rxn))]

kbRi = float(Step3_par.Step3_par_val[np.logical_and(Step3_par.Step3_par_rxn=='vbR',Step3_par.Step3_par_idx==int(1))])
kdR0 = float(Step3_par.Step3_par_val[np.logical_and(Step3_par.Step3_par_rxn=='vdR',Step3_par.Step3_par_idx==int(0))])
nR = float(Step3_par.Step3_par_val[np.logical_and(Step3_par.Step3_par_rxn=='vbR',Step3_par.Step3_par_idx==int(2))])
k50R = float(Step3_par.Step3_par_val[np.logical_and(Step3_par.Step3_par_rxn=='vbR',Step3_par.Step3_par_idx==int(3))])
     
ps6ki = x0['pS6K']
f1 = (ps6ki**nR)/(k50R**nR+ps6ki**nR)
Rt = x0['Ribosome']
kbR0 = Rt*kdR0 - kbRi*f1
Step3_par.Step3_par_val[(Step3_par['Step3_par_rxn']=='vbR')&(Step3_par['Step3_par_idx']==0)] = kbR0

[model.setFixedParameterById(Step3_par_ids[i],Step3_par.Step3_par_val[i]) for i in range(len(Step3_par_ids))]
kTLnew3, rdata_new, x3, flagA = kTLadjustwhile(model,solver,x2, obs0, kTL_id, kTLnew2, kTL_mod, k50E_id, k50E_values, ObsMat, S_TL, 1)

# %% [markdown]
#### Adjust Cyclin D (Cd) and p21 levels

# %% [code]
totalcyclinDfromdata = sum(pExp_nM[np.array([list(model_genes).index(x) for x in ['CCND1', 'CCND2', 'CCND3']])])
totalp21fromdata = pExp_nM[list(model_genes).index('CDKN1A')]
kC82_id = params_all.index[np.logical_and(params_all.rxn=='vC104',params_all.idx==0)][0]
kC82 = pd.read_csv('Initializer.txt',sep='\t',usecols=['Step4_par','Step4_par_val'],index_col='Step4_par', squeeze=True)['kC82']
x_in = rdata_new['x'][-1]
th=0.001
ratio_cd=0.5
ratio_p21 = 0.5
model.setInitialStates(x_in)
model.setFixedParameterById(kC82_id,kC82)
cd_sp = np.argwhere(ObsMat.loc[:,'Cd'].values>0).flatten()
p21_sp = np.argwhere(ObsMat.loc[:,'p21'].values>0).flatten()
while (ratio_cd < (1-th) or ratio_cd > (1+th)) or (ratio_p21 < (1-th) or ratio_p21 > (1+th)):
    rdata_loop = amici.runAmiciSimulation(model,solver)
    ratio_cd = totalcyclinDfromdata/sum(rdata_loop['x'][-1][cd_sp])
    if ratio_cd < (1-th) or ratio_cd > (1+th):        
        f = 1 + (ratio_cd-1)*0.125
        kC173 = kC173*f
        kTL10_12 = kC173*17/sum(mExp_nM[Cd_genes])
        [model.setFixedParameterById(Cd_kTL,kTL10_12) for Cd_kTL in np.array(kTL_id)[Cd_genes]]
    ratio_p21 = totalp21fromdata/sum(rdata_loop['x'][-1][p21_sp])
    if ratio_p21 < (1-th) or ratio_p21 > (1+th):  
        p = 1 + (ratio_p21-1)*0.5
        kC82 = kC82/p
        model.setFixedParameterById(kC82_id,kC82)
x4 = pd.Series(data=rdata_loop['x'][-1], index=ObsMat.index)
x4[x4.values<1e-6] = 0.0
kTLnew3[Cd_genes] = kTL10_12

# %% [markdown]
#### Adjust Caspase 8 (C8) basal cleavage rate

# %% [code]
ts = 1000*3600*0.5
model.setTimepoints(np.linspace(0,ts,1000))
kA77_id = params_all.index[np.logical_and(params_all['rxn']=='vA77',params_all['idx']==0)][0]
kA87_id = params_all.index[np.logical_and(params_all['rxn']=='vA87',params_all['idx']==0)][0]
kA77 = pd.read_csv('Initializer.txt', sep='\t',usecols=['Step5_kA77','Step5_kA77_val'],index_col='Step5_kA77', squeeze=True)['kA77']
model.setFixedParameterById(kA77_id, kA77)
kA87s = pd.read_csv('Initializer.txt', sep='\t',usecols=['Step5_kA87s'], squeeze=True)
kA87s = kA87s.values[~np.isnan(kA87s.values)]
for k in range(len(kA87s)):   
    model.setFixedParameterById(kA87_id, kA87s[k])
    kTLnew4, rdata_loop, x5, flagA = kTLadjustwhile(model,solver,x4, obs0, kTL_id, kTLnew3, kTL_mod, k50E_id, k50E_values, ObsMat, S_TL, 1)
    if flagA == 0:
        x5last = x5
        kTLnew4last = kTLnew4
    if flagA == 1:
        kA87 = kA87s[k-1]
        model.setFixedParameterById(kA87_id, kA87)
        break
x5 = x5last
kTLnew4 = kTLnew4last
x5[x5.values<1e-6] = 0.0
model.setInitialStates(x5.values)
[model.setFixedParameterById(kTL_id[k], kTLnew4[k]) for k in range(len(kTL_id))]

# %% [markdown]
#### Adjust basal DNA damage response

# %% [code]
BRCA2 = x5['BRCA2']
MSH6 = x5['MSH6']
MGMT = x5['MGMT']
Me = x5['Me']
Ma = x5['Ma']

fixdsb1 = model.getFixedParameterById(params_getid('vD63',0))
fixmsh = model.getFixedParameterById(params_getid('vD64',0))
fixmgmt = model.getFixedParameterById(params_getid('vD65',0))
kDDE = model.getFixedParameterById(params_getid('vD66',1))
kDEtop = model.getFixedParameterById(params_getid('vD66',3))
Etop = model.getFixedParameterById(params_getid('vD66',2))
kDnSP = model.getFixedParameterById(params_getid('vD66',4))
kDkmSP = model.getFixedParameterById(params_getid('vD66',5))
kDkmSS = model.getFixedParameterById(params_getid('vD17',2))
kDkmDS = model.getFixedParameterById(params_getid('vD14',2))

kDDbasal = 1e-6 # set this value manually
damageDSB_cycling = kDDbasal/(fixdsb1*BRCA2)
damageSSB_cycling = kDDbasal/(fixmsh*MSH6+fixmgmt*MGMT)

if damageDSB_cycling > kDkmDS:
    np.disp('ERROR --- DSB damage is too high, must reduce kDDbasal or increase strength of repair')    
if damageSSB_cycling > kDkmSS:
    np.disp('ERROR --- SSB damage is too high, must reduce kDDbasal or increase strentgh of repair')

vdamage_on = (kDDbasal + kDDE*(Etop/(Etop+kDEtop)))*(((Me+Ma)**kDnSP)/(((Me+Ma)**kDnSP)+(kDkmSP**kDnSP)))
damageDSB = vdamage_on/(fixdsb1*BRCA2)
damageSSB = vdamage_on/(fixmsh*MSH6+fixmgmt*MGMT)
kDDbasal_id = params_getid('vD66',0)
model.setFixedParameterById(kDDbasal_id, kDDbasal)
x5['damageDSB'] = damageDSB
x5['damageSSB'] = damageSSB
rdata_new = amici.runAmiciSimulation(model,solver)
x6 = rdata_new['x'][-1]
x6[x6<1e-6] = 0
x6 = pd.Series(data=x6, index=ObsMat.index)

# %% [markdown]
#### Calculate the leak terms

# %% [code]
genereg = pd.read_csv('GeneReg.txt', sep='\t', header=0, index_col=0)
numberofgenes = int(len(genereg.index))
numberofTARs = int(len(genereg.columns))

for gene_symbol in Step1_mrna.index:
    mExp_mpc[gene_symbol] = Step1_mrna[gene_symbol]
gExp_mpc = np.float64(gene_params.loc[:,'Exp GCN'].values)
kGin = np.float64(gene_params.loc[:,'kGin'].values)
kGac = np.float64(gene_params.loc[:,'kGac'].values)
kTCd = np.float64(gene_params.loc[:,'kTCd'].values)
xgac_mpc_D = (kGac*gExp_mpc)/(kGin+kGac)

TARs0 = (genereg.values)
tcnas = np.ones((numberofgenes, numberofTARs))
tck50as = np.zeros((numberofgenes, numberofTARs))
tcnrs = np.ones((numberofgenes, numberofTARs))
tck50rs = np.zeros((numberofgenes, numberofTARs))
for qq in range(numberofgenes):
    for ww in range(numberofTARs):
        pars = TARs0[qq,ww].find(';')
        if pars>0:
            nH = np.float(TARs0[qq,ww][0:pars])
            kH = np.float(TARs0[qq,ww][pars+2::])
            if nH>0:
                tcnas[qq,ww] = nH
                tck50as[qq,ww] = kH
            else:
                tcnrs[qq,ww] = abs(nH)
                tck50rs[qq,ww] = kH

mpc2nmcf_Vn = 1.0E9/(Vn*6.023E+23)
# Convert to molecules per cell
tck50as = tck50as*(1/mpc2nmcf_Vn)
tck50rs = tck50rs*(1/mpc2nmcf_Vn)

spnames = [ele for ele in model.getStateIds()]
spIDs = []
for qq in range(numberofTARs):
    sps = spnames.index(genereg.columns[qq]) 
    spIDs.append(sps)
TARarr = np.array(x6[spIDs])
TAs = np.zeros((numberofgenes,numberofTARs))
TRs = np.zeros((numberofgenes,numberofTARs))
for qq in range(numberofTARs):
    TAs[tck50as[:,qq] > 0, qq] = TARarr[qq]
    TRs[tck50rs[:,qq] > 0, qq] = TARarr[qq]
TAs = TAs*(1.0/mpc2nmcf_Vn) # convert to mpc from nM
TAs.flatten()
TRs = TRs*(1.0/mpc2nmcf_Vn)
TRs.flatten() 

# make hills
aa = np.divide(TAs,tck50as)
TFa = np.power(aa,tcnas)
TFa[np.isnan(TFa)] = 0.0
bb = np.divide(TRs,tck50rs)
TFr = np.power(bb,tcnrs)
TFr[np.isnan(TFr)] = 0.0
hills = np.sum(TFa,axis=1)/(1 + np.sum(TFa,axis=1) + np.sum(TFr,axis=1))
# With AP1*cMYC exception:
hills[Cd_genes] = np.multiply((TFa[Cd_genes,0]/(1+TFa[Cd_genes,0])),(TFa[Cd_genes,1]/(1+TFa[Cd_genes,1])))

# vTCd
vTCd= np.transpose(np.multiply(kTCd,mExp_mpc));TFa[Cd_genes,1]
vTCd = np.squeeze(np.asarray(vTCd))

kTCmax = 0.1
kTCmaxs = np.ones(len(model_genes))*kTCmax
kTCmaxs = kTCmaxs*mExp_mpc.values.astype('bool').astype('int')

induced = np.multiply(np.multiply(xgac_mpc_D,kTCmaxs),hills)
induced = induced.flatten()

negativecheck = np.array(mExp_mpc,dtype=bool).astype(int)*(vTCd - induced)
i2c = np.nonzero(negativecheck<0)[0]
if len(i2c)!=0:
    np.disp('WARNING -- Some induction term exceed degradation terms')
leak = vTCd-induced
kTCleak_new=leak/xgac_mpc_D
kTCleak_new[np.isnan(kTCleak_new)] = 0
kTCleak_new[np.isinf(kTCleak_new)] = 0

# %% [markdown]
## 3) Save the adjusted species concentration and parameter values into NEW files

We will save the newly calculated values into files with `"FileName"+"i".txt` format. Users can change this depending on their preference. 

# %% [code]
for i in range(np.shape(params_all)[0]):
    params_all.loc[params_all.index[i],'value'] = model.getFixedParameterById(str(params_all.index[i]))
params_all.to_csv('ParamsAll_U87i.txt',sep='\t')
x6.to_csv('Species_U87i.txt',sep='\t',index=True, header=False)

# %% [markdown]
## 4) Create a new SBML file and compile the Initialized model

# %% [markdown]
#### Create the new SBML file

# %% [code]
sbml_model_u87i = sbml_doc.getModel()
for sp in x6.index:
    sbml_model_u87i.getSpecies(sp).setInitialConcentration(float(x6[sp]))   
for par in params_all.index:
    sbml_model_u87i.getParameter(par).setValue(float(model.getFixedParameterById(par)))
sbml_doc_u87i = sbml_model_u87i.getSBMLDocument()
writer = libsbml.SBMLWriter()
writer.writeSBML(sbml_doc_u87i, 'SPARCED_U87i.xml')

# %% [markdown]
#### Define observables and compile using `AMICI` package

# %% [code]
formula_obs = []
for obs in ObsMat.columns:
    sp_obs = ObsMat.index[np.nonzero(np.array(ObsMat.loc[:,obs]>0))[0]]
    sp_obs_id = np.nonzero(np.array(ObsMat.loc[:,obs]>0))[0]
    Vr = VxPARCDL/Vc
    Vf = Vr*ObsMat.loc[:,obs].values
    if len(sp_obs) == 1:
        formula_i = sp_obs[0]+'*'+str(Vf[sp_obs_id][0])
    elif len(sp_obs) == 2:
        formula_i = str(sp_obs[0]+'*'+str(Vf[sp_obs_id][0])+'+'+sp_obs[1]+'*'+str(Vf[sp_obs_id][1]))
    elif len(sp_obs) > 2:
        formula_i = ''
        for j in range(len(sp_obs)-1):
            formula_i = formula_i+sp_obs[j]+'*'+str(Vf[sp_obs_id][j])+'+'
        formula_i = formula_i+str(sp_obs[-1])+'*'+str(Vf[sp_obs_id][-1])
    formula_obs.append(formula_i)
observables = {}
obs_names = list(ObsMat.columns)
for i in range(len(obs_names)):
    observables[obs_names[i]] = {}
    observables[obs_names[i]]['formula'] = formula_obs[i]

# Compile the initialized model
sbml_importer = amici.SbmlImporter('SPARCED_U87i.xml')
model_output_dir_u87 = str(model_output_dir+'_U87i')
model_path = os.path.join(model_output_dir_u87)
constantParameters = np.array(params_all.index)
sbml_importer.sbml2amici('SPARCED_U87i',
                         model_path,
                         verbose=False,
                         observables=observables,
                         constantParameters=constantParameters)

# %% [markdown]
#### Finally, check if the model is successfully compiled and ready to run.

# %% [code]
# Import the model and simulate
sys.path.insert(0, model_path)
model_module = importlib.import_module('SPARCED_U87i')
model = model_module.getModel()
solver = model.getSolver()
solver.setMaxSteps = 1e10
model.setTimepoints(np.linspace(0,10)) # set timepoints
rdata = amici.runAmiciSimulation(model,solver)
```
